# Genetic mechanisms underlying the structural elaboration and dissemination of viral internal ribosomal entry sites

**DOI:** 10.1101/2024.04.17.590008

**Authors:** Irina S. Abaeva, Tatyana V. Pestova, Christopher U. T. Hellen

**Author notes:** Corresponding author: Christopher U. T. Hellen –.

## Abstract

Viral internal ribosomal entry sites (IRESs) form several classes that use distinct mechanisms to mediate end-independent initiation of translation. The origin of viral IRESs is a longstanding question. The simplest IRESs comprise tandem pseudoknots and occur in the intergenic region (IGR) of *Dicistroviridae* genomes (order *Picornavirales*). Larger IGR IRESs contain additional elements that determine specific properties such as binding to the head of the ribosoma l 40S subunit. Metagenomic analyses reported here identified novel groups of structurally distinct IGR-like IRESs. The smallest of these (∼120nt long) comprise three pseudoknots and bind directly to the ribosomal P site. Others are up to 260nt long: insertions occurred at specific loci, possibly reflecting non-templated nucleotide insertion during replication. Various groups can be arranged in order, differing by the cumulative addition of single structural elements, suggesting an accretion mechanism for the structural elaboration of IRESs. Identification of chimeric IRESs implicates recombinational exchange of domains as a second mechanism for the diversification of IRES structure. Recombination likely also accounts for the presence of IGR-like IRESs at the 5’-end of some dicistrovirus-like genomes (e.g. Hangzhou dicistrovirus 3) and in the RNA genomes of *Tombusviridae* (order *Tolivirales*), *Marnaviridae* (order *Picornavirale*s), and the ‘Ripiresk’ picorna-like clade (order *Picornavirale*s).

## Introduction

The positive-sense RNA genomes of picornaviruses, dicistroviruses and some flaviviruses contain internal ribosomal entry sites (IRESs) which are structured RNAs that promote cap- and end-independent initiation of translation. Viral IRESs are classified on the basis of sequence and structural homology, and each type functions by a distinct mechanism that involves direct binding to components of the translation apparatus. Thus type 1 IRESs (e.g. poliovirus; ∼460 nt long), type 2 IRESs (e.g. encephalomyocarditis virus (EMCV); ∼430 nt long) and type 5 IRESs (e.g. Aichivirus; ∼410 nt long) bind to eIF4G [1–3] whereas type 4 IRESs (e.g. hepatitis C virus (HCV); ∼300 nt long) bind to the 40S ribosomal subunit and eIF3 [4, 5]. Type 6 IRESs (e.g. the intergenic region (IGR) of the dicistronic cricket paralysis virus (CrPV) genome) bind directly to 40S subunits and/or to 80S ribosomes such that the adjacent ORF2 capsid protein coding sequence enters the mRNA-binding cleft [6]. These interactions enable IRESs to bypass one or more stages in the canonical initiation process, eliminating the need for one or more eukaryotic initiation factors (eIFs). Indeed, ribosomes recruited to type 6 IRESs bypass initiation completely and enter directly into the elongation phase of translation [7, 8]. IRESs thus enable viral mRNAs to evade cellular innate immune responses that target aspects of the translation process e.g. phosphorylation of eIF2 and sequestration of capped ‘non-self’ mRNAs by interferon-induced proteins with tetratricopeptide repeats (IFITs) [9].

IRES structure is essential for function, orienting conserved motifs so that they can engage productively with components of the translation apparatus. Pairwise sequence identity between members of IRES classes is commonly below 50%, reflecting the high error rate of replication of RNA viral genomes, but structural integrity is maintained by compensatory second site substitutions that restore base-pairing [10–12]. Recombination accounts for the transfer of IRESs between viral genomes [7, 13–17] and for the apparent fusion of domains from type 1 and type 2 IRESs to form type 5 IRESs [3, 18]. IRESs are thus independent entities that can function in heterologous viral contexts. However, as there is no sequence similarity between viral IRESs and elements in cellular mRNAs, it is unlikely that IRES progenitors were captured from cellular RNAs by non-homologous recombination. Maintenance of IRES structure by compensatory substitutions and the exchange of IRESs between genomes by recombination are thus established processes in IRES evolution, but neither one accounts for the appearance of structural divergence between members of each class of IRES. The origin of viral IRESs thus remains obscure.

Metagenomic studies have exponentially increased the number of known dicistrovirus genomes, and analysis of their type 6 IRESs has revealed unexpected structural diversity. Importantly, the appearance of structural elements decorating the conserved core of type 6 IRESs is associated with the gain of novel functions. The simplest type 6 IRESs (type 6c, exemplified by Halastavi arva virus (HaLV)) (Figure 1A) are 123-129 nt long and consist of tandem pseudoknots [19]. Pseudoknot (PK) I is located immediately upstream of the first codon of ORF2, and it binds to the ribosomal peptidyl (P) site of 80S ribosomes, placing the first codon of ORF2 in the aminoacyl (A) site, which is thus able to accept cognate amino-acylated tRNA to begin translation. Type 6c IRESs do not bind stably to 40S subunits alone. PKII contains conserved L1.1a and L1.1b loops that engage with the L1 stalk of the 60S subunit and enable the IRES to bind to 80S ribosomes [19]. All other currently classified subsets of type 6 IRES comprise three pseudoknots, each of which forms the core of a domain (Figures 1A, 1B, 1E, 1F, 1H). PKI mimics initiator tRNA bound to an initiation codon and establishes the reading frame for translation of ORF2 (e.g. 20, 21). The L1.1a/L1.1b loops in PKII of type 6a IRESs (e.g. CrPV (Figure 1F)) [6, 22], type 6b (e.g. Taura syndrome virus) (Figure 1H) [23], type 6d (e.g. Caledonia beadlet anemone virus 1) (Figure 1B) [12] contain conserved HalV-like motifs that bind to the L1 stalk of the 60S subunit (e.g. 19, 24, 25) and are required for recruitment of 80S ribosomes [19, 26, 27]. The L1.1a/L1.1b loops in type 6e IRESs (Figure 1E) contain unrelated motifs but are also required for ribosomal binding [28]. The major structural differences between these subsets of type 6 IRESs are that all except type 6c contain a third pseudoknot (PKIII/domain 2) embedded in PKII/domain 1 and that PKI/domain 3 in type 6b IRESs has a stem-loop (“SLIII”) protruding from helix P3.1. PKIII is highly variable: in type 6e IRESs it is a simple H-type pseudoknot, whereas in type 6d IRESs it contains an additional protruding stem-loop (SLIV) with a U-rich apical loop. PKIII in types 6a and 6b IRESs differs with respect to the length of helix P2.2 (which in type 6a IRESs is characteristically interrupted by a conserved A/A mis-pair [29]), but in both types of IRES, PKIII contains SLIV and a second protruding stem-loop (SLV) that has a conserved apical CAGCC loop. These structural differences are correlated with specific differences in the mechanism of initiation. Thus, SLIV and SLV bind to the head of the 40S subunit [24, 25, 30], are primarily responsible for the affinity of binding of type 6a and 6b IRESs to the ribosome [31, 32] and are responsible for binding of PKI of these IRESs in the ribosomal A site [19]. PKI of type 6a and type 6b IRESs must therefore be translocated to the P site by elongation factor 2 (eEF2) before translation can commence. Type 6c, 6d and 6e IRESs generally do not bind to 40S subunits alone, but they all bind directly to the P site of 80S ribosomes (rather than to the A site) and therefore do not require this initial translocation step for decoding to commence [12, 19, 28]. The SLIII hairpin protruding from PKI in type 6b IRESs is important for translation activity but not for initial ribosomal binding [23, 33, 34]. It mimics hybrid A/P-tRNA, restricts the rotation of the 40S subunit and thereby captures the IRES/80S complex in a pre-translocation conformation that likely favors recruitment of eEF2 [35]. It is becoming evident that type 6 IRESs have a common core consisting of the PKI and PKII pseudoknots and differing peripheral domains and subdomains that modulate the mechanism of IRES-mediated initiation.

**Figure 1.**
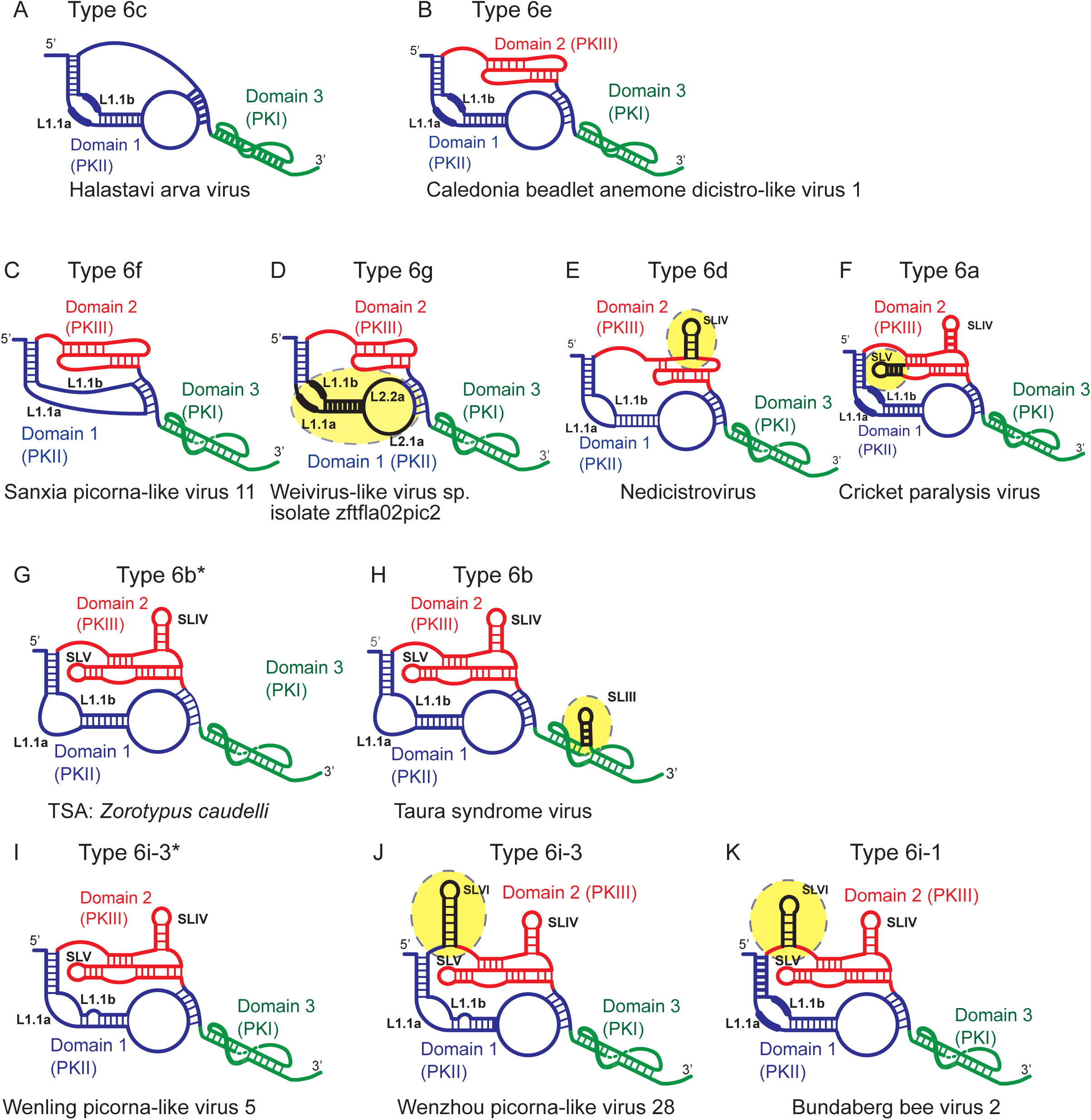
Schematic models of type 6 IRESs. Models of (a) IRES type 6c (e.g. Halastavi arva virus), (b) type 6e (Caledonia beadlet anemone dicistro-like virus 1), (c) type 6f (e.g. Sanxia picorna-like virus 11), (d) type 6g (e.g. Weivirus-like virus sp. isolate zftfla02pic2), (e) type 6d (e.g. nedicistrovirus), (f) type 6a (cricket paralysis virus), (g) type 6b* (e.g. TSA: *Zorotypus caudelli*), (h) type 6b (e.g. Taura syndrome virus), (i) type 6i-3* (e.g. Wenling picorna-like virus 5), (j) type 6i-3 (e.g. Wenzhou picorna-like virus 28) and (k) type 6i-1 (e.g. Bundaberg bee virus 2). Models are labeled to show domains 1, 2 and 3, stemloops SLIII, SLIV, SLV and SLVI, and L1.1a and L1.1b loops (with conserved motifs in these loops indicated by thick lines). IRESs are arranged and novel domains in each IRES type are colored black and highlighted with yellow shaded ovals to illustrate incremental structural additions between (c-f) IRES types 6f, 6g, 6d and 6a), (g, h) IRES types 6b* (lacking SLIII) and 6b, and (i - k IRESs types 6i-3* (which lack SLVI), 6i-3 and 6i-1.

Many dicistrovirus sequences remain uncharacterized, and we therefore extended our studies to identify and classify additional type 6 IRES structures. Here we report four distinct structural variants that we have designated types 6f, 6g, 6h and 6i. Comparison of the sequences of these and previously categorized subtypes revealed that type-specific differences correspond to insertions at a few specific loci: this observation suggests a mechanism for the structural elaboration of IRESs. We also identified examples of apparent recombination involving different classes of type 6 IRESs, implicating recombination as a second mechanism that leads to the diversification of IRESs structure. Our studies revealed type 6 IRESs located at the 5’-end of some dicistrovirus-like genomes (rather than in the intergenic region) as well as at 5’-terminal and internal positions in the single-stranded RNA genomes of viruses other than dicistroviruses. Horizontal gene transfer, previously identified as a mechanism for the dissemination of IRES types 2, 4 and 5, is thus also exploited by type 6 IRESs.

## METHODS

### Sequences

Sequences were analyzed from viruses as indicated in the text and in Supplementary tables.

### Identification of candidate type 6 IRESs

Candidate type 6 IRES sequences were identified in the NCBI database by searching nucleotide collection (nr/nt) and TSA sequences using BLASTN (http://www.ncbi.nlm.nih.gov/BLAST/) and non-redundant protein and TSA protein sequences using BLASTX. Nucleotide and polypeptide searches used the parameters: E, 1000; word size, 11; match/mismatch scores, 1/1; gap costs, 2/1; and E, 1000; word size, 6; Matrix: BLOSUM62; gap costs, 9/1, respectively. Hits were characterized by six-frame translation (http://molbiol.ru/eng/scripts/01_13.html) to identify those sequences that corresponded to genomic fragments of members of the *Picornavirales*. Sequences of potential translation products were used in BLAST searches when appropriate to verify that the C-terminal region of ORF1 encoded the 3D polymerase and that ORF2 encoded capsid proteins; Clustal Omega (http://www.ebi.ac.uk/Tools/msa/clustalo/) was used to align ORF2 and known dicistrovirus ORF2 sequences to identify potential initiation codons and, with ORF1 sequences, to estimate the borders of potential IRESs. IGR sequences were aligned using Clustal Omega and EMBOSS Matcher (http://www.ebi.ac.uk/Tools/psa/emboss_matcher/nucleotide.html) and adjusted manually to match predicted structural elements.

### Modeling of IGR IRES structures

Secondary structures were identified by free energy minimization using Mfold [36]. Tertiary structures were initially modeled using pKiss (http://bibiserv2.cebitec.uni-bielefeld.de/pkiss) [37] implemented using the Andronescu model for thermodynamic folding parameters [38], and using IPKnot (http://rtips.dna.bio.keio.ac.jp/ipknot/) [39] with default parameters. Mutational analysis was undertaken to test structural models and to resolve the structure of regions of ambiguity. The quality of models was assessed by determining their compatibility with sequence variation between viruses. To visualize the structures of pseudoknots, the pKiss base-pairing output in Stockholm format was submitted to RNAComposer [40]. The resulting 3D structure was visualized using ChimeraX [41].

### Plasmids

Transcription vectors for wt IRES-containing mRNAs that were used in *in vitro* translation and *in vitro* reconstitution experiments consisted of a T7 promotor, the sequence 5’-GGGCCCGACCCGGTGACGGGTCGGGCC-3’ (which forms a stable hairpin (ΔG = -29.10 kcal/mol)) and the viral sequence of interest with a common primer binding site (5’-CAAGGCAATCACACC-3’) replacing viral sequences starting 60-72nt downstream of the putative initiation codon, cloned between BamH1 and EcoR1 sites of pUC57 (Genewiz, South Plainfield, NJ). The viral sequences corresponded to Changjiang picorna-like virus 7 nt 5567-6223, Hubei picorna-like virus 19 nt 5242-5982, Sanxia picorna-like virus 11 nt 5331-6050, and Perth bee virus 2 nt 5676-5972 linked to Changjiang picorna-like virus 7 nt 5822-6215 (to extend the coding region beyond the current 3’-border of the current partial Perth bee virus 2 genomic sequence, which includes only ∼140nt of ORF2). These sequences were modified in the 3’-terminal region by substitutions that introduced ATG triplets to increase [^35^S]-methionine labeling of translation products.

Transcription vectors for a Sanxia 11 IGR IRES ’stop codon’ variant comprised a T7 promotor, the stable hairpin sequence and Sanxia 11 nt 5331-5635 cloned between BamH1 and EcoR1 sites of pUC57, with the 5’-CAAGGCAATCACACC-3’ primer binding site replacing viral sequences starting 72 nt downstream of the putative ACA_5485-5487_ initiation codon and a UAA stop codon replacing the putative ACA_5485-5487_ initiation codon.

The HalV IGR-STOP codon transcription vector has been described [19].

### Purification of factors and ribosomal subunits, and aminoacylation of tRNA

40S and 60S ribosomal subunits, eEF1H, eEF2 and total (Σ) aminoacyl-tRNA synthetases were purified from rabbit reticulocyte lysate (Green Hectares, Oregon, WI) [42, 43]. eRF1 and eRF3 were expressed and purified [19]. RelE [44] was a gift from V. Ramakrishnan (MRC LMB, Cambridge, UK). Native total calf liver tRNA (Promega) was aminoacylated using total native aminoacyl-tRNA synthetases [43].

### Assembly and analysis of ribosomal complexes

For toeprinting analysis, ribosomal complexes were assembled as described [7] unless otherwise stated in the text. Aliquots of 2 pmol of IGR-IRES RNA were incubated for 5 min at 37°C in 40 μl reaction volumes that contained buffer A [2 mM dithiothreitol (DTT), 100 mM potassium acetate, 20 mM Tris (pH 7.5), 2.5 mM magnesium acetate, 1 mM ATP, 0.1 mM GTP and 0.25 mM spermidine] and 6 pmol 40S subunits with or without 8 pmol 60S subunits, 20 pmol eRF3 and 15 pmol eRF1 as indicated.

To examine the elongation competence of 80S complexes, reaction mixtures were supplemented with combinations of 10 pmol eEF2, 50 pmol eEF1H, 15 μg aminoacylated native total tRNA (Σaa-tRNA) and 500 μg/ml cycloheximide, and incubation continued for another 10 min. The resulting complexes were analyzed by primer extension using avian myeloblastosis virus reverse transcriptase (Promega) and a primer (5′-GGTGTGATTGCCTTG) complementary to the common primer-binding site 61–75 nt downstream of each putative start codon that had been 3′-[^32^P]-end-labeled [42]. cDNA products were resolved in 6% polyacrylamide sequencing gels, followed by autoradiography, and were compared with appropriate dideoxynucleotide sequence ladders to map the positions of toeprints.

### Analysis of ribosomal complexes by RelE cleavage

Cleavage of ribosome-bound mRNA by RelE was analyzed as described [45]. The 80S ribosomal complexes were assembled with or without eEF2 as described above for analysis of elongation competence and then incubated with 20 pmol RelE for 10 min at 37°C. mRNA was phenol extracted and analyzed by primer extension using AMV RT and the same ^32^P-labeled primers as those used for toeprinting analysis. cDNA products were resolved in 6% polyacrylamide sequencing gels.

### Quantification and statistical analysis

All *in vitro* experiments were repeated at least three times, and they included technical and biological replicates. Representative gel images are shown.

## RESULTS

### Identification of an exceptionally compact class of IGR IRES

Analysis of the dicistronic genomes of Changjiang picorna-like virus 7 (Changjiang 7), Hubei picorna-like virus 19 (Hubei 19), Sanxia picorna-like virus 11 (Sanxia 11), Perth Bee virus 2 (Perth Bee2) and *Picornavirales* sp. isolate 77-k141_404334 [46–48] suggested that they contain related IGR IRESs (Table S1) that are located immediately downstream of the ORF1 stop codon and that are much shorter than canonical ∼190nt-long type 6a IRESs. Models of these IRESs (Figure 2A-2E) suggest that they are only 120-127 nt long (including the non-AUG initiation codon), share 58-72% pairwise nucleotide sequence identity and have a common three-domain structure. Domain 2 consists of a compact, conserved pseudoknot (PKIII) nested within domain 1 (PKII) and domain 3 consists of a single pseudoknot, PKI. In addition to having a conserved structure, the Sanxia 11-type IGR IRESs contain conserved sequence elements in and immediately downstream of PKIII, in PKI and in the L1.1a/L1.1b loops (Table S1).

**Figure 2.**
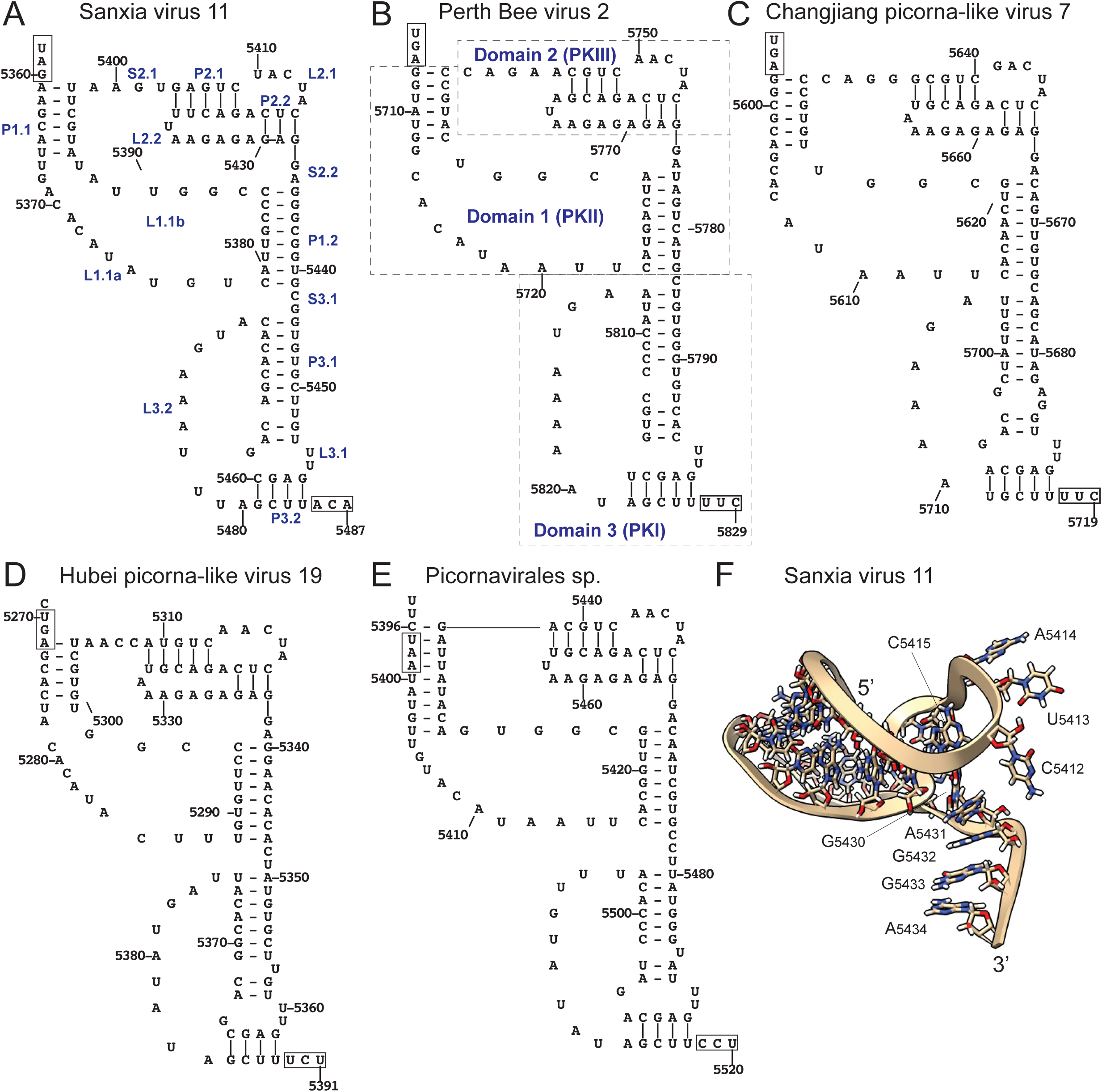
Structures of type 6f intergenic region (IGR) IRESs. Structural models of (A) Sanxia picorna-like virus 11, (B) Perth bee virus 2, (C) Changjiang picorna-like virus 7, (D) Hubei picorna-like virus 19 and (E) Picornavirales sp. isolate 77-k141_404334 IRESs, labeled to show the ORF1 stop codon and the ORF2 initiation codon. Helical segments (P), internal loops (L) and single-stranded regions (S) are labeled sequentially in panel (B) in red font according to the domain in which they occur; panel (B) shows pseudoknots (PK) I, II and III (indicated in blue font), and domains 1 and 2 are demarcated by dashed lines. Nucleotides are labeled at 10 nt intervals. (F) Tertiary structure model of domain 2 of the Sanxia picorna-like virus 11 type 6f IRES, generated using RNAcomposer and visualized using ChimeraX. The model is oriented to show solvent-exposed nucleotides in the L2.1 loop and the adjacent S2.2 single-stranded spacer.

The predicted structures of the Sanxia 11 class of IRESs differ in key respects from other type 6 IRESs: domain 1 is much smaller, primarily because it lacks a P1.2 helix, so that helices P1.1 and P1.3 are instead connected by two large loops. They are designated L1.1a and L1.1b, and they contain conserved (C/U)ACACUA and CGGU motifs, respectively (Figure 2A). PKIII in these IGR IRESs is a conserved, compact (25 -27 nt-long) H-type pseudoknot, and it contains two highly conserved motifs, GUCn**ACUA**<u>CUC</u>AGA and AAGAGA<u>GAG</u>GA (in which the underlined nucleotides are base-paired). It is much smaller than PKIII in type 6a and 6b IRESs, and it lacks equivalents of their SLIV and SLV hairpins that engage with the head of the 40S subunit [25, 49]. Modelling of the three-dimensional structure of PKIII from the Sanxia 11 IRES (Figure 2F) suggests that the conserved ACUA tetranucleotide in motif 1 is solvent-exposed. Although type 6e IRESs also contain a conserved H-type pseudoknot [12], its consensus sequence and that of Sanxia 11 class IRESs are unrelated. Type 6d IRESs [28] contain sequence motifs in PKIII that are related to those of Sanxia 11-like IRESs (see below), but the size and structure of these two types of IRES otherwise differ significantly. Taken together, the exceptionally compact nature and the distinct sequence and structural characteristics of Sanxia 11-like IGR IRESs differentiate them from other IRES classes. We therefore classified them as type 6f IRESs.

### Factor-independent initiation by direct P site ribosomal binding of type 6f IGR IRESs

The ability of type 6f IRESs to mediate internal ribosomal entry was first assayed by translation in rabbit reticulocyte lysate, using mRNAs in which the IGR and adjacent ORF2 were placed downstream of a 5’-terminal hairpin that is sufficiently stable (ΔG = -32.40 kcal/mol) to block end-dependent ribosomal attachment [50] (Figure 3A). Sanxia 11, Perth Bee 2 and Hubei 19 IGR-linked mRNAs yielded translation products of the expected sizes (Figure 3B).

**Figure 3.**
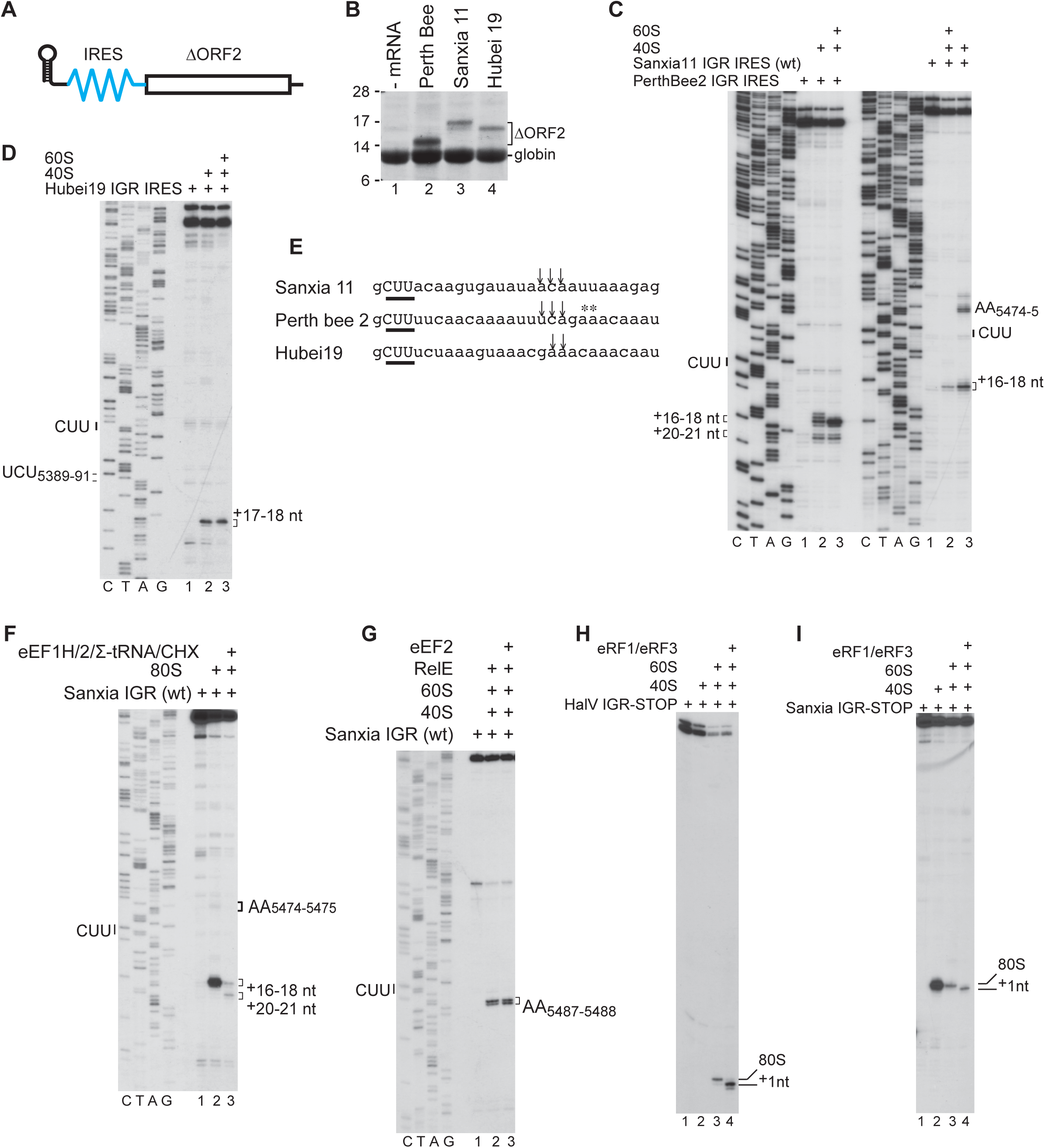
Internal initiation of translation mediated by type 6f IGR IRESs. (A) Schematic representation of a representative monocistronic mRNA containing a stable 5’-terminal hairpin followed by nucleotides corresponding to a type 6f IRES and the adjacent segment of the cognate ORF2 coding sequence. (B) The activity of Perth Bee 2, Sanxia 11 and Hubei 19 IGRs in promoting translation in RRL. The globin translation product arises from translation of endogenous mRNA in the lysate. (C - D) Binding of 40S subunits and assembly of 80S ribosomal complexes on the (C) Sanxia 11 and Perth bee 2 and (D) Hubei 19 IGR IRESs in the presence of the indicated components, assayed by toeprinting. The CUU triplet in PKI and the positions of toeprints at AA_5474-5475_ (Sanxia 11) and relative to the ^+^1 nt of the CUU triplet (all three IRESs) are indicated to the right and left of panels. (E) ORF2 sequences are shown with annotation to indicate the CUU triplet and the positions of the strongest sets of toeprints. (F) Toe-printing analysis of ribosomal movement after a single cycle of elongation on the *wt* Sanxia 11 IGR IRES in the presence of cycloheximide, elongation factors eEF1H and eEF2 and total aminoacylated tRNAs (∑ aa-tRNA). Toeprints corresponding to IRES-bound ribosomal complexes are indicated to the right. (G) Analysis of A-site accessibility in 80S ribosomes bound to the *wt* Sanxia 11 IGR IRES in the presence and absence of eEF2, assayed by mapping mRNA susceptibility to RelE cleavage. The positions of RelE cleavages are indicated to the right. (H, I) Toe-printing analysis of binding of eRF1 and eRF3 to 80S ribosomal complexes assembled on (H) HalV IGR IRES-STOP mRNA and (I) Sanxia 11 IGR IRES-STOP mRNA. The ^+^1nt forward movement of the ribosome induced by eERF1 binding is indicated. (C, D, F, G) A dideoxynucleotide sequence generated with the same primer was run in parallel on these gels (lanes C, T, A and G) to show the corresponding viral sequences.

Initiation mediated by IGR IRESs involves their factor-independent binding directly to an 80S ribosome or to a 40S subunit followed by joining of a 60S subunit [6, 7, 12, 28, 51]. To determine whether ribosomes bind directly to a specific site on type 6f IRESs, we used toeprinting, which involves inhibition of reverse transcriptase-mediated extension of a primer that has been annealed to mRNA downstream of a ribosomal complex. Ribosomes yield toeprints ∼ +15-+17 nt from the first (^+^1) nucleotide of the P-site codon on which they have assembled [1]. Sanxia 11, Hubei 19 and Perth Bee 2 IGR IRESs bound stably to 40S subunits and 80 ribosomes, yielding toeprints ^+^16-^+^18 nt relative to the CUU triplet in PKI (Figures 3C , 3D; summarized in Figure 3E). Ribosomal binding to the Perth Bee 2 IRES yielded an additional weaker set of toeprints at ^+^20-^+^21 positions (Figure 3C, lanes 2, 3). Judging from the position of the common set of toeprints at ^+^16-^+^18 positions relative to the CUU triplet in PKI, these IRESs most likely primarily bind in the P site, in which case the second set of toeprints seen with the Perth Bee 2 IRES may reflect binding of PKI in the ribosomal E site. Inclusion of 60S subunits with 40S subunits and the Sanxia 11 IRES led to the appearance of additional toeprints at AA_5474-5475_ in the L3.2 loop of PKI (Figure 3C, lanes 5, 6) which suggests that this loop may also engage in a specific interaction with the 80S ribosome.

We selected the Sanxia 11 IRES as a representative of this class for further analysis. To confirm that this IRES binds directly to the ribosomal P site, it was first incubated with 80S ribosomes, total aminoacylated tRNAs, eEF1H and eEF2 in the presence of cycloheximide, which binds in the ribosomal E site interfering with translocation and on CrPV-like IRESs inhibits elongation after PKI reaches the E site [7, 52]. This incubation yielded toeprints at +19-20nt positions (Figure 3F, lane 3), indicative of forward translocation of the ribosomal complex by one codon. By contrast, ribosomal complexes assembled on the CrPV type 6a IRES, which binds in the A site, underwent two cycles of forward translocation in these conditions [7]. These experiments therefore suggested that the CUU triplet in PKI of the Sanxia 11 IRES binds directly in the P site.

To obtain further confirmation of P site IRES binding, we employed the bacterial toxin RelE that cleaves ribosomal-bound mRNA in the A site [19, 53]. Whereas RelE cleavage at the 3’ border of the CrPV IRES occurs only after eEF2-mediated translocation of PKI from A to P sites, cleavage downstream of the HalV IRES is eEF2-independent because PKI binds in the P site and the A site is accessible to ligands [19]. RelE cleaved Sanxia 11 IRES mRNA at the same position just downstream of the PKI CCU codon in the presence and in the absence of eEF2 (Figure 3G, lanes 2, 3) indicating that the A site in IRES-bound ribosomes is vacant, and that eEF2-mediated translocation is not required for the codon in this site to be decoded.

Binding of the eukaryotic release factor eRF1 to the A site induces a ^+^1-^+^2 nt shift in the toeprint, due to compaction of the stop codon and the adjacent downstream nucleotide in the A site [54, 55]. We therefore compared binding of eRF1/eRF3 to 80S ribosomal complexes assembled on HalV and Sanxia 11 IRES variants in which the codons that immediately follow the PKI CUU codons have been substituted by stop codons. In both cases, we observed a similar ^+^1-^+^2 nt shift in the toeprint (Figures 3J-K) indicating that the A sites contain stop codons and are accessible for eRF1. In contrast, eRF1/eRF3 binding to 80S ribosomes assembled on the CrPV IRES only occurred after a cycle of eEF2-mediated translocation [19, 56]

Thus, these results are all consistent with a model in which type 6f IRESs bind to ribosomes so that PKI enters the P site and the adjacent downstream codon is accessible for decoding by aminoacyl-tRNA/eEF1A-GTP.

### Related PKIII pseudoknots in multiple classes of IGR IRES

The unusual properties of type 6f IRESs prompted us to investigate their relationship to other IGR IRESs. We have established a pipeline to identify divergent IRESs by analysis of metagenomic sequences, including metagenome-assembled genomes (MAG) and transcriptome shotgun assembly (TSA) sequences [12, 19, 28]. IGR IRESs were identified on the basis of their location in viral genomes and genomic fragments, flanked by open reading frame ORF1 that encodes nonstructural proteins (including the 3C protease and 3D polymerase) and ORF2 that encodes structural proteins that contain characteristic dicistrovirus-specific motifs [57–59]. Here we report the discovery and classification of additional groups of type 6 IRESs, several of which have conserved type 6f-related sequence motifs in PKIII but differ in that they contain an incrementally increasing number of additional structural elements.

The first group (type 6g) was identified in members of the metazoan phyla *Arthropoda* and *Mollusca*. This group consists of two subsets of IGRs that are 144 - 173 nt long, up to and including the initiation codon. Members of both subsets contain conserved type 6f-related sequence motifs in PKIII and a large central helix (P1.2) in domain 1 (Figures 4A, 4B) that is notably not present in type 6f IRESs. The subsets differ from each other in that subset 6g-1 e.g. Weivirus-like virus sp. Isolate zftfla02pic2 (Figure 4A) contains conserved AUAUACG and CAGGA motifs in the L1.1a and L1.1b loops, respectively (Table S2A) whereas subset 6g-2 (e.g. Changjiang picornalike virus 6 [46]) (Figure 4B)) contains a conserved UUUA motif in the L1.1a loop and a minimal L1.1b loop (Table S2B). In other respects, these subsets are similar: for example, they have related motifs in loop L1.2a (GUACAUA in subset 6g-1 and ACAUA in subset 6g-2) and in loop L1.2b (CGGC and CGG, in 6g-1 and 6g-2, respectively). In addition to six closely related members in subset 6g-1 and eight closely related members in subset 6g-2, there are divergent members that have the same motifs in loop L2.1 and helix P2.2 as type members, but in which other elements in domain 2 differ, resulting variously in extended L2.1 and L2.2 loops, extended P2.1 and P2.2 helices and variable S2.1 and S2.2 spacers (e.g. Figures 4C, 4D). Of note, in some type 6g IRESs (e.g. Hubei picorna-like virus 21, Sanxia picorna-like virus 13 and TSA: *Ranatra linearis*) the P2.2 helix is extended, potentially containing five to seven base-pairs (bp) flanking two opposed adenosine residues reminiscent of type 6a IRESs, and in others, the long S2.1 spacer in the *Lactuca sativa* dicistroviridae strain pt151-dic-17 IRES (subset 6g-2) has the potential to form a short stem-loop (Figures 4C, 4D).

**Figure 4.**
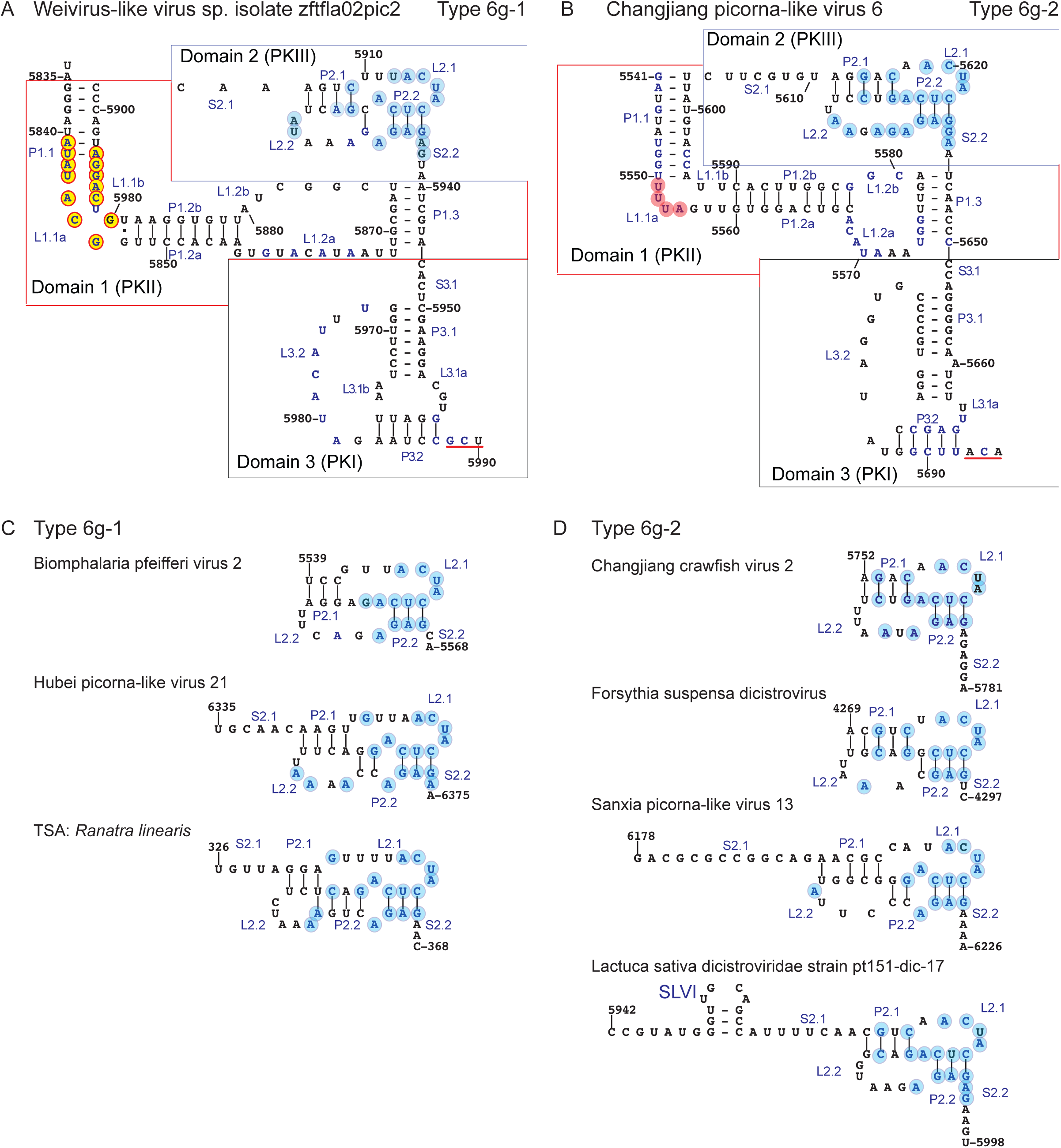
Structures of type 6g intergenic region (IGR) IRESs. (A, B) Structural models of (A) Weivirus-like virus sp. isolate zftfla02pic2 (GenBank: MT138418.1) and (B) Changjiang picorna-like virus 6 strain CJLX30470 (GenBank: NC_032794.1) IRESs. Secondary structure elements, including base-paired helical segments (P), internal loops (L) and single-stranded regions (S) are labeled sequentially in blue font in panel (A) according to the domain in which they occur. Domain 1 (pseudoknot (PK) II) , domain 2 (PKIII) and domain (3 PKI) are demarcated in panels A and B by dashed lines, and the ORF2 initiation codon is underlined. Nucleotides are labeled at 10 nt intervals. Conserved nucleotides in L1.1a and L1.1b are indicated by yellow circles in (A) and by pink circles in (D); conserved nucleotides in PKIII in (A - D) are indicated by light blue circles. (C, D) Structural models of domain 2 of the IGR IRESs of (C) *Biomphalaria pfeifferi* virus 2 (GenBank: MT108489), Hubei picorna-like virus 21 strain WHBL73090 (GenBank: NC_033069.1) and TSA: Ranatra linearis C97379_a_3_0_l_1271 (GAYZ02023980.1) (members of subset 6g-1) and (D) Changjiang crawfish virus 2 strain CJLX30764 (GenBank: KX884558.1), Forsythia suspensa dicistrovirus strain pt110-dic-2 (GenBank: MN728807.1), and Sanxia picornavirus 13 strain SXXX34014 (GenBank: NC_033266.1) and *Lactuca sativa* dicistroviridae strain pt151-dic-17 (GenBank: MN722413.1) (members of subset 6g-2).

The second group of IRESs with conserved type 6f-like motifs in PKIII was defined as type 6d [28]. Further analysis revealed numerous additional IRESs in this class (Table S3). They are 160-193 nt long, share 26-97% sequence identity, and as in type 6f and type 6g IRESs, the L1.1a and L1.1b loops contain AUAUACG and GnCAGG motifs, respectively, and PKIII contains conserved **UACUA**<u>CUC</u>AG and CA<u>GAG</u>AA motifs, in which the underlined nucleotides are base-paired to form a CUC/GAG mini-helix and nucleotides in bold form an exposed loop (Table S3). Type 6d IRESs contain a large central helix (P1.2) in domain 1, like type 6g IRESs, but they differ from them and from type 6f IRESs in additionally containing a short hairpin with a conserved apical U-rich loop nested in the L2.1 loop of PKIII (Figure 5). This hairpin is analogous to SLIV of type 6a and type 6b IRESs. The L1.1a and L1.1b loops in previously reported type 6d IRESs such as those of nedicistrovirus (NediV), Antarctic picorna-like virus 1 (APLV1) [28] and Hangzhou dicistro-like virus 4 Figure 5A) contain conserved, functionally important motifs that are different from those in other classes of IGR IRES [28]. Numerous additional members of this group (e.g. Changjiang picorna-like virus 10) contain all of the type-specific motifs in L1.1a and L1.1b loops and in PKIII, but the P2.1 and P2.3 helices in PKIII are extended relative to NediV (Figure 5B). For example, Weivirus-like virus sp. isolate cra070shi3 and *Lactuca sativa* dicistrovirus strain pt151-pil-9-plant 151 [61] contain type-specific motifs but also have extended P2.1 and P2.3 helices and an enlarged L2.3 loop that in some instances can form a short hairpin (Figures 5C, 5D).

**Figure 5.**
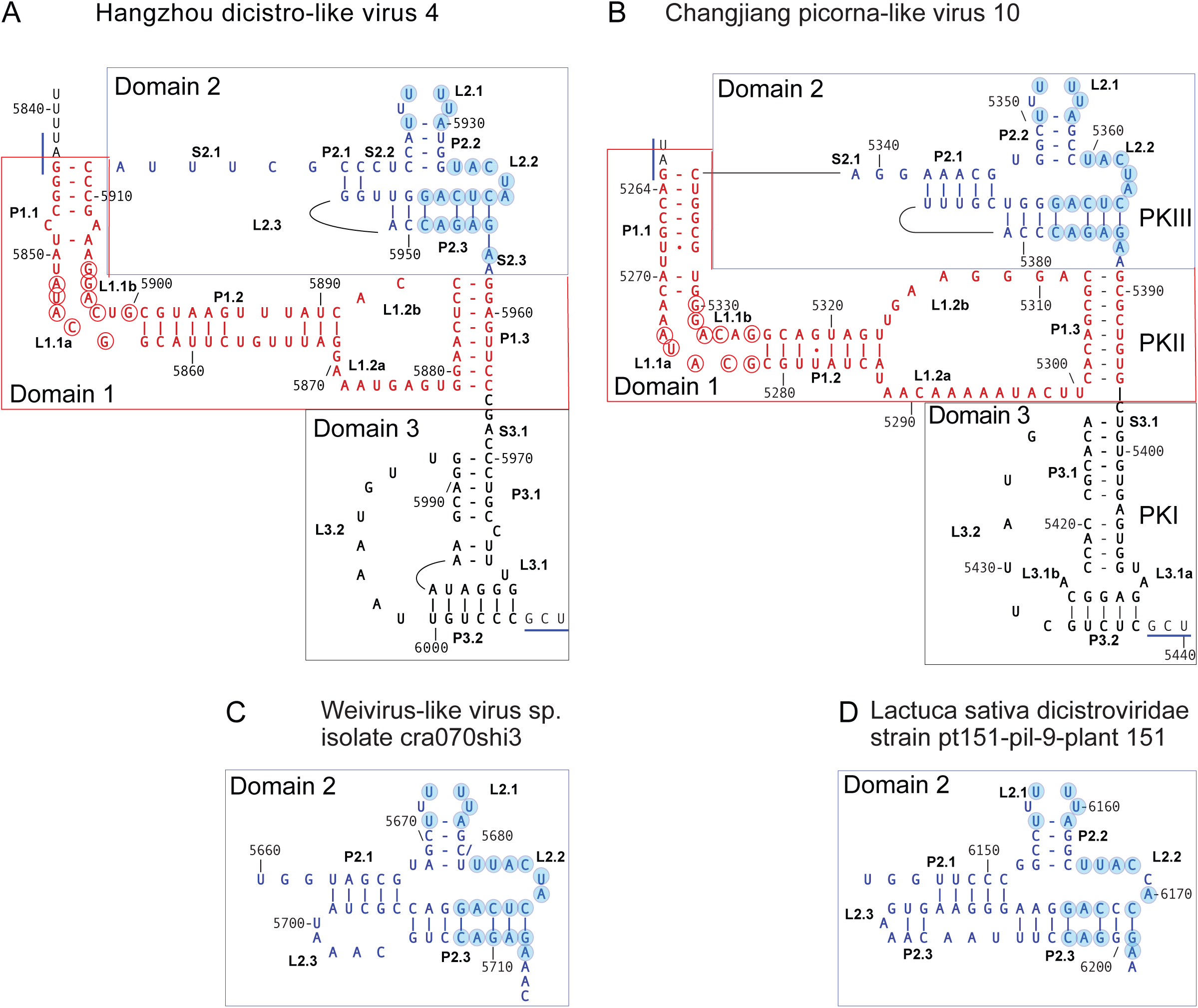
Structures of type 6d intergenic region (IGR) IRESs. (A, B) Models of the IGR IRESs of (A) Hangzhou dicistro-like virus 4 (Genbank: OQ363017), labeled to show secondary structure elements, including helices (P), internal loops (L) and single-stranded regions (S) and (B) Changjiang picorna-like virus 10 (NC_032823.1), labeled to show pseudoknots (PK) I, II and III, stem-loop (SL) IV and domains 1, 2 and 3. The ORF2 initiation codon is underlined. (C-E) Models of domain 2 of (C) Weivirus-like virus sp. isolate cra070shi3 (Genbank: MT138207.1) and (D) Lactuca sativa dicistroviridae strain pt151-pil-9-plant-151 (Genbank: MN841303.1). (A-D) Nucleotides are labeled at 10 nt intervals, and conserved nucleotides in L1.1a/L1.1b, in SLIV and in L2.2/P2.3/S2.3 are indicated by red unfilled and light blue circles, respectively.

The third group of IRESs with conserved type 6f-like motifs in PKIII constitute a subset of type 6a IRESs. They contain a central P1.2 helix in domain 1, and SLIV and SLV hairpins in domain 2 (Figure 6). The presence of SLV distinguishes these IRESs from type 6e and 6f IRESs. This subset of IRESs is very extensive, and only a representative selection of them is listed in Table S4. These IRESs contain conserved motifs in L1.1a, L1.1b, SLIV and SLV, and conserved nucleotides at other locations that are typical of canonical type 6a IRESs (e.g. [29]). Many of these previously undescribed type 6a IRESs, such as Guiyang dicistrovirus 2 (Figure 6A) have the same or very similar motifs in PKIII as the **UACUA**<u>CUC</u>AG and CA<u>GAG</u>AA motifs in the type 6e, 6f and 6g IRESs (Table S4: IRESs #1 - #15). These motifs are degenerate in other type 6a IRESs, although the substitutions are generally transition mutations, so that the length of base-pairing in helical elements is maintained. For example, the TSA: *Cyclosa laticauda* IRES has a CUC/GGG minihelix instead of CUC/GAG minihelix (indicated by a dashed box in Figure 6B; Table S4: #16) and numerous IRESs e.g. Wuhan arthropod virus 2 (Table S4: IRESs #18-#24) have a CCC/GGG minihelix at this position (Figure 6C). Despite sequence differences elsewhere in these IRESs, the structure of the exposed face of PKIII that is formed by the **UACUA**<u>CUC</u>AG and CA<u>GAG</u>AA motifs is therefore likely conserved.

**Figure 6.**
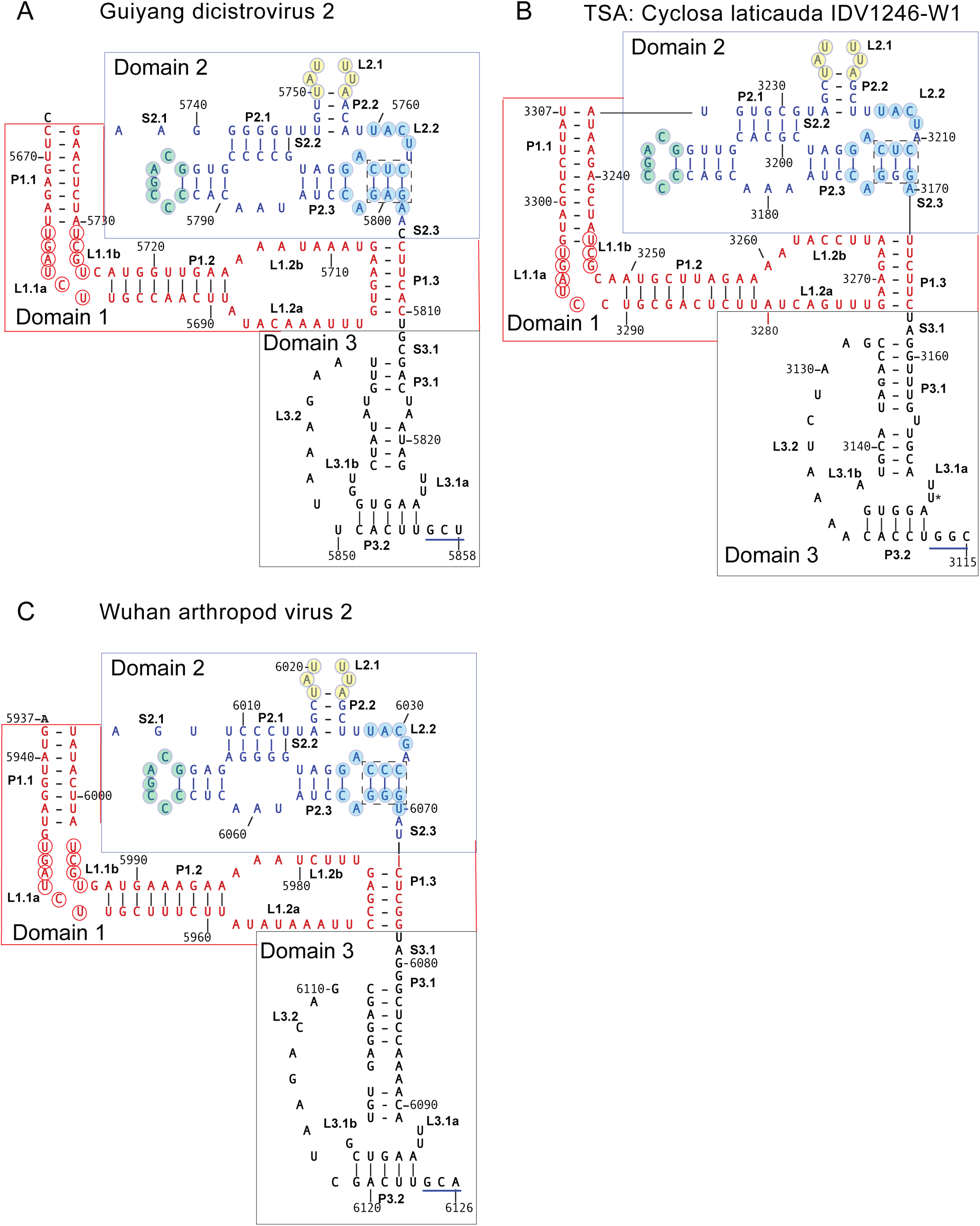
Structures of type 6a intergenic region (IGR) IRESs. (A, B) Models of the type 6a IRESs of (A) Guiyang dicistrovirus 2 (Genbank: MZ209774.1), (B) TSA: *Cyclosa laticauda* (Genbank: IAWK01021638.1) and (C) Wuhan arthropod virus 2 strain WHSF0 (Genbank: KX884282.1) (Shi et al., 2016), annotated to show the ORF2 initiation codon (underlined), secondary structure elements, including part of the helical element P2.3 (indicated by a dashed box), stem-loops SLIV and SLV, pseudoknots PKI, PKII and PKIII, and domains 1 - 3. Nucleotides are labeled at 10 nt intervals according to the Genbank sequences. Conserved nucleotides in L1.1a/L1.1b, SLIV, SLV and L2.2/P2.3/S2.3 are indicated by clear, pale yellow, grey, and light blue circles respectively.

We have thus identified four distinct but related classes of dicistrovirus IGR IRESs (types 6d, 6f, 6g and a subset of type 6a). They all have a three-domain structure and contain closely related motifs at identical locations in PKIII that likely form a conserved supporting substructure and exposed surface. However, they differ by the presence of a progressively increasing number of additional structural elements in domain 2 (helix P1.2) and domain 3 (SLIV, SLV and the extended P2.1 and P2.3 helices).

### Sequence variability in type 6b IRESs

The principal distinguishing feature of type 6b IRESs is SLIII in PKI, but there are also more subtle differences between them and type 6a IRESs. Helix P2.2 in type 6b IRESs is continuous whereas it is interrupted by an A:A mismatch in type 6a IRESs, the L2.2 loop is shorter in type 6b IRESs than in type 6a IRESs, whereas the L1.1a and L1.1b loops are larger in type 6b than in type 6a IRESs and contain UGGUUACC and UAAGGCUU motifs, respectively, rather than the UGUGAUCU and UGCUA motifs in type 6a IRESs [29, 31]. These observations were based on analysis of only five type 6b IRESs but were confirmed here by sequence alignment of a more extensive dataset (Table S5). 44 nucleotides are conserved in >90% of these sequences, and they map to the L1.1a, L1.1b and L1.2a loops in domain 1, to helix P2.1, loop L2.2, SLIV and SLV in domain 2 and to loop L3.4, helix P3.3 and nucleotides flanking SLIII in domain 3 (e.g. Figure 7E). These elements include the principal contact points for the ribosome on type 6b IRES [25, 35, 61].

**Figure 7.**
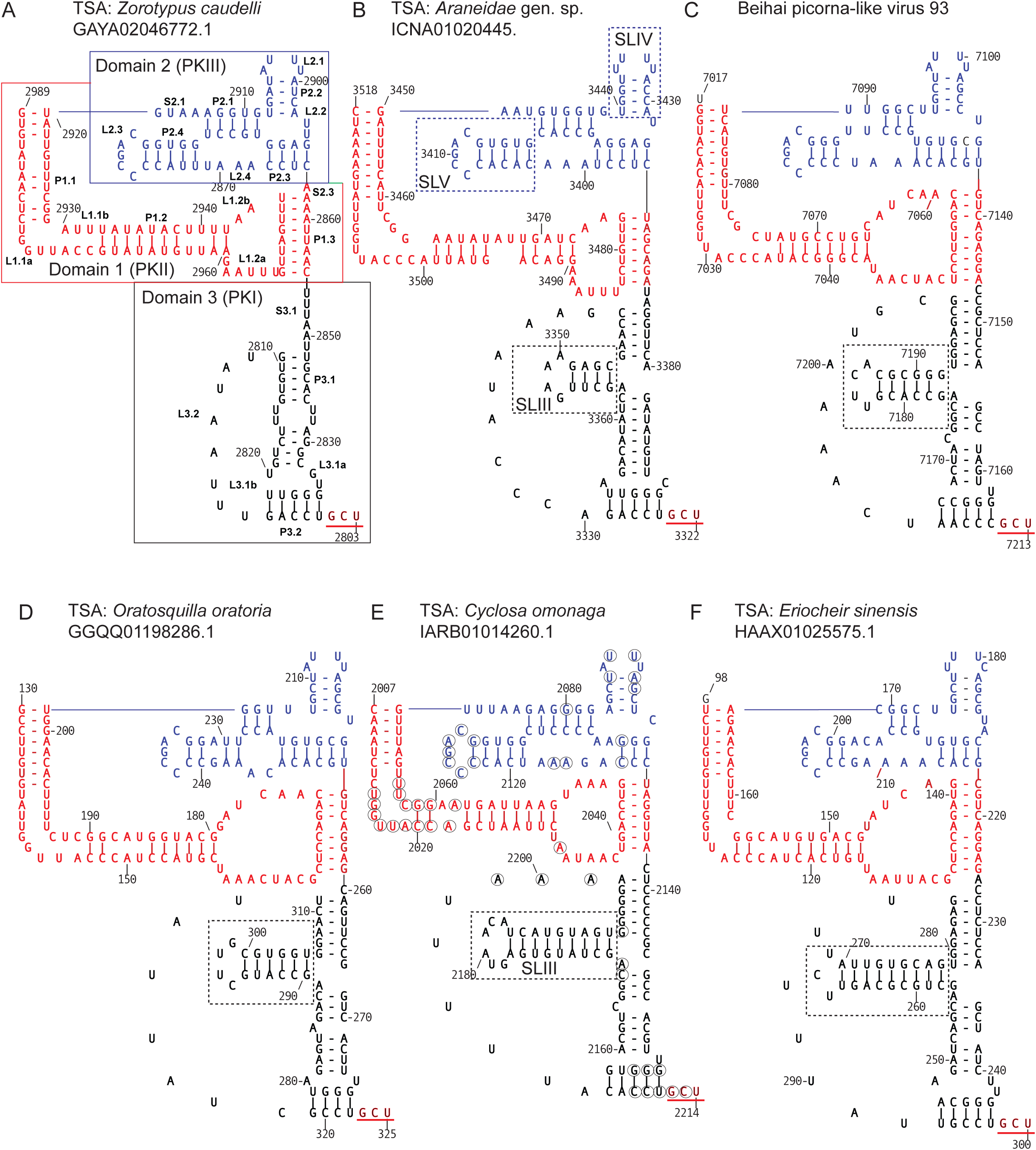
Structures of type 6b IRESs. Models of the type 6b IRESs of (A) TSA: *Zorotypus caudelli* s3164_L_3735_2_a_74_4_l_5165 (GenBank: GAYA02046772.1), (B) TSA: *Araneidae gen*. sp. IDV 2115 (GenBank: ICNA01020445), (C) Beihai picorna-like virus 93 (GenBank: NC_032588.1), (D) TSA: *Oratosquilla oratoria* Unigene 52730_HQ (GenBank: GGQQ01198286.1), (E) TSA: *Cyclosa omonaga* IDV#6323 (GenBank: IARB01014260.1) and (F) TSA: *Eriocheir sinensis* contig Unigene25727_A0A (GenBank: HAAX01025575.1), annotated to show the ORF2 initiation codon (underlined). Domains 1, 2 and 3 are in red, blue, and black font, respectively. (A, B) Models are annotated to show (A) secondary structure elements and domains 1 - 3 and (B) SLIII, SLIV and SLV. Nucleotides are numbered according to the Genbank sequences. (E) Conserved nucleotides in >90% of type 6b IRESs (excluding chimeras) are indicated by black circles.

Sequence alignment indicated that whereas the length and sequence of SLIV and SLV are highly conserved, SLIII is variable. The *Zorotypus caudelli* IRES contains nearly all of the conserved nucleotides that are characteristic of type 6b IRESs but has only two unpaired nucleotides at the SLIII locus (Figure 7A). SLIII in the *Araneidia* gen.sp. IDV 2115 IRES consists of a four bp helix and a 4nt loop (Figure 7B), but it is significantly larger in most other type 6b IRESs, containing five bp in Beihai picorna-like virus 93 (Figure 7C), six bp in Taura syndrome virus [31] and in TSA sequences from *Oratosquilla oratoria* and *Leptomastix dactylii*, and nine bp in TSA sequences from *Cyclosa omonaga* and *Eriocheir sinensis* (Figures 7E, 7F). These observations indicate that SLIII occurs at a locus that tolerates nucleotide insertion and that the ribosomal intersubunit space can accommodate a range of sizes of the SLIII hairpin.

### Identification of classes of type 6 IRESs containing a novel subdomain

The observation that the extended S2.1 spacer sequences in *Latuca sativa* dicistroviridae strain pt151-dic-7 (Figure 4D) and Aparavirus sp. Isolates 220-k141_46450 and 220-k141_171406 (Figure 9E) that link domains 1 and 2 could form small stemloops suggest that this locus in type 6 IRESs has the potential to accommodate a hairpin subdomain. By searching for dicistrovirus sequences that exhibit an abrupt loss of sequence homology with known type 6 IRESs immediately upstream of PKIII, we identified three large groups of type 6 IRESs with hairpins of varying sizes in this spacer. They occurred in dicistrovirus genome fragments derived from a wide range of organisms, including members of seven metazoan phyla (*Annelida*, *Arthropoda*, *Cnidaria*, *Echinodermata*, *Mollusca*, *Porifera* and *Rotifera*). We designated these IRESs type 6i-1 (epitomized by Bundaberg bee virus 2 [47]), type 6i-2 (epitomized by TSA: *H. yanbaruensis*) and type 6i-3 (epitomized by Wenzhou picorna-like virus 28) [46] (Figure 8). Subsets of each of these three IRES types are listed in Tables S6-S8. They all had a conventional three domain/three pseudoknot core in which PKI is directly adjacent to a non-AUG codon (predominantly GCU, GCC or GCA) that initiates the ORF that encodes the capsid protein precursor.

**Figure 8.**
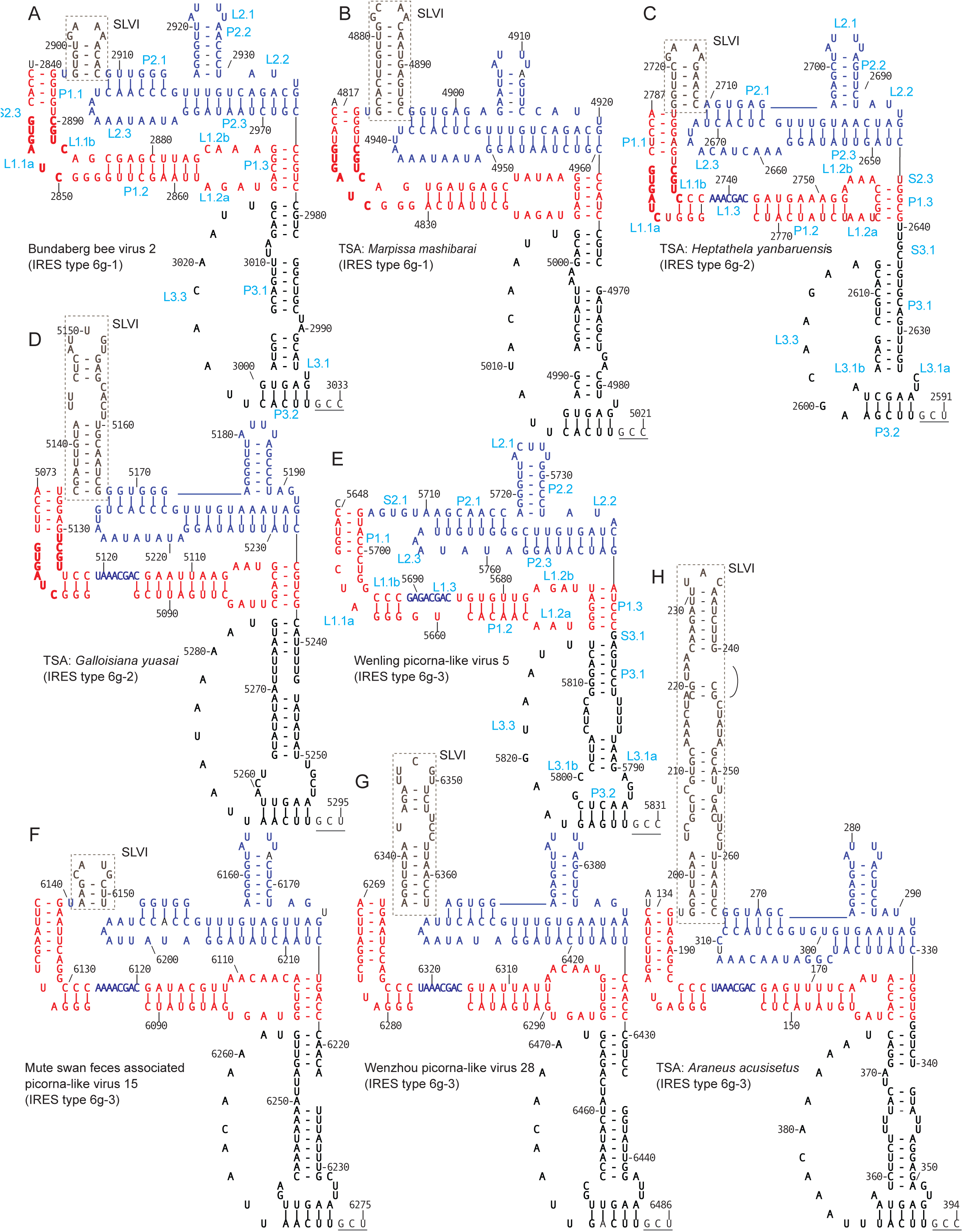
Models of the structure of type 6i IRESs. Models of (a, b) the type 6i-1 IRESs of (a) Bundaberg bee virus 2 isolate QLD-4 (GenBank: MG995700.1) and (b) TSA: *Marpissa mashibarai* (GenBank: IBBC01006465.1), (c, d) the type 6i-2 IRESs of (c) TSA: *Heptathela yanbaruensis* (GenBank: IAWJ01025451.1) and (d) TSA: *Galloisiana yuasai* (GenBank: GAWN02036639.1), and (e - h) the type 6i-3 IRESs of (e) Wenling picorna-like virus 5 strain WLJQ102796 (GenBank: NC_032834.1), (f) Mute swan feces associated picorna-like virus 15 (GenBank: MW588148.1), (g) Wenzhou picorna-like virus 28 strain WZRBX42853 (GenBank: NC_032975.1) and (h) TSA: *Araneus acusisetus* (GenBank: IBEJ01033805.1). Models are annotated to show helices (P), loops (L), single-stranded elements (S), pseudoknots (PK) I, II and III, domains 1, 2 and 3, and where appropriate, stemloop (SL) VI. Nucleotides are numbered at 20nt intervals according to GenBank sequences, and the underlined triplet adjacent to PKI is the first codon of ORF2.

**Figure 9.**
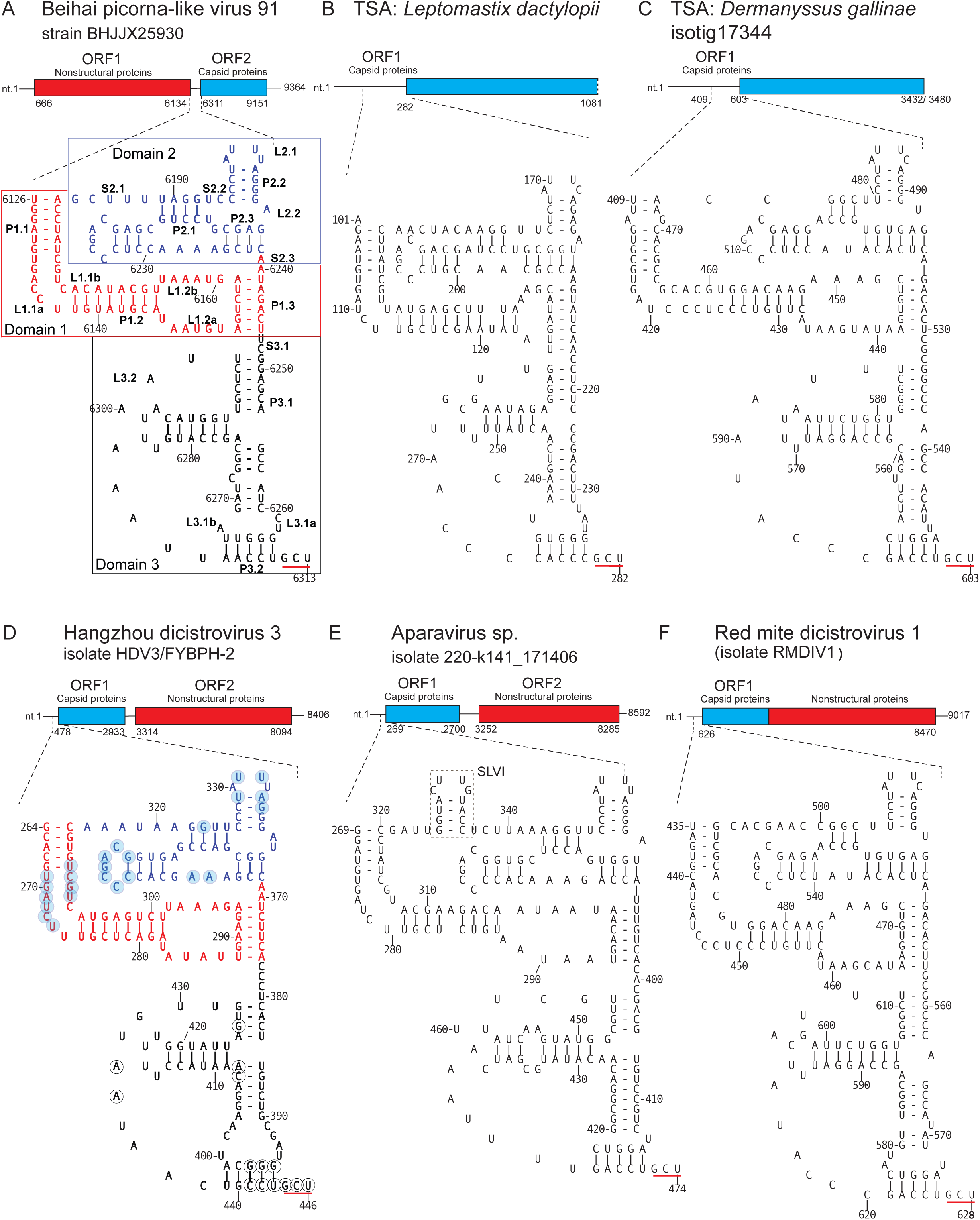
Structural models of chimeric type 6a (domain 1/type 6b (domains 2 +3) IRESs. Models of the chimeric type 6a/type 6b IRESs of (a) Beihai picorna-like virus 91 (GenBank: NC_032638.1), (b) TSA: *Leptomastix dactylopii* (GenBank: GBNE01033170.1), (c) TSA: *Dermanyssus gallinae* (GenBank: GAIF01027790.1), (d) Hangzhou dicistrovirus 3 (GenBank: OM751363.1), (e) MAG: Aparavirus sp. isolate 220-k141_171406 (GenBank: MZ679071.1) and (f) Red mite dicistrovirus 1 isolate RMDIV1 (GenBank: ON160029.1). Panel (a) is annotated to show helices (P), loops (L) and single-stranded elements (S). Panel (b) is annotated to show the SLVI stemloop, pseudoknots (PK) I, II and III, and domains 1, 2 and 3. Nucleotides are numbered at 20nt intervals according to GenBank sequences, and the underlined triplet adjacent to PKI is the first codon of ORF2.

In all three subtypes of 6i IRESs, domain 2/PKIII is the most highly conserved region. It contains an SLIV hairpin that in types 6i-1 and 6i-2 has a 6 base-pair stem and a conserved U-rich apical loop, and in type 6i-3 varies in length from 3 to 8 base-pairs and in many instances, has an apical loop that is not U-rich. All three groups lack a SLV hairpin, and instead have an A-rich L2.3 loop at the equivalent position that resembles the L2.3 loop in some type 6d IRESs (Table S3). The P3.1 helix in domain 3 (PKI) of all three classes is commonly interrupted by mis-paired or unpaired nucleotides, and domain 3 is thus likely more flexible in these IRESs than in other type 6 IRESs. Conserved nucleotides localize to the L3.1 and L3.3 and to the two base-pairs of helix P3.2 adjacent to the start codon. Domain 1/PKII contains a central helix (P1.2), short L1.2a and L1.2b loops and L1.1a and L1.1b loops with GUGAUC and CUGC motifs in IRES types 6i-1 and 6i-2, and shorter, less conserved L1.1a and L1.1b loops in type 6i-3. Type 6i-2 and 6i-3 IRESs contain differ from all other type 6 IRESs in that P1.2 is interrupted by a highly conserved, single stranded motif (AGCAAA in type 6i-2, and AGCAAAG in type 6i-3) that is separated from L1.1a/ L1.1b loops by a stable, conserved minihelix (GGG/CCU in type 6i-2, and GGG/CCC in type 6i-3).

The second distinguishing feature of most type 6i IRESs is that they contain a hairpin (SLVI) in the S2.1 space, immediately adjacent to the P1.1/P1/1’ helix. It is not present in a subset of type 6i-3 IRESs (Table S8) e.g. Wenling picorna-like virus 5 (Figure 8A), but in others, ranges from 10 to ∼70 nt in length (Figure 8). Base-pairing in the helical elements of intermediate and large stem-loops is interrupted by internal loops. The SLVI hairpins in type 6i-1 and type 6i-2 IRESs range from 12 nt to 42 nt in length. Notably, SLVI hairpins in the three types of IRESs are variable and lack conserved loop motifs. For example, the *Ryuthela ishigakiensis*, *Scolopocryptops rubiginosus* and *Sinopoda* sp. type 6i-2 IRESs share 98-99% nucleotide identity excluding SLVI, which ranges in length from 25 to 33nt (Table S7) and is the most variable element in these IRESs. The ∼260 nt-long type 6i-1 IRES in TSA: *Araneus acusisetus* IDV#3440 mRNA (Genbank: IBEJ01033805.1) (Figure 8F) is 98% identical to the IRES in TSA: *Uroctea compactilis* IDV#746 mRNA (IARD01011679.1) and, excluding SLVI, they share ∼80% nucleotide identity with the type 6i-1 IRESs in TSA: *Phycosoma mustelinum* IDV#9 (IBDG01023349.1) and TSA: *Himalaphantes azumiensis* IDV#4087 (IBSV01002144.1) mRNAs. However, highlighting the variability of SLVI in type 6i IRESs, it is 71nt long in *A. acusisetus* and *U. compactilis* IRESs, but only 32nt long in the *P. mustelinum* IRES and 40nt long in the *H. azumiensis* IRES.

### Generation and transfer of chimeric IRESs by recombination

The L1.1a and L1.1b loops in a subset of the IRESs analyzed here (e.g. Behai picorna-like virus 91 (Figure 9A), *Pycnopodia helianthoides* assocated picornavirus 2 and TSA sequences from *Leptomastix dactylopii* (Figure 9B), *Dermanyssus gallinae* (Figure 9C), *Paikiniana keikoae, Metaseilus occidentalis* and *Zelotes potanini*) match the type 6a consensus sequence whereas their domains 2 and 3 contain type 6b-specific structural elements and sequence motifs. These IRESs thus appear to be chimeras formed by recombination of type 6a and type 6b IRESs at a breakpoint that maps near to the S2.1/P2.1 junction. Notably, SLIII in these chimeric IRESs is also variable, as in canonical type 6b IRESs, ranging in length from 5 bp in TSA: *Leptomastix dactylopii* to eight bp in the *D. gallinae* IRES (Table S9; Figures 9B, 9C).

Database searches identified related chimeric IRESs in several dicistrovirus genomes, but instead of occurring in the intergenic region, they are located in the 5’UTRs of genomes in which the order of open reading frames encoding the capsid protein and nonstructural protein precursors is reversed. These IRESs therefore also promote translation of capsid proteins despite their altered genomic location. These genomes include those of Hangzhou dicistrovirus 3 (Figure 9D), Aparavirus sp. Isolate 220-k141 (Figure 9E), Jingmen bat dicistrovirus 1 [62] and PNG bee virus 2 which have dicistronic genomes, *Plasmopara viticola* lesion associated dicistro-like virus 1, which has a tricistronic genome, and Red mite dicistrovirus (Figure 9F) and Jingmen bat dicistrovirus 5 in which capsid and nonstructural protein-coding sequences are fused to form a single open reading frame. Sequence identity in these IRESs ranges from 46% -98%, and they are 182-194 nt long (Table S9), with the exception of Aparavirus sp. isolates 220-k141_171406 (Figure 9E) and 220-k141_46450 (MZ679093.1), which are 204 and 206nt long, respectively. These two IRESs share ∼70% nucleotide sequence identity with the PNG bee virus 2 IRES and they differ principally by the presence of a stemloop in the S2.1 region between domains 1 and 2. Hairpins occur at this locus in some type 6g IRESs, as noted above (Figure 4D).

Domain 1 in the 207 nt-long bivalve RNA virus G2 IRES (Figure 9A) has a conventional size, domain 2 contains an SLIV-type hairpin, and domain 3 includes an SLIII hairpin. Domains 1 and 2 contain nucleotides that match almost all of the nucleotides that are conserved in type 6d IRESs [28], and domain 3 contains some of the conserved nucleotides that are characteristic of type 6b IRESs. This IRES is thus apparently a type 6d/type 6b chimera. An identical IRES occurs in TSA: *Procambarus clarkii* Pc_185626 (GenBank: GBEV01185500.1) and IRESs that are 98 - 99% identical occur in the genomes of Picornavirales sp. isolates 215-k141_21231 (GenBank: MZ678976.1) and 281-k141_163112 (GenBank: MZ678968.1). Taken together, these observations suggest that chimeric IRESs have been formed by recombination of domains derived from IRES types 6a, 6b and 6d, and that recombination has resulted in the transfer of IRESs to novel locations in dicistrovirus genomes.

### Type 6a IRESs located in the coding region of dicistrovirus genomes

Database searches identified numerous type 6a IRES sequences in viral genome fragments that do not correspond to dicistrovirus intergenic regions. They included fragments of monocistronic genomes in which the dicistrovirus capsid and nonstructural protein coding regions are fused. In these genomes, a type 6a IRES is located wholly within the single polyprotein-encoding open reading frame, positioned to promote translation of the capsid protein precursor (Table S10). One group of embedded IRESs is related to the internal IRES of Cripavirus NB-1/2011/HUN [63] and the second is epitomized by the internal *Diversinervus elegans* virus 1 IRES (Figure 9B). They both have a conventional type 6a structure, except that helix P2.4 in the latter comprises only two base-pairs. These viral mRNAs could therefore potentially be translated to yield a polyprotein precursor containing nonstructural and structural proteins and via the IRES, a shorter precursor containing only structural proteins.

### Transfer of type 6a IRESs within and between viral genomes by recombination

In a significant number of dicistrovirus genomes, the order of ORFs is reversed, so that ORF1 encodes a polyprotein precursor of the capsid proteins and the downstream ORF2 encodes the nonstructural protein precursor. The Laverivirus genome [64] contains an IRES upstream of ORF1 that contains almost all of the conserved nucleotides and sequence motifs that are characteristic of type 6a IRESs (Figure 9C). Its presence at the 5’-end of the genome indicates that recombination can result in the transfer of type 6a IRESs to new locations in dicistrovirus genomes and, as noted above for chimeric type 6a/type 6b IRESs such as that of red mite dicistrovirus 1 suggests that other unrelated viral genomes might also contain type 6 IRESs.

BLAST searches also identified type 6 IRESs in several members of the *Tombusviridae*, including isolates s42-k141_944859 (Chen et al., 2022) (Figure 9E), 241-k141_1067879 (GenBank: MZ218436.1), 281-k141_21179 (MZ218468.1), 294-k141_79309 (MZ218531.1), 309R-k141_368388 (MZ218496.1), and the *Solemoviridae* sp. isolate YSN9007 (MW826524.1) [65]. They are ∼180 nt-long and are positioned to promote translation of the capsid protein. The Tombusvirus isolate s42-k141_944859 IRES shares 69 - 75% nucleotide identity with the other *Tombusviridae* IRESs and ∼70% nucleotide identity with type 6a IRESs of *Dicistroviridae* such as Guiyang dicistrovirus 3 (Genbank: MZ209775.1), Wuhan arthropod virus 2 (Figure 6C) and Wufeng shrew dicistrovirus 12 (GenBank: OQ715902.1) [62]. Recombination can therefore lead to the transfer of type 6 IRESs between genomes of *Dicistroviridae* and unrelated RNA viruses. Recombination events that account for the appearance of a chimeric type 6 IRES in another family of RNA viruses are discussed below.

### A structurally distinct group of type 6 IRESs in a novel clade of *Picornaviridae*

Analysis of viral sequences and metagenomic datasets derived from arthropods, environmental samples and plant-associated material revealed 41 members of a structurally distinct group of type 6 IRES in picornavirus-like genomes or genome fragments (Table S11), including those of Hubei Picorna-like virus 51 [46], Robinvale bee virus 6 [47], Cherry virus Trakiya [66], Aphis glycines virus 1 [67] and rice curl dwarf-associated virus [68]. These viruses have recently been identified as members of a novel clade (Ripiresk picorna-like group) that falls between established *Secoviridae* and *Iflaviridae* families of the order *Picornavirales* [69]. These genomes are dicistronic, and the order of open reading frames that encode capsid proteins (ORF1) and non-structural proteins (ORF2) is reversed relative to the order in dicistroviruses. The IGR-like IRESs precede ORF1.

To minimize redundancy, further analysis excluded four IRESs with >97% sequence identity to another member of this group. These IRESs are 144 - 156nt long including the initiation codon, which in different IRESs is AAC/AAU (Asn), ACU/ACA (Thr), AUG (Met), CAA (Gln), GCA/GCU (Ala) or UCA (Ser). They are thus ∼35nt shorter than type 6a IRESs but have a similar predicted three-domain structure (Figure 10D). Domain 1 comprises a pseudoknot (PKII) and relative to type 6a IRESs, has a very short helix P1.2 and internal loops L1.1a/L1.1b and L1.2a/L1.2b. Domain 2 contains a pseudoknot (PKIII) with a protruding SLIV-like hairpin with a pyrimidine-rich loop (L2.2), but has an internal loop (L2.3) instead of SLV. Pairwise sequence identity between these IRESs ranges from 39% to <97%, and there are 52 highly conserved nucleotides (identical in >85% of sequences) that map primarily to domain 2 and adjacent regions of domains 1 and 3 (Figure 10D). These IRESs therefore have structural properties such as the three-domain structure and the conserved SLIV hairpin that occur in other types of IRES and properties that are unique, such as the short P1.2 helix and internal loops in domain 1. They consequently constitute a separate subclass of IGR-like IRES, here designated type 6h.

**Figure 10.**
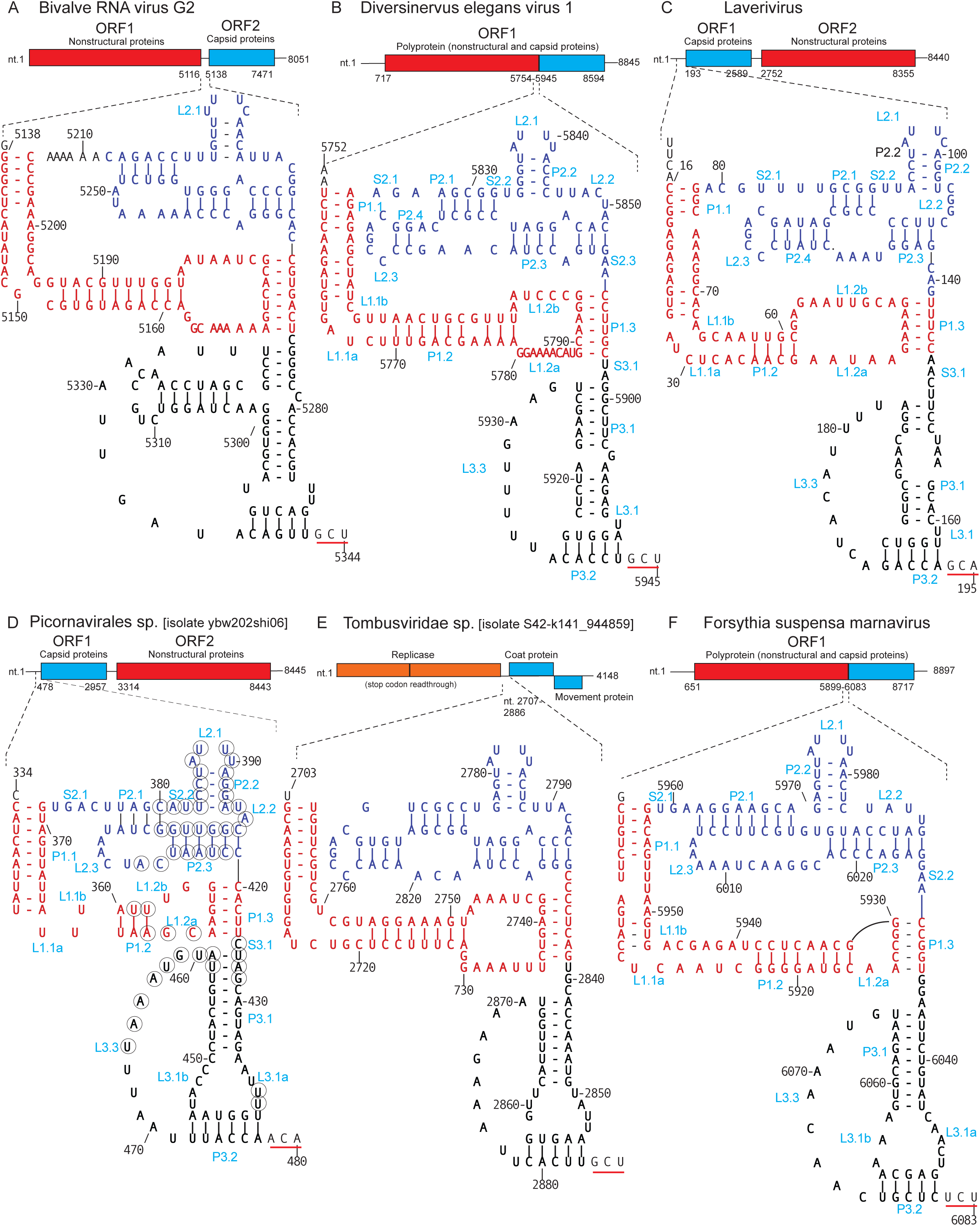
Structural models of chimeric and translocated type 6 IRESs. Models of (a) the chimeric type 6d/type 6b IRES of Bivalve RNA virus G2 (GenBank: NC_032113.1), the type 6a IRESs of (b) *Diversinervus elegans* virus 1 (Genbank: MT127560.1) and (c) Laverivirus (GenBank: KF510029.2), (d) the type 6h IRES of *Picornavirales* sp. Isolate ybw202shi06 (GenBank: MT138149.1), (e) the type 6a IRES of *Tombusviridae* sp. Isolate S42-k141_944859 (GenBank: MZ218538.1) and (f) the type 6i IRES of *Forsythia suspensa* marnavirus strain pt110-mar-7 (GenBank: MN823685.1). Panels (b), (c) and (d) are annotated to show helices (P), loops (L) and single-stranded elements (S) in light blue font. Nucleotides are numbered according to GenBank sequences, and the underlined triplet adjacent to PKI is the first codon of ORF2. (a-e) Models of IRESs are mapped onto schematic representations of viral genomes, labeled to show open reading frames and nucleotides at the 5’ and 5’ ends of genomes or genomic fragments and the first and last nucleotides in open reading frames. (d) Positions of nucleotides that are conserved in >85% of type 6h IRESs are indicated by black circles

### Type 6i-like IRESs in marnavirus genomes

Type 6i-like IRESs were identified in several members of the *Marnaviridae* family (order *Picornavirales*) (Table S12). Marnaviruses infect unicellular aquatic organisms and have monocistronic or dicistronic genomes that encode polyprotein precursors of capsid and non-structural proteins [70]. We identified 193 -194 nt long type 6 IRESs that share 75-90% nucleotide sequence identity in Crogonang virus 17 isolate CGH03, Crogonang virus 61 isolate CGH03 and *Picornavirales* sp. isolate HPLV-37 [71, 72]. Their genomes encode proteins that are characteristic of marnaviruses, and the genomic location of the IRESs positions them to promote translation of the ORF2 capsid protein precursor. The intergenic regions of *Forsythia suspensa* marnavirus strain pt110-mar-7 (Figure 10F) and *Trichosanthes kirilowii* marnavirus strain pt111-mar-8 contain identical 191 nt-long IRESs that share 44-46% nucleotide sequence identity with the three other IRESs. All five IRESs have a conventional three domain structure: the L1.1a and L1.1b loops in domain 1 contain UGC and CAGU motifs, and domain 2 contains an SLIV hairpin but lacks SLV and instead has a large L2.3 loop at the equivalent locus. The region of the *F. suspensa* marnavirus IRES from the S2.1 spacer to the initiation codon is highly homologous to the Wenzhou picorna-like virus 28 IRES and many other type 6i IRESs. The marnavirus IRESs do not contain a SLVI element or the AGCAAA motif in domain 1 that are characteristic of these type 6i IRESs, and they are therefore likely chimeras in which domains 2 and 3 derive from a type 6i-like progenitor. Recombination may therefore have led both to acquisition of type 6i-like IRESs by a subset of marnavirus genomes, and to the exchange of domains between IRESs to generate the structure and sequence motifs in these marnavirus IRESs.

## Discussion

We have identified four new structurally distinct classes of type 6 IRESs that we designated type 6f (e.g. Sanxia picornalike virus 11 (Fig. 2)), type 6g (divided into two subclasses that are exemplified by Weivirus-like virus sp. isolate zftfla02pic2 and Changjiang picornalike virus 6, respectively (Fig. 4)), type 6h (e.g. Hubei picorna-like virus 51 (Fig. 10D; Table S11)) and type 6i (divided into three subclasses, exemplified by Bundaberg bee virus 2, TSA *H. yanbaruensis* and Wenzhou picorna-like virus 28, respectively (Fig. 8)). Each class or subclass is defined by a distinct constellation of sequence and structural characteristics. In addition to IRESs in the intergenic region, we identified type 6 IRESs within the coding region of several dicistroviruses, at the 5’-end of dicistrovirus-like genomes (e.g. Hanzhou dicistrovirus 3) and of picorna-like genomes in the proposed Ripiresk clade of viruses (e.g Robinvale bee virus 6), at internal positions in the genomes of members of the *Marnaviridae* family and between ORFs in the genomes of members of the *Tombusviridae* family. These novel IRESs were identified in viral mRNAs obtained from the eukaryotic phyla *Annelida*, *Arthropoda*, *Cnidaria*, *Echinodermata*, *Mollusca*, *Porifera* and *Rotifera*; a previous study also identified type 6 IRESs in *Bryozoa* and *Entoprocta* [12]. These and other observations [12, 19, 23, 28, 73] show that type 6 IRESs are structurally diverse, and are more widespread than previously recognized, with respect to the virus families in which they occur and the variety of organisms that they infect. For example, type 6f IRESs are only 120-127nt long and contain a very small domain 1 that lacks a P1.2 helix, a compact H-type pseudoknot in domain 2/PKIII that is distinct from other PKIII domains and a conventional domain 3. Type 6c IRESs are 123-129nt long and consist of two pseudoknots [19]. At the other extreme, type 6i IRESs are up to 260 nt long and contain an extended SLIV hairpin, a conserved AGCAAA loop interrupting the P1.2 helix in domain 1 and a highly variable stemloop (SLVI) in the S2.1 spacer adjacent to the P1.1 helix. Neither the AGCAAA loop nor SLVI have previously been identified in type 6 IRESs.

Structural differences between classes of type 6 IRESs influence aspects of their mechanism of action such as their mode of ribosomal binding and the fidelity of reading frame selection. PKI of type 6f IRESs binds stably in the P site of 40S subunits and 80S ribosomes (Fig. 3), consistent with their lack of SLIV and SLV hairpins. These hairpins restrict initial binding of type 6a IRESs to the A site [19]. Binding of PKI of type 6f IRESs in the P site places the first codon of ORF1 in the A site so that translation can commence without the ‘pseudotranslocation’ step that is required by type 6a and 6b IRESs. The principal structural differences in the newly identified classes of type 6 IRESs described here are likely to impact their mechanism of action, and future experiments will focus on determining which aspects of the initiation process are influenced by e.g. the AGCAAA loop and SLVI stemloop of type 6i IRESs. We note that a subset of type 6b IRESs (e.g. Israeli acute paralysis virus) is preceded by a 5’-adjacent stemloop that augments IRES function and contributes to positioning of PKI in the A site [74–76]. The location of SLVI in type 6i IRESs, adjacent to helix P1.1, suggests that it might have a similar function.

Members of each class of type 6 IRES have closely related structures despite a level of sequence identity that may be as low as 26%. Covariant nucleotide substitution has a key role in the maintenance of structure and consequently of function of type 6 and other types of IRES [10–12, 31, 51, 73]. Covariant substitutions are distributed throughout most structural elements of the different subclasses of type 6 IRESs rather than being restricted to specific loci, suggesting that naturally occurring IRESs containing substitutions, reflecting the quasispecies nature of viral populations, retain sufficient residual activity to support viability so that second-site suppressor mutations can appear in progeny. This conclusion is consistent with observations that many single nucleotide substitutions introduced experimentally into type 6 IRESs reduced but did not abrogate their activity (e.g. [12, 22, 28, 75]. Despite changes in genotype (nucleotide substitutions), the phenotype (IRES structure and activity) is therefore mutationally quasi-robust (likely evincing altered thermodynamic stability, but retaining sufficient base-pairing to maintain the structure so that loss of activity is only partial). Mutationally robust phenotypes have high evolvability because tolerance of cryptic genetic variation can permit a spectrum of genetic backgrounds to exist against which additional mutations can manifest themselves and because recombination may juxtapose variant RNA domains with new partners [77–79]. Notably, robustness of genotypes and phenotypes to nucleotide substitution is correlated with their robustness to insertions [80]. This consideration is relevant to the discussion below of the widespread appearance of insertions at specific locations in type 6 IRESs.

Analysis of the sequence and structure of different type 6 IRESs suggests two processes that are responsible for their structural diversification. Subsets of type 6 IRESs can be ordered so that one differs from the next by the presence of a single structural element (Figure 1). Thus, type 6a IRESs contains a P1.2 helix, SLIV and SLV (Fig. 1F), type 6d IRESs contain the P1.2 helix, SLIV but not SLV (Figure 1E), type 6g IRESs contain the P1.2 helix but neither SLIV nor SLV (Figure 1D) and type 6f IRESs do not contain any of these elements (Figure 1C). Similarly, canonical type 6b IRESs contain SLIII in domain 3, which is lacking in the *Zorotypus caudelli* IRES (Figure 7A), even though it contains other conserved nucleotides that are characteristic of type 6b IRESs (Figure 7E). Most type 6i IRESs contain an SLVI hairpin in the S2.1 spacer, but it is absent in a subset of this class (e.g. Wenling picorna-like virus 5)(Figure 8E). A few members of IRES types 6d and 6g-2 (Figures 4D, 8) also contain SLVI hairpins at this locus. Notably, these novel elements do not appear as a unique conserved structure in many of these IRESs but instead occur as a spectrum of variants. Thus, numerous type 6d IRESs have canonical sequence motifs in domain 2 but also contain extended variants of the P2.1 and P2.3 helices and of the L2.3 loop, SLIII in type 6b IRESs ranges from 4 to 9 bp in length, and the SLVI stemloop in type 6i IRESs is extraordinarily variable, containing up to 71 nt in *A. acusisetus* and *U. compactilis* IRESs, varying in size from 25nt to 33nt in length in TSA: *R. ishigakiensis* and several near-identical IRESs, but in other IRESs such as that of TSA: *S. lamarckii*, contains only 9 nt.

We previously suggested that IRES evolution occurs by the cumulative accretion of elements onto a pre-existing scaffold comprising functionally important structural elements and conserved sequence motifs [12, 14]. The observations reported here suggest that in type 6 IRESs, structural expansion occurs at a few defined loci and is likely limited by thermodynamic constraints, by the dimensions of the ribosomal intersubunit space and by the requirement to maintain the integrity of existing functional elements in the IRES. This expansion process is reminiscent of the evolution of ribosomal RNA, in which the sequence and structure of the structural core are conserved but this core is decorated with rapidly diversifying ‘expansion segments’ [81]. More specifically, this expansion process in IRESs corresponds to the process of “lateral helix formation” in which expansion of a bulge loop or spacer forms a new helical element without disrupting the pre-existing structure [82]. Although we cannot exclude a role for imprecise recombination events in the generation of insertions in the IRES, their appearance could also be accounted for by the non-templated insertion of nucleotides during genome replication, initially extending a single-stranded region followed by base-pairing of these nucleotides to form or extend helices and hairpins. Template-dependent reiterative transcription [83] and non-templated insertion of nucleotides [88, 85] have been widely observed, with the latter being influenced by structural elements downstream of the insertion locus and by sequence motifs that promote polymerase slippage.

The sequence and structure of a novel stemloop would be subject to additional selective pressures once it has gained a specific function (such as interaction with a component of the ribosome) in order to optimize the architecture of the IRES and its contribution to this novel function. From this perspective, elements such as SLIV and SLV, which exhibit little variation in length or in the sequence of apical loop residues in different classes of type 6 IRESs, may correspond to the structures that have been optimized (in the case of these elements, for productive interaction with proteins eS25, uS7 and uS11 in the head of the 40S subunit) whereas SLVI is highly variable and may thus constitute a locus at which structural variation is tolerated and has not yet been optimized and become ‘fixed’. Similar hypervariability occurs in type 2 IRESs, in which the Ic hairpin at the apex of domain I is typically 10-15 nt long e.g. in EMCV, foot-and-mouth disease virus and mosavirus A2 IRESs [16, 86], but can be completely absent (as in caprine capripivirus [87] or hyperextended (61nt long, as in Gallivirus A (genus *Gallivirus* of *Picornaviridae*) [88] or even 66-67nt long, in several members of *Caliciviridae* [17]). Similarly, in type 4 IRESs such as HCV, the apical IIIb hairpin is ∼57nt long, but can be much shorter (e.g. 9nt long in members of the *Limnipivirus* genus of *Picornaviridae*) [14] or hyperextended (∼120nt long in members of the *Megrivirus* genus of *Picornaviridae*)([14, 89]. We do not exclude the possibility that the appearance and subsequent optimization of a new structural element with a new function might lead to attrition of elements that determined a function in a progenitor that has been superseded. For example, the UACUA, CUC and GAG elements in PKIII that are invariant in type 6d and type 6f IRESs (which bind to the ribosomal P site) and in type 6g IRESs correspond to elements that are degenerate in some type 6a IRESs (which contain SLIV and SLV elements that determine binding of those IRESs to the A site). The functional requirement for the putatively more ancestral P-site-specific motifs may have been reduced following the appearance of SLIV and SLV.

The second mechanism that contributes to structural diversification of type 6 IRESs is recombination, a process that has previously been recognized as being responsible for the shuffling of domains from type 1 IRESs derived from different co-infecting enteroviruses (e.g. [15] and for the recombination of domains from type 1 and type 2 IRESs to form type 5 IRESs [3, 16, 18, 90]. Here, we identified naturally occurring chimeric IRESs that consist of domains derived from type 6b and type 6a or type 6d IRESs (Figures 9, 10A). Their occurrence is consistent with the viability of chimeric IRESs generated *in vitro* from domains from different classes of type 6 IRESs [19, 34, 91]. Recombination has also been implicated in the transfer of IRESs between genomes of viruses belonging to different genera of *Caliciviridae*, *Flaviviridae* and *Picornaviridae* [11, 13, 14, 16, 17]. Here, we identified type 6 IRESs not only in *Dicistroviridae*, but also in *Tombusviridae*, *Marnaviridae* and the proposed Ripiresk picorna-like group of positive-sense RNA viruses [69], and in other instances, relocated to the 5’-terminal regions of dicistrovirus genomes. Recombination therefore serves to diversify the structure of type 6 IRESs by shuffling domains from different subsets of these IRESs, to move them to different locations in dicistrovirus genomes, and to transfer them horizontally between unrelated genomes.

## Data Availability

All relevant data are available in the manuscript and the supplementary materials.

## Acknowledgements

We thank V. Ramakrishnan for RelE.

## REFERENCES

1. Pestova, T.V., Hellen, C.U. and Shatsky, I.N. (1996) Canonical eukaryotic initiation factors determine initiation of translation by internal ribosomal entry. Mol. Cell. Biol., 16, 6859–6869.

2. de Breyne, S., Yu, Y., Unbehaun, A., Pestova, T.V. and Hellen, C.U. (2009) Direct functional interaction of initiation factor eIF4G with type 1 internal ribosomal entry sites. Proc. Natl. Acad. Sci. USA, 106, 9197–9202.

3. Yu, Y., Sweeney, T.R., Kafasla, P., Jackson, R.J., Pestova, T.V. and Hellen, C.U. (2011) The mechanism of translation initiation on Aichivirus RNA mediated by a novel type of picornavirus IRES. EMBO J., 30, 4423–4436.

4. Pestova, T.V., Shatsky, I.N., Fletcher, S.P., Jackson, R.J. and Hellen, C.U. (1998) A prokaryotic-like mode of cytoplasmic eukaryotic ribosome binding to the initiation codon during internal translation initiation of hepatitis C and classical swine fever virus RNAs. Genes Dev., 12, 67–83.

5. Brown, Z.P., Abaeva, I.S., De, S., Hellen, C.U.T., Pestova, T.V. and Frank, J. (2022) Molecular architecture of 40S translation initiation complexes on the hepatitis C virus IRES. EMBO J., 41, e110581.

6. Wilson, J.E., Pestova, T.V., Hellen, C.U. and Sarnow, P. (2000) Initiation of protein synthesis from the A site of the ribosome. Cell, 102, 511–520.

7. Pestova, T.V. and Hellen, C.U. (2003) Translation elongation after assembly of ribosomes on the Cricket paralysis virus internal ribosomal entry site without initiation factors or initiator tRNA. Genes Dev., 17, 181–186.

8. Jan, E., Kinzy, T.G. and Sarnow, P. (2003) Divergent tRNA-like element supports initiation, elongation, and termination of protein biosynthesis. Proc. Natl. Acad. Sci. USA, 100, 15410–15415.

9. Mears, H.V. and Sweeney, T.R. (2018) Better together: the role of IFIT protein-protein interactions in the antiviral response. J. Gen. Virol., 99, 1463–1477.

10. Jackson, R.J. and Kaminski, A. (1995) Internal initiation of translation in eukaryotes: the picornavirus paradigm and beyond. RNA, 1, 985–1000.

11. Hellen, C.U. and de Breyne, S. (2007) A distinct group of hepacivirus/pestivirus-like internal ribosomal entry sites in members of diverse picornavirus genera: evidence for modular exchange of functional noncoding RNA elements by recombination. J. Virol. 81, 5850–5863.

12. Abaeva, I.S., Young, C., Warsaba, R., Khan, N., Tran, L.V., Jan, E., Pestova, T.V. and Hellen, C.U.T. (2023) The structure and mechanism of action of a distinct class of dicistrovirus intergenic region IRESs. Nucleic Acids Res., 51, 9294–9313.

13. Pisarev, A.V., Chard, L.S., Kaku, Y., Johns, H.L., Shatsky, I.N. and Belsham, G.J. (2004) Functional and structural similarities between the internal ribosome entry sites of hepatitis C virus and porcine teschovirus, a picornavirus. J. Virol., 78, 4487–4497.

14. Asnani, M., Kumar, P. and Hellen, C.U. (2015) Widespread distribution and structural diversity of type IV IRESs in members of *Picornaviridae*. Virology, 478, 61–74.

15. Muslin, C., Joffret, M.L., Pelletier, I., Blondel, B. and Delpeyroux, F. (2015) Evolution and emergence of enteroviruses through intra- and inter-species recombination: plasticity and phenotypic impact of modular genetic exchanges in the 5’ untranslated region. PLoS Pathog., 11, e1005266.

16. Arhab, Y., Bulakhov, A.G., Pestova, T.V. and Hellen, C.U.T. (2020) Dissemination of internal ribosomal entry sites (IRES) between viruses by horizontal gene transfer. Viruses, 12, 612.

17. Arhab, Y., Miścicka, A., Pestova, T.V. and Hellen, C.U.T. (2022) Horizontal gene transfer as a mechanism for the promiscuous acquisition of distinct classes of IRES by avian caliciviruses. Nucleic Acids Res., 50, 1052–1068.

18. Sweeney, T.R., Dhote, V., Yu, Y. and Hellen, C.U. (2012) A distinct class of internal ribosomal entry site in members of the Kobuvirus and proposed Salivirus and Paraturdivirus genera of the *Picornaviridae*. J. Virol. 86, 1468–1486.

19. Abaeva, I.S., Vicens, Q., Bochler, A., Soufari, H., Simonetti, A., Pestova, T.V., Hashem, Y. and Hellen, C.U.T. (2020) The Halastavi árva virus intergenic region IRES promotes translation by the simplest possible initiation mechanism. Cell Rep., 33, 108476.

20. Costantino, D.A., Pfingsten, J.S., Rambo, R.P. and Kieft, J.S. (2008) tRNA-mRNA mimicry drives translation initiation from a viral IRES. Nat. Struct. Mol. Biol., 15, 57–64.

21. Au, H.H., Cornilescu, G., Mouzakis, K.D., Ren, Q., Burke, J.E., Lee, S., Butcher, S.E. and Jan, E. (2015) Global shape mimicry of tRNA within a viral internal ribosome entry site mediates translational reading frame selection. Proc. Natl. Acad. Sci. USA, 112, E6446–6455.

22. Sasaki, J. and Nakashima, N. (1999) Translation initiation at the CUU codon is mediated by the internal ribosome entry site of an insect picorna-like virus in vitro. J. Virol., 73, 1219–1226.

23. Hatakeyama, Y., Shibuya, N., Nishiyama, T. and Nakashima, N. (2004) Structural variant of the intergenic internal ribosome entry site elements in dicistroviruses and computational search for their counterparts. RNA, 10, 779–786.

24. Fernández, I.S., Bai, X.C., Murshudov, G., Scheres, S.H. and Ramakrishnan, V. (2014) Initiation of translation by cricket paralysis virus IRES requires its translocation in the ribosome. Cell, 157, 823–831.

25. Koh, C.S., Brilot, A.F., Grigorieff, N. and Korostelev, A.A. (2014) Taura syndrome virus IRES initiates translation by binding its tRNA-mRNA-like structural element in the ribosomal decoding center. Proc. Natl. Acad. Sci. USA, 111, 9139–9144.

26. Jang, C.J., Lo, M.C. and Jan, E. (2009) Conserved element of the dicistrovirus IGR IRES that mimics an E-site tRNA/ribosome interaction mediates multiple functions. J. Mol. Biol., 387, 42–58.

27. Pfingsten, J.S., Castile, A.E. and Kieft, J.S. (2010) Mechanistic role of structurally dynamic regions in Dicistroviridae IGR IRESs. J. Mol. Biol., 395, 205–217.

28. Miścicka, A., Lu, K., Abaeva, I.S., Pestova, T.V. and Hellen, C.U.T. (2023) Initiation of translation on nedicistrovirus and related intergenic region IRESs by their factor-independent binding to the P site of 80S ribosomes. RNA, 29, 1051–1068.

29. Nakashima, N. and Uchiumi, T. (2009) Functional analysis of structural motifs in dicistroviruses. Virus Res., 139, 137–147.

30. Muhs, M., Yamamoto, H., Ismer, J., Takaku, H., Nashimoto, M., Uchiumi, T., Nakashima, N., Mielke, T., Hildebrand, P.W., Nierhaus, K.H. and Spahn, C.M. (2011) Structural basis for the binding of IRES RNAs to the head of the ribosomal 40S subunit. Nucleic Acids Res., 39, 5264–5275.

31. Nishiyama, T., Yamamoto, H., Shibuya, N., Hatakeyama, Y., Hachimori, A., Uchiumi, T. and Nakashima, N. (2003) Structural elements in the internal ribosome entry site of *Plautia stali* intestine virus responsible for binding with ribosomes. Nucleic Acids Res., 31, 2434–2442.

32. Costantino, D. and Kieft, J.S. (2005) A preformed compact ribosome-binding domain in the cricket paralysis-like virus IRES RNAs. RNA, 11, 332–343.

33. Pfingsten, J.S., Costantino, D.A. and Kieft, J.S. (2007) Conservation and diversity among the three-dimensional folds of the Dicistroviridae intergenic region IRESes. J. Mol. Biol., 370, 856–869.

34. Jang, C.J. and Jan, E. (2010) Modular domains of the Dicistroviridae intergenic internal ribosome entry site. RNA, 16, 1182–1195.

35. Acosta-Reyes, F., Neupane, R., Frank, J. and Fernández, I.S. (2019) The Israeli acute paralysis virus IRES captures host ribosomes by mimicking a ribosomal state with hybrid tRNAs. EMBO J., 38, e102226.

36. Zuker, M. (2003) Mfold web server for nucleic acid folding and hybridization prediction. Nucleic Acids Res., 31, 3406–3415.

37. Janssen, S. and Giegerich, R. (2015) The RNA shapes studio. Bioinformatics, 31, 423–425.

38. Andronescu, M., Condon, A., Hoos, H.H., Mathews, D.H. and Murphy, K.P. (2007) Efficient parameter estimation for RNA secondary structure prediction. Bioinformatics, 23, i19–28.

39. Sato, K., Kato, Y., Hamada, M., Akutsu, T. and Asai, K. (2011) IPknot: fast and accurate prediction of RNA secondary structures with pseudoknots using integer programming. Bioinformatics, 27, i85–93.

40. Popenda, M., Szachniuk, M., Antczak, M., Purzycka, K.J., Lukasiak, P., Bartol, N., Blazewicz, J. and Adamiak, R.W. (2012) Automated 3D structure composition for large RNAs. Nucleic Acids Res., 40, e112.

41. Pettersen, E.F., Goddard, T.D., Huang, C.C., Meng, E.C., Couch, G.S., Croll, T.I., Morris, J.H. and Ferrin, T.E. (2021) UCSF ChimeraX: Structure visualization for researchers, educators, and developers. Protein Sci., 30, 70–82.

42. Pisarev, A.V., Unbehaun, A., Hellen, C.U. and Pestova, T.V. (2007) Assembly and analysis of eukaryotic translation initiation complexes. Methods Enzymol., 430, 147–177.

43. Zinoviev, A., Ayupov, R.K., Abaeva, I.S., Hellen, C.U.T. and Pestova, T.V. (2020) Extraction of mRNA from stalled Ribosomes by the Ski complex. Mol. Cell, 77, 1340–1349.e6.

44. Neubauer, C., Gao, Y.G., Andersen, K.R., Dunham, C.M., Kelley, A..C, Hentschel, J., Gerdes, K., Ramakrishnan, V. and Brodersen, D.E. (2009) The structural basis for mRNA recognition and cleavage by the ribosome-dependent endonuclease RelE. Cell, 139, 1084–95.

45. Skabkin, M.A., Skabkina, O.V., Hellen, C.U. and Pestova, T.V. (2013) Reinitiation and other unconventional posttermination events during eukaryotic translation. Mol. Cell, 51, 249–264.

46. Shi, M., Lin, X.D., Tian, J.H., Chen, L.J., Chen, X., Li, C.X., Qin, X.C., Li, J., Cao, J.P., Eden, J.S. et al. (2016) Redefining the invertebrate RNA virosphere. Nature 540, 539–543.

47. Roberts, J.M.K., Anderson, D.L. and Durr, P.A. (2018) Metagenomic analysis of Varroa-free Australian honey bees (*Apis mellifera)* shows a diverse *Picornavirales* virome. J. Gen. Virol., 99, 818–826.

48. Chen, Y.M., Sadiq, S., Tian, J.H., Chen, X., Lin, X.D., Shen, J.J., Chen, H., Hao, Z.Y., Wille, M., Zhou, Z.C. et al. (2022) RNA viromes from terrestrial sites across China expand environmental viral diversity. Nat. Microbiol., 7, 1312–1323.

49. Schüler, M., Connell, S.R., Lescoute, A., Giesebrecht, J., Dabrowski, M., Schroeer, B., Mielke, T., Penczek, P.A., Westhof, E. and Spahn, C.M. (2006) Structure of the ribosome-bound cricket paralysis virus IRES RNA. Nat. Struct. Mol. Biol., 13, 1092–1096.

50. Babendure, J.R., Babendure, J.L., Ding, J.H. and Tsien, R.Y. (2006) Control of mammalian translation by mRNA structure near caps. RNA, 12, 851–861.

51. Jan, E. and Sarnow, P. (2002) Factorless ribosome assembly on the internal ribosome entry site of cricket paralysis virus. J. Mol. Biol., 324, 889–902.

52. Myasnikov, A.G., Kundhavai Natchiar, S., Nebout, M., Hazemann, I., Imbert, V., Khatter, H., Peyron, J.F. and Klaholz, B.P. (2016) Structure-function insights reveal the human ribosome as a cancer target for antibiotics. Nat. Commun., 7, 12856.

53. Andreev, D., Hauryliuk, V., Terenin, I., Dmitriev, S., Ehrenberg, M. and Shatsky, I. (2008) The bacterial toxin RelE induces specific mRNA cleavage in the A site of the eukaryote ribosome. RNA, 14, 233–239.

54. Alkalaeva, E.Z., Pisarev, A.V., Frolova, L.Y., Kisselev, L.L. and Pestova, T.V. (2006) In vitro reconstitution of eukaryotic translation reveals cooperativity between release factors eRF1 and eRF3. Cell, 125, 1125–1136.

55. Brown, A., Shao, S., Murray, J., Hegde, R.S. and Ramakrishnan, V. (2015) Structural basis for stop codon recognition in eukaryotes. Nature, 524, 493–496.

56. Muhs, M., Hilal, T., Mielke, T., Skabkin, M.A., Sanbonmatsu, K.Y., Pestova, T.V. and Spahn, C.M. (2015) Cryo-EM of ribosomal 80S complexes with termination factors reveals the translocated cricket paralysis virus IRES. Mol. Cell, 57, 422–432.

57. Liljas, L., Tate, J., Lin, T., Christian, P. and Johnson, J.E. (2002) Evolutionary and taxonomic implications of conserved structural motifs between picornaviruses and insect picorna-like viruses. Arch. Virol., 147, 59–84.

58 Jia, H. and Gong, P. (2019) A structure-function diversity survey of the RNA-dependent RNA polymerases from the positive-strand RNA viruses. Front. Microbiol., 10, 1945.

59. Yuan, X. and Kadowaki, T. (2022) DWV 3C protease uncovers the diverse catalytic triad in insect RNA viruses. Microbiol. Spectr., 10, e0006822.

60. Yang, S., Mao, Q., Wang, Y., He, J., Yang, J., Chen, X., Xiao, Y., He, Y., Zhao, M., Lu, J. et al. (2020) Expanding known viral diversity in plants: virome of 161 species alongside an ancient canal. Environ. Microbiome, 17, 58.

61. Abeyrathne, P.D., Koh, C.S., Grant, T., Grigorieff, N. and Korostelev, A.A. (2016) Ensemble cryo-EM uncovers inchworm-like translocation of a viral IRES through the ribosome. Elife, 5, e14874.

62. Chen, Y.M., Hu, S.J., Lin, X.D., Tian, J.H., Lv, J.X., Wang, M.R., Luo, X.Q., Pei, Y.Y., Hu, R.X., Song, Z.G., Holmes, E.C. and Zhang, Y.Z. (2023) Host traits shape virome composition and virus transmission in wild small mammals. Cell, 186, 4662–4675.e12.

63. Reuter, G., Pankovics, P., Gyöngyi, Z., Delwart, E. and Boros, A. (2014) Novel dicistrovirus from bat guano. Arch. Virol., 159, 3453–3456.

64. Greninger, A.L. and DeRisi, J.L. (2015) Draft genome sequence of Laverivirus UC1, a dicistrovirus-like RNA virus featuring an unusual genome organization. Genome Announc., 3, e00656–15.

65. Zhu, W., Yang, J., Lu, S., Jin, D., Pu, J., Wu, S., Luo, X.L., Liu, L., Li, Z., Xu, J. (2020) RNA virus diversity in birds and small mammals from Qinghai-Tibet plateau of China. 2022. Front. Microbiol., 13, 780651.

66. Milusheva, S., Phelan, J., Piperkova, N., Nikolova, V., Gozmanova, M. and James, D. (2019) Molecular analysis of the complete genome of an unusual virus detected in sweet cherry (*Prunus avium*) in Bulgaria. Eur. J. Plant Pathol., 153, 197–207.

67. Yasmin, T., Thekke-Veetil, T., Hobbs, H.A., Nelson, B.D., McCoppin, N.K., Lagos-Kutz, D., Hartman, G.L., Lambert, K.N., Walker, D.R. and Domier, L.L. (2020) Aphis glycines virus 1, a new bicistronic virus with two functional internal ribosome entry sites, is related to a group of unclassified viruses in the *Picornavirales*. J. Gen. Virol., 101, 105–111

68. Zhang, T., Li, C., Cao, M., Wang, D., Wang, Q., Xie, Y., Gao, S., Fu, S., Zhou, X., Wu, J. (2021) A novel rice curl dwarf-associated picornavirus encodes a 3C serine protease recognizing uncommon EPT/S cleavage sites. Front. Microbiol., 12, 757451.

69. Sadiq, S., Harvey, E., Mifsud, J.C.O., Minasny, B., McBratney, A.B., Pozza, L.E., Mahar, J.E. and Holmes, E.C. (2024) Australian terrestrial environments harbour extensive RNA virus diversity. Virology, 593, 110007.

70. Lang, A.S., Vlok, M., Culley, A.I., Suttle, C.A., Takao, Y., Tomaru, Y., ICTV Report Consortium. (2021) ICTV Virus Taxonomy Profile: *Marnaviridae* 2021. J. Gen. Virol., 102, 001633.

71. Zell, R., Groth, M., Selinka, L. and Selinka, H.C. (2022) Picorna-like viruses of the Havel river, Germany. Front. Microbiol., 13, 865287.

72. Richard, J.C., Blevins, E., Dunn, C.D., Leis, E.M. and Goldberg, T.L. (2023) Viruses of freshwater mussels during mass mortality events in Oregon and Washington, USA. Viruses, 15, 1719.

73. Kanamori, Y. and Nakashima, N. (2001) A tertiary structure model of the internal ribosome entry site (IRES) for methionine-independent initiation of translation. RNA, 7, 266–274.

74. Firth, A.E., Wang, Q.S., Jan, E. and Atkins, J.F. (2009) Bioinformatic evidence for a stem-loop structure 5’-adjacent to the IGR-IRES and for an overlapping gene in the bee paralysis dicistroviruses. Virol J., 6, 193.

75. Ren, Q., Wang, Q.S., Firth, A.E., Chan, M.M., Gouw, J.W., Guarna, M.M., Foster, L.J., Atkins, J.F. and Jan, E. (2012) Alternative reading frame selection mediated by a tRNA-like domain of an internal ribosome entry site. Proc. Natl. Acad. Sci. USA, 109, E630–639.

76. Au, H.H.T., Elspass, V.M. and Jan, E. (2018) Functional insights into the adjacent stem-Loop in honey bee dicistroviruses that promotes internal ribosome entry site-mediated translation and viral infection. J. Virol., 92, e01725–17.

77. Wagner, A. (2008) Robustness and evolvability: a paradox resolved. Proc. Biol. Sci., 275, 91–100.

78. Hayden, E. J., Ferrada, E. and Wagner, A. (2011) Cryptic genetic variation promotes rapid evolutionary adaptation in an RNA enzyme. Nature, 474, 92–95.

79. Payne, J.L. and Wagner, A. (2019) The causes of evolvability and their evolution. Nat. Rev. Genet., 20, 24–38.

80. Martin, N.S. and Ahnert, S.E. (2021) Insertions and deletions in the RNA sequence-structure map. J. R. Soc. Interface, 18, 20210380.

81. Bowman, J.C., Petrov, A.S., Frenkel-Pinter, M., Penev, P.I. and Williams, L.D. (2020) Root of the tree: the significance, evolution, and origins of the ribosome. Chem. Rev., 120, 4848–4878.

82. Caisová, L. and Melkonian, M. (2014) Evolution of helix formation in the ribosomal Internal Transcribed Spacer 2 (ITS2) and its significance for RNA secondary structures. J. Mol. Evol., 78, 324–337.

83. Kempf, B.J., Kelly, M.M., Springer, C.L., Peersen, O.B. and Barton, D.J. (2013) Structural features of a picornavirus polymerase involved in the polyadenylation of viral RNA. J. Virol., 87, 5629–5644.

84. Olspert, A., Chung, B.Y., Atkins, J.F., Carr, J.P. and Firth, A.E. (2015) Transcriptional slippage in the positive-sense RNA virus family Potyviridae. EMBO Rep., 16, 995–1004.

85. Stewart, H., Olspert, A., Butt, B.G. and Firth, A.E. (2019) Propensity of a picornavirus polymerase to slip on potyvirus-derived transcriptional slippage sites. J. Gen. Virol., 100, 199–205.

86. Pilipenko, E.V., Blinov, V.M., Chernov, B.K., Dmitrieva, T.M. and Agol, V.I. (1989) Conservation of the secondary structure elements of the 5’-untranslated region of cardio- and aphthovirus RNAs. Nucleic Acids Res., 17, 5701–5711.

87. Boros, Á., Pankovics, P., László, Z., Urbán, P., Herczeg, R., Gáspár, G., Tóth, F. and Reuter, G. (2023) The genomic and epidemiological investigations of enteric viruses of domestic caprine (*Capra hircus*) revealed the presence of multiple novel viruses related to known strains of humans and ruminant livestock species. Microbiol. Spectr., 11, e0253323.

88. Boros, Á., Nemes, C., Pankovics, P., Kapusinszky, B., Delwart, E. and Reuter, G. (2012) Identification and complete genome characterization of a novel picornavirus in turkey (*Meleagris gallopavo*). J. Gen. Virol., 93, 2171–2182.

89. Boros, Á., Pankovics, P., Knowles, N.J., Nemes, C., Delwart, E. and Reuter, G. (2014) Comparative complete genome analysis of chicken and Turkey megriviruses (family *Picornaviridae*): long 3’ untranslated regions with a potential second open reading frame and evidence for possible recombination. J. Virol., 88, 6434–6443.

90. Pankovics, P., Boros, Á., Phan, T.G., Delwart, E. and Reuter, G. (2018) A novel passerivirus (family Picornaviridae) in an outbreak of enteritis with high mortality in estrildid finches (*Uraeginthus* sp.). Arch. Virol., 163, 1063–1071.

91. Hertz, M.I. and Thompson, S.R. (2011) In vivo functional analysis of the Dicistroviridae intergenic region internal ribosome entry sites. Nucleic Acids Res., 39, 7276–7288.

