## Supplemental Tables 1 - 12 for "Genetic mechanisms underlying the structural elaboration and dissemination of viral internal ribosomal entry sites"

Table S1. Sequence alignment of type 6f IRESs.

| | | P1.1 P1.2 P2.1 P2.2 |
|---|---|---|
| 1 | Perth Bee | 5707 ---GGUAUG-GCACAUAAUUCAUGACUACGGU--CAUGCCC--AGAACGUCAACUACUCAGACGAU |
| 2 | Changjiang7 | 5598 ---GGCGCA-GCACAUAAUUCACAACUGCGGU--UGUGCCC--AGGGCGUCGACUACUCAGACGUA |
| 3 | Sanxia11 | 5361 --AAGCAUUGACACAUAUGUCAUUGCCCCGGUUAUAUGCUU-AAGUGAGUCUACUACUCAGACUUU |
| 4 | Hubei19 | 5272 ---AGCACU-ACACAUACUUUGUGUUCCCGGU---GUGCU--AACCAUGUCAACUACUCAGACGUA |
| 5 | Picornav. | 5396 CUAAUAUGUUGUACAUAAUUCACGGUUGCGGUGACAUAUUAG----ACGUCAACUACUCAGACGUU |
| | | L1.1a L1.1b S2.1 L2.1 |

| | | P1.2 P2.2 P3.2 P2.1 |
|---|---|---|
| 1 | Perth Bee | AAGAGAGAGGAUAGUCAUGCUGUGGGUGUCACUUGAGCU-GUGCCCCAUAAGUAAAAUAGCUUUUC 5829 |
| 2 | Changjiang7 | AAGAGAGAGGACAGUUGUGCAGCAUAG-AGGUUUGAGCAGACGCUAUGUUAGUAAAAU-GCUUUUC 5719 |
| 3 | Sanxia11 | AAGAGAGAGGAGGGCGAUGCGGUGUGCU-UGUUUGAGCG-ACAGCACACAUGAAAUUUAGCUUACA 5487 |
| 4 | Hubei19 | AAGAGAGAGGAGGAACACACUAUGUGCU-UGUUUGAGCG-ACGGCACAUUAGU-AUAUAGCUUUCU 5391 |
| 5 | Picornav. | AAGAGAGAGGACAAUCGUGCCUUGUGGG-UAUGUGAGCG-AUCCCACAUUUGU-AUAUAGCUUCCU 5520 |
| | | L2.2 S2.2 S3.1 L3.1 L3.2 L3.3 |

Aligned sequences from (1) Perth bee virus 2 isolate WA1-14 (GenBank: MG995726.1), (2) Changjiang picorna-like virus 7 strain CJLX30780 (NC_032860.1), (3) Sanxia picorna-like virus 11 strain SXXX37695 (NC_033221.1), (4) Hubei picorna-like virus 19 strain SZmix17531 (KX883724.1) and (5) MAG: Picornavirales sp. isolate 77-k141_404334 (MZ679003.1), annotated to show nucleotide numbers, structural elements (labelled as in Figure 1), the ORF2 initiation codon (indicated by red font) and conserved nucleotides (indicated by bold font).

Table S2. (A) Sequence and structural alignment of type 6g-1 IRESs.

| # | Viral sequence | P1.1 / P1.2 / P1.3 |
|---|---|---|
| 1 | Gingko-dic13 | 5770 GACCUCAUAUACGGCAGCCUCA-UGUACAUACC-GUGUUCCGGCAUGUUGGCUGUGUCAGGAGAGGGUCC- |
| 2 | Lactuca-dic19 | 5612 UGACCUUAUAUACGCCCAUUU-ACUGUACAUACCGAAGUUGCGGCA-GGAAUUGGGGACAGGAU-GGGUCA- |
| 3 | Wenzhou33 | 5439 UAGGCCCAUACACGCAUUCACGCA-GUACAUAGCGUCUUGGCGGC-UGCGUUGGGUGGCAGGACAGCCUAC- |
| 4 | Biomphalaria | 5471 AGCGCCCAUAUACGCAUCCCGCAUUGUAAAUAA-GUUUCAGCGGCAUAUCAGGAUGG-CAGGAUAGCGCUU- |
| 5 | Wei_zftfla | 5836 GGGAUAUAUACGGUUCCACAAGUGUACAUAAUUUGCGAUCGGCUAUUGUGGAAUGUCAGGAUGACCCCAA |
| 6 | Crogonan | 2167 UAGCUUAUAUACGCAUGCGUUCCCGUCCAUAGCGGAACUUCGGCGGG-GCGUGUGGUCAGGAG-AGCUA |
| | | L1.1a L1.2a L1.2b L1.1b S2.1 |
| | | P1.3 P2.1 P2.2 P3.1 P3.2 |
| 1 | Gingko-dic13 | CCG-UACUACUCA-GAGGAU--UCAGAGAG-AGAACACAAC-UUUUGGCGUUUGUGUAGGCGCCAA---UUGUACAUAGUUUACACGCU 5917 |
| 2 | Lactuca-dic19 | CCGUUACUACUCA-GAGGAU--UCAGAGAGAACAACUUCAC--CAGUGUUCGCGUGUGCGCGCGGCACUGUGUACAUAG--AGCACGCU 5761 |
| 3 | Wenzhou33 | CCGUUACUACUCA-GAGGAU--UCAGAGAGUACCAAGACACUUACAUUGGCUUGUGCGAGCCAUGUA---UGUACAUAG-UAGCACGCG 5589 |
| 4 | Biomphalaria | UCCGUUACUACUCA-GAGGAU--UCAGAGAGCACUGAAACAACUUGUCGCGCCAGUGCUAGCGCGGCA--UUUAUCAUAG--AGCACGCG 5621 |
| 5 | Wei_zftfla | AGUCUUUACUACUCACGACUAUAAAGAGAGAGUAAUCGCACAC-UCCAAGGACGUGGAUUAAUCCUUGG---UUUACAUAG--AAUCCGCU 5990 |
| 6 | Crogonan | CCGUUACUACUCA-GGGGUU--CCAGAGAGUAGAGUUCCACCCACGUAUGCUUGUGCGAGCG-ACGUGG-UGUACAUAC--AGCACGCU 2315 |
| | | L2.1 L2.2 L2.3 L2.4 S3.1 L3.1a L3.1b L3.2 |

Table S2. (A) Sequence alignment of type 6g-1 IRESs, numbered to indicate 5' and 3' terminal nucleotides and annotated to show structural elements (as shown in Fig. 3A), the initiation codon (red font) and nucleotides conserved in over 83% of members of this group (bold). Nucleotides that are predicted to engage in base-pairing in helical elements P1.1, P1.2, P1.3, P2.1, P2.2, P3.1 and P3.2 are shaded and bracketed. Aligned sequences are from: (1) Ginkgo biloba dicistrovirus strain pt112-dic-13 (MN720005.1), (2) Lactuca sativa dicistroviridae strain pt151-dic-19 (MN722415.1), (3) Wenzhou picorna-like virus 33 strain WZFSL76563 (NC_032939.1), (4) MAG: Biomphalaria pfeifferi virus 2 isolate BP-Kasabong (MT108489.1), (5) MAG: Weivirus-like virus sp. isolate zftfla02pic2 (MT138418.1), and (6) MAG: Crogonang virus 38 isolate CGH03 (OR270296.1).

Table S2. (B) Sequence and structural alignment of type 6g-2 IRESs.

| | | |
|---|---|---|
| | | P1.1 P1.2 P1.3 |
| 1 | Lactuca_dic18 | 5319 --GUCUGGUUUUAAAGUUGGGUGGUACACAUAAUUGACAGAACCGG-UGCCAUCCACACCAUAUUGGUUG |
| 2 | Crawfish 3 | 5382 --GCGUGGUUUUA---AGCCAUGACUUACAUACUUGA-AGACCCGGAGUGUGUGGCUGCCACAUUGUUGCUUGGUUA |
| 3 | Wei_swa066 | 5606 --GCGUGGUUUUA---AGCCAUGACUUACAUACUUGA-AGACCCGGAGUGUGUGGCUGCCACAUUGUUGCUUGGUUA |
| 4 | Changjiang 6 | 5542 AUGUAUGGUUUUAGUU--GUGGACUGCACAUAAAUGGUUGA--CGG-CGGUUCACUUACCAUGUAUUCUUCGUGGU |
| 5 | Crawfish 2 | 5683 AUGUAUGGUUUUAGAAGUAUAACUUGUACAUAAGUGA-UGCACCGG-CAUGUUAU-UGCCACAAUGUGGUAU |
| 6 | Chanjiang 2 | 2683 AUGUAUGGUUUUAGAAGUAUAACUUGUACAUAGGUGA-UGCACCGG-CAUGUUAU-UGCCACAAUGUGGUAC |
| 7 | Wenzhou 31 | 5674 AUGUUUGGUUUUAA--AGAACUUGUCCACAUAUUUGAU-GCUUCGG--GAUAAGUGAGCCUCCUUGUUGUCGAGGUGA |
| 8 | Crogonang 136 | 5242 CGAUGUGGUUUUAA-AGUAUAACCUGCACAUGAUUGAU-GCAACGG-CAUGUUAUA-GCCACUGUGUGGUAA |
| | | L1.1a L1.2a L1.2b L1.1b S2.1 |

| | | |
|---|---|---|
| | | P1.3 P2.1 P2.2 P3.1. P3.2 |
| 1 | Lactuca_dic18 | UGCGACUACUCAGCG---AAGUGAGAGGAGUUUUGUCCGUGAUGGGUGUU--UGAGCAGACCCCAUU-UUGUAUUUUGCUUGCU 5463 |
| 2 | Crawfish 3 | UGACCACUACUCAGUCAGUAAGAGAGAGGAGGUCU-UCCUUGGAUCUUUGC-GUGAGCUGCGAGAUCCGUUGUAAAUAGCUUACU 5535 |
| 3 | Wei_swa066 | UGACCACUACUCAGUCACUAAGAGAGAGGAGGUCU-UCCUUGGGUCUUUGC-GUGAGCUGCGAGAUCCGUUGUAAAUAGCUUACU 5759 |
| 4 | Changjiang 6 | GGACAACUACUCAGUCCUUAAGAGAGAGGAA-UCAACCCCAGGGGCAAUCU-UUGAGCC-AGGUGCCCCGUGGAUAUGGCUUACA 5695 |
| 5 | Crawfish 2 | AGACAACUACUCAGUCUUUAAUAGAGAGGAGUGCA-UCCGAGACCGUGAUUCGUGAGCU-GAACCUGGUGUGUAAAUAGCUUACA 5834 |
| 6 | Chanjiang 2 | AGACAACUACUCAGUCUUAAUAGAGAGGAGUGCAUUCGA-ACCAGUGUCUUUUGAGCU-GAGCCUGGUGUGUAAAUAGCUUACA 2834 |
| 7 | Wenzhou 31 | AGACUACUACUCAGUCUUUAAAAGAGAGGAGAGCG-UCCAGAGUGUGUGUCU-UGAGCA-GACCACACUUUAGUAUUUGCUUACA 5826 |
| 8 | Crogonang 136 | AAGACGACUACUCAGUCUUUAAGAGAGAGGAGUGCA-UCCGAACCAGAGUCUCGUGAGCG-GAGCCUGGUGUGUAUAUAGCUUACA 5394 |
| | | L2.1 L2.2 L2.3 L2.4 S3.1 L3.1a L3.1b L3.2 |

Table S2. (B) Sequence alignment of type 6g-2 IRESs, numbered to indicate 5' and 3' terminal nucleotides and annotated to show structural elements (as shown in Fig. 3B), the initiation codon (red font) and nucleotides conserved in over 87% of members of this group (bold). Nucleotides that are predicted to engage in base-pairing in helical elements P1.1, P1.2, P1.3, P2.1, P2.2, P3.1 and P3.2 are shaded and bracketed.

Aligned sequences are from: (1) Lactuca sativa dicistroviridae strain pt151-dic-18 (MN722414.1), (2) Changjiang crawfish virus 3 strain CJLX30786 (NC_032868.1), (3) MAG: Weivirus-like virus sp. isolate swa066shi2 (MN917675.1), (4) Changjiang picorna-like virus 6 strain CJLX30470 (NC_032794.1), (5) Changjiang crawfish virus 2 strain CJLX30764 (NC_032850.1), (6) MAG: Chanjiang crawfish virus 2 isolate ML176 clone mfalml176 (OR270417.1), (7) Wenzhou picorna-like virus 31 strain WZFSL72092 (NC_032959.1), and (8) MAG: Crogonang virus 136 isolate GCR11 (OR270240.1).

Table S3. Sequence alignments of L1.1a and L1.1.b loops and domain 2 of type 6d IRESs.

| # | Virus | Loop L1.1a | Loop L1.1b | Domain 2 | |
|---|---|---|---|---|---|
| | | | | P2.1 / P2.2/L2.1 / P2.3 | |
| 1 | NediV | UAUACGG | UGACAGG | AUUUCACCCCCAUUUUCAUGG-UACUACUCAGGUUGG--------------ACCAGAGAA | 1444 |
| 2 | APLV1 | UAUAUACG | GUCAGGAC | GCAUUGCCCCUGCUUUUUAGC-UUACUACUCAGACGGA--------------GUCAGAGAAU | 5555 |
| 3 | Pallasea | UAUACACG | GUCAGGAC | GCAAUGCCCUUGCUUUUUAGC-UUACUACUCAGACUGA--------------GUCAGAGAAU | 1083 |
| 4 | Chemarfal virus 33 | CCACAUACG | GUCAGGACA | UGCCCUUGCUUUUUAGC-UUACUACUCAGACUGA--------------GUCAGAGAA | 5367 |
| 5 | Changjiang 9 | AAAUACG | GACAGG | CCUUUAAACCU-AGCUUUUCAGCUUUACUACUCAGGCGGA--------------GCCAGAGAG | 5667 |
| 6 | Beihai78 | AUAUACG | GUCAGG | UCCCCCCUGCCUUU-GGC-UUACUACUCAGGUGGA--------------ACCAGAGAGCA | 5579 |
| 7 | Gingko pt112d11 | AUAUACG | GUCAGGA | AUUCGCCCUCCAUUUCUAUGG-UACUACUCAGGUUGG--------------ACCAGAGAA | 5946 |
| 8 | Hangzhou-4 | UAUACG | GUCAGGAA | AUUCGCCCUCCAUUUUUAUGG-UACUACUCAGGUUGG--------------ACCAGAGA | 5956 |
| 9 | Picornav_R35 | AUAUACG | GUCAGG | UCACUGC--UGCUUUUUAGC-UUACUACUCAGGCGCA--------------GCCAGAGAA | 5845 |
| 10 | Picornav_S64 | GCAUAUACG | GACAGGAA | AGCCCC--UGCUUUUUAGC-UUACUACUCAGGCGGA--------------GCCAGAGAA | 5901 |
| 11 | Sanxia 12 | GUAUAUACGG | GUCAGGAA | UGCUCC--UGCUUUUUAGC-UUACUACUCAGAUGGA--------------AUCAGAGAAU | 5667 |
| 12 | Hangzhou-6 | AUACACGCG | GUCAGG | CAUUCC--GUGGUUCUGCC-UUACUACUCAGGGCCGGUU----------GCCCAGAGAA | 4807 |
| 13 | Colobanthus | AUAUACG | GACAGG | GUUGAACCGUAGCUUUUCAGCUUUACUACUCAGGGUCGGU-----------ACCCAGAGAA | 6067 |
| 14 | Striga Sa2 | AAAUACG | GACAGG | AGAAACGUGGCUUUUUAGCCUUACUACUCAGGACCGUUU----------GUCCAGAGAA | 4289 |
| 15 | A.halleri 1 | ACACAUACG | GACAGG | AAGAAACGUGGCUUUUUAGCCUUACUACUCAGGGCCGUUU----------GUCCAGAGAA | 5886 |
| 16 | Changjiang 10 | AAAUACG | GACAGG | CAGGAAACGUGGCUUUUUAGCCUUACUACUCAGGGUCGUUU----------ACCCAGAGAA | 5388 |
| 17 | Picornavir_H1 | AAAUACG | GACAGG | AGGAAACGUAGCUUUUUAGCUUUACUACUCAGGGCCGUUU----------GUCCAGAGAAA | 5590 |
| 18 | Dicis. Rpf114 | UGCAUAUACG | GACAGGGC | AGGAAGCGUGGCUUUUUAGCCUUACUACUCAGGGCCGUAA----------GUCCAGAGAA | 5376 |
| 19 | Dicistrov_AliP | AUAUACG | GACAGG | GGUAGCGUAGCUUUUUAGCUUUACUACUCAGGACCGUAA--------- GUCCAGAGAA | 5376 |
| 20 | Crane70 | ACAUACG | GACAGG | GUCAACGUGGUUUUUUAACCUUACUACUCAGGGCCGUUU----------GCCCAGAGAAA | 5470 |
| 21 | Pico172-k141 | AAAUACG | GACAGG | GGUACACCUUGGCUAUUUAGCCUUACUACUCAGGGCGGGUUAA--------GUCCAGAGAU | 6857 |
| 22 | Pico351-k141 | CAUAAUAUACG | GUCAGGAA | CUUACCGUACAUUUUCAUGUUUACUACUCAGGGCCGGUUUU--------GCCCAGAGA | 5511 |
| 23 | L. sativa 151 | UGCAAAUACG | GACAGGGC | GUAUACCCGGGCUGUUUAGCCUUACUACUCAGGACGGGUUAA--------GUCCAGAGAU | 6814 |
| 24 | Hydrochara | AUAUACG | GACAGG | AGAACUGUAGCUUUUUAGCUUUACUACUCAGGGUCAGUAUA--------ACCCAGAGAA | 739 |
| 25 | Hubei 20 | AUAUACG | GACAGG | UAGUAGCGUAGCUAUUUAGUUUUACUACUCAGGGCCGCUUUA-------CGUCCAGAGAA | 5525 |
| 26 | Changjiang_11 | AUAUACG | GACAGG | UUAUAUAAUAAGAUGCGCUUUUUAGCGUUACUAGUCAGGAUGUCUUUGAA------AUCCAGACGA | 5288 |
| 27 | E.sinensis-1 | AUAUACG | GACAGG | UCAUUUAAAGAGAUGCAUUAUUUAAUGUUACUAGUCAGGAUGUCUCUAA------UAUCCAGACAC | 5433 |
| 28 | Weivirus-cra070shi3 | AAAUACG | GACAGG | UGGUAGCGUAGCUUUUUAGCUUUACUACUCAGGACCGCUAUAAA-----CGUCCAGAGAAAC | 5716 |
| 29 | Gingko pt112pil6 | AAAUACG | GACAGG | UAAGCACCAUUGGCUUUUUAGCCUUACUACUCAGGCGUGGUGUGAG-----UUGCCAGAGAAGAU | 6323 |
| 30 | Sanxia atyid V | AAAUACG | GACAGG | UGGACUCGUAACUAUUUAGUUUUACCACCUAGGACCGAGUUUAAA---UUGUCCAGGGAA | 5865 |
| 31 | Lactuca_pil8 | AAAGACG | GACAGG | UGGUUCCUGGCCUGUUUAGGCUUACCACCUAGGAAGGGAAGUAAAAUAAUUUCCAGGGAA | 5933 |
| 32 | Lactuca_pil9 | AAAUACG | GACAGG | UGGUUCCCGGCCUUUUUAGGCUUACCACCCAGGAAGGGAAGUGAAACAAUUUCCAGGGAA | 6203 |
| 33 | Gingko_-pt112-pil7 | UGCAAAUACG | GACAGGGC | UGGUUCCCGGCCUUUUUAGGCUUACCACCCAGGAAGGGAAGUGAAACAAUUUCCAGGGAA | 6322 |

**Table S3. Sequence alignments of L1.1a and L1.1.b loops and domain 2 of type 6d IRESs.**

Aligned sequences are from (1) Nedicistrovirus (JQ898341.1), (2) Antarctic picorna-like virus 1 (KM259869.1), (3) TSA: Pallasea cancelloides c22367_g2_i1 (GEQX01016691.1), (4) MAG: Chemarfal virus 33 isolate ML176 (OR270236.1), (5) Changjiang picorna-like virus 9 strain CJLX26226 (KX884541.1), (6) Beihai picorna-like virus 78 strain BHBei73249 (KX883307.1), (7) Gingko biloba dicistrovirus strain pt112-dic-11 (MN729613.1), (8) MAG: Hangzhou dicistro-like virus 4 isolate gl953 (OQ363017.1), (9) MAG: Picornavirales sp. isolate s64-k141_2464283 (MZ678982.1), (10) MAG: Picornavirales sp. isolate R35-k141_316374 (MZ678988.1), (11) Sanxia picorna-like virus 12 strain SXXX24153 (NC_032403.1), (12) MAG: Hangzhou dicistro-like virus 6 isolate gl235 (OQ363019.1), (13) TPA_asm: Colobanthus quitensis dicistro-like virus 2 isolate Cq2 (BK059249.1), (14) Striga asiatica dicistro-like virus 2 isolate Sa2 (MZ598484.1), (15) TPA_asm: Arabidopsis halleri dicistro-like virus 1 isolate Ah1 (BK059246.1), (16) Changjiang picorna-like virus 10 strain CJLX30646 (NC_032823.1), (16) MAG: Picornavirales sp. isolate H1_Bulk_30_scaffold_14 (MN035381.1), (18) Dicistroviridae sp. isolate Rpf114PicorV02-8 (MZ556256.1), (19) Dicistroviridae sp. isolate Rpf114PicorV02-8 (MZ556256.1), (20) MAG: Riboviria sp. isolate crane70_contig_3236 (OQ423883.1), (21) MAG: Picornaviridae sp. isolate 172-k141_140884 (ON162074.1), (22) MAG: Picornaviridae sp. isolate 351R-k141_276547_2 (ON162245.1), (23) Lactuca sativa dicistroviridae strain pt151-dic-14 (MN722412.1), (24) TSA: *Hydrochara caraboides* C113640_a_34_0_l_3466 (GDPR01004164.1), (25) Hubei picorna-like virus 20 strain WGML146806 (NC_033000.1), (26) Changjiang picorna-like virus 11 strain CJLX30672(NC_032833.1), (27) MAG: *Eriocheir sinensis* dicistrovirus 1 isolate Tangshan (OP019098.1), (28) MAG: Weivirus-like virus sp. isolate cra070shi3 (MT138207.1), (29) Ginkgo biloba dicistrovirus strain pt112-pil-6-plant 112 (MN841300.1) (Yang et al., 2022), (30) Sanxia atyid shrimp virus 3 strain SXXX37589 (NC_033228.1), (31) Lactuca sativa dicistroviridae strain pt151-pil-8-plant 151 (MN841302.1), (32) Lactuca sativa dicistroviridae strain pt151-pil-9-plant 151 (MN841303.1) and (33) Ginkgo biloba dicistrovirus strain pt112-pil-7-plant 112 (MN841301.1). Sequences that shared 98-100% identity with the listed IRESs were excluded from further analysis. These sequences are annotated to show the 3'-terminal nucleotide number of domain 2, structural elements (labelled as in Fig. 4A), and conserved nucleotides (in bold).

Table S4. Sequence alignments of L1.1a and L1.1.b loops and domain 2 of a subset of type 6a IRESs.

| # | Virus | Loop L1.1a | Loop L1.1a | Domain 2<br>P2.1 P2.3<br>P2.2/L2.1 P2.4/L2.3 | |
|---|---|---|---|---|---|
| 1 | Riboviria (Crane70)* | GUCU | UUCGCG | CCUGGACUAUUUAGUCUUACUACUCAGGAUGGAAGGUGGCAGCCCCACCAAUAUCCAGAGUC | 410 |
| *2* | *S. fonscolombii* | UGAUCUU | UUAUGC | AAGGGUGUUGAAUAUUUAUUCUUACUACUCAAGAUGCACC-GGAGCAGCCCUCC-AAUAUCUAGAGAAC | 168 |
| 3 | Guiyang dicistrov 2 | UGAUCU | CUGCU | AAGGGGGUUUGUUAUUUAACAUUACUUCUCAGGAUGCCCC-GUGGCAGCCCCAC-AAUAUCCAGAGAAC | 5804 |
| 4 | A.gracilipes2 | GUGAUC | UGCU | UGGAAGAACGCUACCUUUUAAGGUUUACCACUCAGGAUGCGUU-GGUGCAGCCCACU-AAUAUCCAGAGAA | 4291 |
| 5 | Dicistro_Rpf101 | GUGAUCGC | AAGUGCU | GAGAAGGACGUUAGUUAUUUAGCUUUACCACUCAGGAUGCGUC-AGUGCAGCCCACU-AAAAUCCAGAGAA | 6489 |
| 6 | TriatR119-k141 | GAUCGC | AAGUGCU | AAGAAGAGCGUUAGUUAUUAAACUUUACCACUCAGGAUGCGCU-GGUGCAGCCCACU-AAUAUCCAGAGAA | 6535 |
| 7 | M. scabrinodis V2 | GUGAUCU | GUGCU | AUAAGAAGAGCGUUAGAUAUUUAUCUUUACCACUCAUGAUGCGCU-GGUGCAGCCCACC-UAUAUCAAGAGAA | 6324 |
| 8 | Triato_s81-k141 | GUGAUCU | GUGCU | AUAUGAUCCUUAUCUAUUUAGAUUUACUGCUCAAGAUGGAU-GAGGCAGCCCCUC-AAUAUCUAGAGCA | 6022 |
| 9 | Dicistro_mointe135 | GUGAUCU | GUGCU | UCAUAUGAUCCUUAUCUAUUUAGAUUUACUGCUCAAGAUGGAU-GAGGCAGCCCCUC-AAUAUCUAGAGCA | 6043 |
| *10* | *U. diversus* | UGAUCU | GUGC | AAGAUCCUUACCUAUUUAGGUUUACUGCUCAAGAUGGAU-GUGGCAGCCCCAC-AAUAUCUAGAGCAC | 2544 |
| *11* | *Araneidae* IDV2115* | GUGAUCUGC | CAGUGCU | AGGAAGAGCGUUAGUUAUUUAACUUUACCACUCAGGAUGCGCU-GGUGCAGCCCACU-AAUAUCCAGAGAA | 4038 |
| *12* | *T. mandibulata** | GUGAUCGC | AAGUGCU | AAGAGCGUUAGUUAUUUAACUUUACCACUCAGGAUGCGCU-GGUGCAGUCCACC-AAUAUCCAGAGAAC | 1524 |
| *13* | *E. sakiedaorum* * | UGUGAUCU | UUGCUG | AAGGCUCUUAACUAUUUAGUUUUAAUGCUCAGGAUGGAGC-GUGGCAGUCCCAC-AAUAUCCAGAGCAC | 146 |
| 14 | Soybean Thrips | GUGAU | AGUGCU | AGGAAGAGCGUUAGAUAUUUAUCUUUACCACUCAAGAUGCGCU-GAGGCAGCCCCUC-AAUAUCUAGAGAA | 3793 |
| *15* | *T.mandibulata** | UGAUCGC | AAGUGC | AAGAGCGUUAGUUAUUUAACUUUACCACUCAGGAUGCGCU-GGUGCAGUCCACC-AAUAUCCAGAGAA | 1524 |
| *16* | *C. laticauda** | UGAUCC | ACGC | GUGCGUAGCUAUUUAGCUUUACUACUCAGGAUCGCACGUUGGCAGCCCCAGCAAAAUCCAGGG | 3171 |
| 17 | Bat dicistrov 4 | UGAUCG | ACGC | AAGACCGUAGCUAUUUAGCUUUACUGCUCAGGGUCGGUCGUUGGCAGCCCCAACAAUACCCAGAAC | 7208 |
| 18 | Dicis.R120-k141 | UGAUCU | CUGC | AAUUAGGUUAGCUAUUUAGCUUUACCACCCAGGAUGCCUA-GUGGCAGCCCCAC-AAUAUCCAGGGUAC | 6242 |
| *19* | *K. oculiprominens** | UGAUC | UGCU | UAUGGCUGGGCUAUUUAGCCUUACCUCCCAGGAUGGCCA-GUGGCAGCCCCAC-AUUAUCCAGGGAAC | 4230 |
| 20 | Dicistro_wpk049 | UGAUCU | GUGC | CAUGACCUGCGCUAUUUAGCGUUACCUCCCAGGAUGGGUC-GUGGCAGCCCCAC-AAUAUCCAGGGAA | 5883 |
| 21 | Triatov_77-k141 | UGAUC | UGC | GAGGACGUGCGCUAUUUAGCGUUACCACCCAGGAUGCGUC-GUGGCAGCCCCAC-AAUAUCCAGGGUA | 6424 |
| 22 | Wuhan 2 | UGAUCU | GUGC | AGUUCCCUUAGCUAUUUAGCUUUACGACCCAGGAUGGGGA-GAGGCAGCCCCUC-AAUAUCCAGGGUA | 6071 |
| 23 | Guiyang1 | UGAUCU | UUGCU | GGACAUCCUGCGCUAUUUAGCGUUACCACCCAGGAUGGGAU-GUGGCAGCCCCGC-AAUAUCCAGGGUA | 6071 |
| 24 | Bat dic_Mm/2010 | UGAUCU | CUGC | CAUGAGGCUC-GGACUAUUUAGUCUUACCUCCCAGGAUGAGCC-GUGGCAGCCCCAC-AAUAUCCAGGGAA | 6091 |
| 25 | Dicistro_rfb093dic1 | GUGAUCU | GUGCU | AUAAGAUGUUAGUUAUUUAACUUUACUGCUUAGGAUGCAUC-GUGGCAGCCCCAUAAAAAUCCAGAGCC | 6226 |
| 26 | R. padi virus | UGAUCU | UUGC | UGGUUAGCUAUUUAGCUUUACUAAUCAAGACGCCGUC-GUGCAGCCCAC-AAAAGUCUAGAUAC | 7058 |
| 27 | Dicistro_coa195dic1 | UGAUCU | UUGC | ACACUCGGUUAGCUAUUUAGCUUUACUAAUCAAGACGCCAUC-GUGCAGCCCAC-AAAAGUCUAGAUA | 7046 |
| 28 | Woodpecker | UGAUCU | AUGC | GUUUGGCUGGGUUAUUUAACCUUACCUUCCAGGAUGGCCA-GUGGCAGCCCCAC-AUUAUCCAGGGAA | 6262 |
| 29 | Flumine20 | UGAUC | UGC | UAUUUAGGUUAGCUAUUUAGCUUUACGUUCCAGGAUGCCUA-GUGGCAGCCCCAC-AAUAUCCAGGAAAC | 2325 |
| 30 | A.gracilepes 1 | UGAUCU | UUGC | UUAUCCC-UACCUAUUUAGGUUUACGUUCCAGGAUGGAU-GUGGCAGCCCCAC-AAUAUCCAGGAA | 6152 |
| 31 | Nelson Picornalike | UGAUCU | UUGC | UUUUCCUUUGCUAUUUAGCAUUACGUUCCAAGAUGGGAA-GAGGCAGCCCCUC-AAUAUCUAGGAAA | 5849 |
| 32 | Dicistro_R119-k141 | | | UAAUGAGGUUAACUAUUUAGUUUUACU-UCCAGGAUGCCUCAAUAGCAGCCCUAUUUAAAUCCAGGAAC | 4674 |
| 33 | Flumine dicistrovirus 1 | UGAUCG | ACGC | AGUAGGACCGUAGCUAUUUAGCUUUACUGAUCAGGGUCGGUCGUUGGCAGCCCCAACUAUACCCAGAA | 7072 |
| 34 | Bundaberg bee V1 | UGAUCG | ACGC | GUAAUGACCGUAGCUAUUUAGCUUUACUGCUCAGGACCGGUCGGUGGCAGCCCCACUAAAGUCCAGAA | 7127 |
| | | | | S2.1 S2.2 L2.1 L2.2 L2.3 L2.4 S2 . 3 | |

**Table S4. Sequence alignments of L1.1a and L1.1.b loops and domain 2 of a subset of type 6a IRESs containing UUACUA and related motifs in the L2.2 loop of PKIII.**

Sequences are annotated to show the 3'-terminal nucleotide number of domain 2, structural elements (labelled as in Fig. 5A), and conserved nucleotides (in bold). Aligned sequences are from (1) MAG: Riboviria sp. isolate crane70_contig_262 (GenBank: OQ423838.1), (2) TSA: *Sympetrum fonscolombii* breed wildtype C86879_a_27_0_l_4049 (GCLN01018907.1),(3) MAG: Guiyang dicistrovirus 2 isolate YZZGZ282461 (MZ209774.1), (4) *Anoplolepis gracilipes* virus 2 (MT108240.1), (5) Dicistroviridae sp. isolate Rpf101dic04-8 (MZ556249.1),(6) MAG: Triatovirus sp. isolate R119-k141_5185 (MZ679098.1), (7) *Myrmica scabrinodis* virus 2 (MH477289.1), (8) MAG: Triatovirus sp. isolate s81-k141_262366 (MZ679080.1), (9) Dicistroviridae sp. isolate mointe135dic01-8 (MZ556258.1), (10) TSA: *Uloborus diversus* IDV#2099 mRNA, IDV2099-W1_S220034389 (IAVI01034389.1), (11) TSA: *Araneidae* gen. sp. IDV 2115 IDV#2115 (ICNA01035142.1), (12) TSA: *Tetragnatha mandibulata* IDV#62 mRNA, IDV62-W1_S110014326 (IAJA01014326.1), (13) TSA: *Eriovixia sakiedaorum* IDV#5652 mRNA, IDV5652-W1_S100011448 (IBEB01011448.1), (14) Soybean thrips picorna-like virus 4 strain STN1 (MT240799.1) (15) TSA: *Tetragnatha mandibulata* IDV#62 mRNA, IDV62-W1_S110014326 (IAJA01014326.1), (16) TSA: *Cyclosa laticauda* IDV#1246 mRNA, IDV1246-W1_S160021638 (IAWK01021638.1), (17) MAG: Bat faecal associated dicistrovirus 4 isolate CP02/aus/2 (ON872534.1), (18) MAG: Cripavirus sp. isolate R120-k141_301338 (MZ679083.1), (19) *TSA: Keijiella oculiprominens* IDV#4340 mRNA, IDV4340-W1_S150031619 (IANK01031619.1), (20) MAG: Dicistroviridae sp. isolate wpk049shi01 (MN917670.1), (21) MAG: Triatovirus sp. isolate 77-k141_427369 (MZ679090.1), (22) Wuhan arthropod virus 2 strain WHSF0 (NC_033437.1), (23) MAG: *Guiyang argiope* bruennichi dicistrovirus 1 isolate ZZGZLJ150573 (MZ209806.1), (24) Bat dicistrovirus strain Mm/2010 (MH370347.1), (25) MAG: Dicistroviridae sp. isolate rfb093dic1 (MN918755.1), (26) *Rhopalosiphum padi* virus (NC_001874.1), (27) MAG: Dicistroviridae sp. isolate coa195dic1 (MN918733.1), (28) MAG: Riboviria sp. isolate woodpecker139_contig_5 (OQ425164.1), (29) Flumine dicistrovirus 20 (OM953863.1), (30) *Anoplolepis gracilipes* virus 1 (MT108239.1), (31) Nelson Picorna-like virus 6 isolate Vvul_virus_15 (MZ443583.1), (32) MAG: Dicistroviridae sp. isolate R119-k141_127746 (MZ679342.1), (33) Flumine dicistrovirus 1 (OM953859.1) and (34) Bundaberg bee virus 1 isolate QLD-6 (MG995701.1).

Table S5. Sequence and structural alignment of type 6b IRESs.

| # | Viral sequence | | P1.1 P1.2 P1.3 |
|---|---|---|---|
| 1 | Beihai4 | 6799 | AAUGUGAGUUUUGGUUACUCAUCGCAGGUGAACUACGCUCCAGACAACUAACCUGCG--UAUAGCUUGCUCACAUU |
| 2 | Beihai93 | 7018 | --GGUACACAUUGGUUACGCAACGGGCA-UAACUACUCCUCAGACAACUACUGUCCG--UAUCGCUUUGUGUACU |
| 3 | Wenzhou7 | 6637 | -GCCUCGUGAUUGGUUA-CCCAUCACUGCAAAUUACUCUCCAGACGGCUGUGCAGUG--UAAGGCUUUCACGAGGU |
| 4 | E. sinensis | 742 | --UCUUGUGUUUGGUUA-CCCAUCACUGUUAAUUACGCUCCAGAUGACUAUGCAGUG--UACGGCUUUCACAAGA |
| 5 | Mudcrab | 7022 | -UCUCCAUGUCUGGUUA-CCUAUCUAUCUUAACUACGCUCCAGACGACUAAAUUUAG--UAUGGCUUUCAUGGAGA |
| 6 | Taura | 6761 | -CUCCAUGUAUUGGUUA-CCCAUCUGCAUCGAAAA-CUCUCCGAACACUAGGUGCAG--UAAGGCUUUCAUGGAG |
| 7 | O. oratoria | 13 | -CCUUGUGUAUUGGUUA-CCCAUCCAUGCUAACUACGCUCCAGACAACUAGGCAUGG--UACGGCUUUCACAAGG |
| 8 | Beihai5 | 6848 | -CUCUACAUACUGGAUA-CCCAUCUAAACUAACUACGCUCCAGACGACUAAAUUUAG--UAAGGCUUAUGUAGAG |
| 9 | Macrobrachium | 6891 | -UUCUAUAUAUUGGAUA-CCCAUCUAAAUUAACUACGCUCCAGACGACUAAAUUUAG--UAAGGCUUAUAUAGGA |
| 10 | Z. caudelli | 2989 | GUGUAUAACUCUGGUUA-CCGUAUAUGUUAAGAAUUUGUUAGUUUAAUUUUCAUAUAU--UAGGCUUUGUUAUA |
| 11 | Zoanthus | 3397 | ---UCACUACCUGGUUA-CC-AAGUAACACUCAUAAACCGGUUGUAAAAGGUGUUACU--CAGGCUUGUAGUGA |
| 12 | C. maenas | 5846 | --CUACUACUCUGGUUA-CCCAACUUGUCAUAUUACUGAUCAAACAUAAAUGACGAG-CCAUAGCUUUGUAGUAG |
| 13 | D. gallinae | 3072 | ---UUAGUGCGUGAUC--CCUCCCUGUUCAUAAGUAUAAGUGUCGAAAGAACAGG---UGCACGCCGCACGAA |
| 14 | Pycnopodia | 6424 | --CUCAUUGUGUGAUC----UUGCUUGUAACAAUUACUUCCAAUGCACUAGUUACU-UGCACUGCUACAAUGAG |
| 15 | Behai91 | 6126 | ----UGGAUGUGAUC----UUGUAUGCAUAUAAUGUAGUCUAGUAAAUUGCAUAC----ACUGCUAUCCA |
| 16 | Leptomastix | 1015 | --AGCAGAAUGUGAUC----UU--GCUCGAAUAUA---UUGAUACGAAAUUUCGAGU---AUUGCUAUUCAACU |
| 17 | M.occidentalis | 320 | ---UGGAUGUGAUC------UUGCUACUUUAUA--UUGAAGAGAAAUAAGUAGU----AAUGCUAUCCA |
| 18 | C. omonaga | 2007 | ---CAAAUCUCUGGUUA-CC-AGCUAAUUC-UAAUAACUGACCUGAAAUGAAUUAGU---AAGGCUUUGAUUUG |
| 19 | P. keikoae | 3173 | -----CUUUUUAUGUGA-UCCCUCUUCUUCCAAACAUUGAAAUUUAAUAGGAUAGAAUGCAC-GCUAUAAAAAG |
| 20 | Z. potanini | 598 | --AUAGCUCUAAUGUGA-UCCCUCCUCUUCCGAACCUUGAAACUAAAUAGGAUAGAGUGUAC-GCUGUUAGAGU |
| 21 | Photinus | 4597 | -GUCUUAUUAUUGGUUA-CC-AUGUGUUAAUUAUAA--GUUUCCUCAAGAUUAAC---UAAGGCUUUAGUAAGAU |
| 22 | A. potteri | 1831 | -CGUAUUGUAUUGGUUA-CCCAUCAACUUAAUAAUU--UGUCUUAAA-UAAGUUG----UAAGGCUUGCAAUACG |
| 23 | Hubei25 | 5996 | ACUCUCUUAUUUGGUUA-CCGAAUUUUCUAAUAAA-CACCUCGAAUAUGAAAAUU----UUAGGCUUAUAAGAGU |
| 24 | C. osculatum | 355 | -GCUUAGAUAUUGGUUA-CCGAUGAAAUUUCGUAA-CUGAUCACAAGAAAUUUCAU---UA-GGCUUAUCUAAGU |
| 25 | M. chinense | 718 | -GCAAUUUCACUGGUUA-CCCAUGCAAUCGAGAAUUUAGGUGAAUAAAUUGAUUGU---UAAGGCUUUGAAAUUGU |
| 26 | Araneidae | 3518 | -CUAAAAGUAUUGGUUA-CCCAUUAUGACAGGAAAUUUCUGUUGAACUAGUUAUA----UAAGGCUUACUUUUAG |
| 26 | Argiope | 3458 | -CAACACAACAUGGUUA-CCCAUAAAUUGAGGAAAUUUCCAAUAAACUCAAUUU-----UAAGGCUUGUUGUGUUG |
| 28 | A. bruennichi | 2132 | -CUUUCUCUUAUGGUUA-CCCAUCAUUUGAGGAAAUUUCCAAUAAACUCAAAUG-----UAAGGCUUAGAGAAAG |
| 29 | ABPV-144I | 6331 | -CAACACAACAUGGUUA-CCCAUAAAUUGAGGAAAUUUCCAAUAAACUCAAUUU-----UAAGGCUUGUUGUGUUG |
| 30 | Dicistro_bsk | 6172 | -CUUUCUCUUAUGGUUA-CCCAUCAUUUGAGGAAAUUUUCAAUAAACUCAAAUG-----UAAGGCUUAGAGAAAG |
| 31 | F. pressilabris | 33 | -CUUUCUCUUAUGGUUA-CCCAUCAUUUGAGGAAAUUUCCAAUAAACUCAAAUG-----UAAGGCUUAGAGAAAG |
| 32 | Dicistro_jyt032 | 6851 | -CUUUCUCUUAUGGUUA-CCCAUCAUUUGAGGAAAUUUCCAAUAAACUCAAAUG-----UAAGGCUUAGAGAAAG |
| 33 | KBV | 6428 | -CACUGUUUCAUGGUUA-CCCAUCAUUUGAGGAAAUUUCCAAUAAACUUGAAUG-----UAAGGCUUGAAACAGUG |
| 34 | Pogonomyrmex | 3101 | -CGUUCAUUCAUGGUCA-CCCAUCACUUGAGGAAAUUUCCAAUAAACUUAAGUG-----UAAGGCUUGAAUGAACG |
| 35 | Fex1 | 6431 | -CAUCUAUUCAUGGUCA-CCCAUCACUUGAGGAAAUUUCCAAUAAACUUAAGUG-----UAAGGCUUGAAUAGAUG |
| 36 | SN11649 | 6439 | -CAUCUAUUCAUGGUCA-CCCGUCACUUGAGGAAAUUUCCAAUAAACUUGAGUG-----UAAGGCUUGAAUAGAUG |
| 37 | IAPV | 6493 | -CAUCAUUCUAUGGUUA-CCCAUCAUUAGAGGAAAUUUCCAAUAAACUCUGGUG-----UAAGGCUUAGAGUGAUG |
| 38 | SIV5 | 6267 | -CAUUCUUUCAUGGUCA-CCCAUCACAUAAGGAAAUUUCCAAUAAACUUGUGUG-----UAAGGCUUGAAAGAAUG |
| 39 | O. navus | 573 | -CAUCUUCUCAUGGUUA-CCCAUCAUUCAAGGAAAUUUACAAUAAACUUGAAUG-----UAAGGCUUAAGAAGCUG |
| 40 | C. monticola | 802 | -CAUCUUUUCAUGGUCA-CCCAUCAUUCAAGGAAAUUUCCAAUAAACUUGAAUG-----UAAGGCUUGAGAAGAUG |
| 41 | D. umbrophila | 5242 | -CAUCUUCUCAUGGUUA-CCCAUCAUUCAAGGAAAUUUCCAAUAAACUUGAAUG-----UAAGGCUUGAGAAGAUG |
| 42 | Apara114-k141 | 6530 | -CGUUCUUCUAUGGUUA-CCCAUCACUUGAGGAAAUUUCCAAUAAACUUGAGUG-----UAAGGCUUAGAAGAAUG |
| 43 | C. holgerseni | 3129 | -CAUUCUUUCAUGGUUA-CCCAUCACUCAAGGAAAUUUCCAAUAAACUUGAGUG-----UAAGGCUUGGAAGAAUG |
| 44 | Novo_mesto | 6524 | UCUAAACUUUCUGGUUA-CCUAGUGGAU-AGGAAAUUGUCAAUUAACUACAUCCAU---GAUGGCUUUAAGUUUAGG |
| 45 | M. rubra | 6510 | UCUAAACUUUCUGGUUA-CCCAAUGAAC-AGGAAAUUGUCAAUUAACUAGUUUCAU---GAUGGCUUUAAGUUUAGG |
| 46 | M. japonica | 3000 | UCUAAACUUUCUGGUUA-CCUAGUGAAC-AGGAAAUUGUCAAUUAACUAGCUUCAU---GAUGGCUUUAAGUUUAGG |
| 47 | A. stella | 1135 | -CUAAACUUUCUGGUUA-CCUAGUGAAC-AGGAAAUUGUCAAUUAACUAGUUUCAU---GAUGGCUUUAAGUUUAG |
| 48 | H. azumiensis | 2112 | -CUAAACUUUCUGGUUA-CCUAGUGAAC-AGGAAAUUGUCAAUUAACUAGUUUCAU---GAUGGCUUUAAGUUUAG |
| 49 | M. chinense2 | 3716 | -CUACUAAUUUUGGUUA-CCUAGGAAAU-AGGAAAAUGUCUUUUAACUAUAUUUCU---GAUGGCUUUAUUAGUAG |
| 50 | W. contortipes | 257 | -GAGAUUUUUCUGGUUA-CCUAAGAAAU-AGGAAAUUGUCAAUAAACUACAUUUCU---GAUGGCUUUAAAAUCUU |
| 51 | E. cambridgei | 6479 | -GAGAUUUUUCUGGUUA-CCUAGGAAAU-AGGAAAUUGUCAAUAAACUACAUUUCU---GAUGGCUUUAAAAUCUU |
| 52 | L. humile | 6454 | -CUAAUUUCUCUGGUUA-CCCAGCUCAU-AGGAAAAUGUCUUUUAACUA-UUGAGU---CAAGGCUUAGAAAUUAG |
| 53 | H.yanbaruensis | 887 | -GUCUCUAUUCUGGUUA-CCCGCUCAGU-AGGAAAUUUCCAUUGAACUACACUGAG---UAAGGCUUUAUAGAGAU |
| 54 | SIV1 | 4223 | --UUUCUCUUCUGGUUA-CCAGCAACAU-AGGAAAUUGUCGUUGAACUACAUGUUG---UAAGGCUUUAGAGAAAU |
| 55 | M. illinoiensis | 276 | --UUUCUGUUCUGGUUA-CCCAUCUAUCGAGGAAAAUGUCGUUUAACUAUGAUAG----UAAGGCUUUACAGAAAU |
| 56 | SIV12 | 5893 | -CUAAAAUCUAUGGUUA-CCCAACCUAA-AGGAAAUUUCCUAUAAACUUUAAGGC----CAAAGCUUAGAUUUUAG |
| 57 | P. tepidariorum | 3437 | -UUAGAACUUCUGGUUA-CCCAGCCUAGA-GGAAAUUUCCAAUAAACUUUAAGGU----CAAGGCUUUAGUUCUAA |
| 58 | Hubei1 | 5938 | -CUAGAACUUCUGGUUA-CCCAGCUUAAA-GGAAAUUUCCAGUAAACUUUAAAGU----CAAGGCUUUAGUUCUAG |
| 59 | P. cryptcolens | 1437 | -UUAGAACUUCUGGUUA-CCCAGCUUAAA-GGAAAUUUCCAAUAAACUUUAAAGU----CAAGGCUUUAGUUCUAA |
| 60 | Holoplatys | 4264 | -CACUAACUUCUGGUUA-CCCAGCCAAUA-GGAAAAUGUCUUUUAACUAAUUGGU----CAAGGCUUAAGUUAGUG |
| 61 | A. yesoensis | 6259 | -CAAUUCUUUCUGGUUA-CCUAGUAGAUA-GGAAAUUUCCAAUUAACUACAUCUAU---GAUGGCUUUAAGAAUUG |
| 62 | Haeterina | 29 | -CAUCUCUUUCUGGUUA-CCUAGGGAGUA-GGAAAUUGCCAAUUAACUACACUCCU---GAUGGCUUUGAGAGAUG |
| 63 | P. labialis | 1549 | -GCAUCUAUCCUGGUUA-CCUAGGGAGUA-GGAAAUUGCCAAUUAACUACACUCCU---GAAGGCUUUAUAGAUGU |
| 64 | Wuhan11 | 6107 | -AUUCUCUUUCUGGUUA-CCUAACGAACA-GGAAAUUGCCAAUUAACUAGUUUCGU---GAUGGCUUUAAGAGAAU |
| 65 | E. nubilus | 3927 | --ACUUCUUUCUGGUUA-CCUAGUGAGUA-GGAAAUUGCCAAUUAACUACACUCAU---GAUGGCUUUAAGAAGUG |
| | | | L1.1a L1.2a. L1.2b L1.2a |

| # | Viral sequence | P1.3 / P2.1 / P2.2/L2.1 / P2.3 / P2.4/L2.3 |
|---|---|---|
| 1 | Beihai4 | -------GGUUUGCUUUUUAGCGUGUGUGUACCA---GAGGCAGUCCCUUA-UACACAUGUCAGGAG |
| 2 | Beihai93 | -----UUGGCUUGCUAUUUAGCGUGCGUGUGCCUU---GGGCAGCCCCCUA-AACACGUGUCAGAGG |
| 3 | Wenzhou7 | -------GGCUUGCUUUUCAGCGUACGUGUGCCAC---AGGCAGCCCCGAA-AACACGCGUCAGGAG |
| 4 | E. sinensis | ------CGGCUUGCUUUUCAGCGUACGUGUGCCAC---AGGCAGCCCCGAA-AACACGCGUCAGGAG |
| 5 | Mudcrab | -------GGUUUGCUAUUCAGCGUGCGUGUACCAU---CGGCAGCCCCAAA-AACACGCGUCAGGAG |
| 6 | Taura | ------UGGUUUGCUAUUUAGCGUACGUGUACCAU---AGGCAGCCCCAAA-AACACGUGUGAGGAGA |
| 7 | O. oratoria | ------UGGUUUGCUAUUUAGCGUGCGUGUACCUU---AGGCAGCCCCGAA-AACACGUGUCAGGAG |
| 8 | Beihai5 | ------UGGUUUGUUAUUUAACGUGCGUGUACCCU---AGGCAGCCCCAAA-AACACGCGUCAGGAG |
| 9 | Macrobrachium | ------UGGUUUGUUAUUCAACGUGCGUGUACCCC---AGGCAGCCCCGAA-AACACGCGUCAGGAG |
| 10 | Z. caudelli | -UGUAAAGGUGUGAUAUUUAUCAUGAG-GUGCCUG--GUGGCAGCCCCAUUUAAACCUCAAAAUUAAC |
| 11 | Zoanthus | -UUUAAGAGCUUCCUAUUUAGGGUUGGUGAGCUC---GGAGCAGCCCUCUA-UUCACCACACAACC |
| 12 | C. maenas | ---AUCGUGGUUGCUAUUUAGCGUAGGUGUCCAC---GGAGCAGCCCUCCA-UACACCGUUAGAUCA |
| 13 | D. gallinae | -----CCGGCUUCCUAUUCAGGGUGAGUGUGCCA---GGAGCAGCCCUCCA-UACACUCAGACACU |
| 14 | Pycnopodia | --UUUUUGGGUUGCUAUUUAGCGAAGGUGUCCCA---GGAGCAGCCCUCCA-UACACCGCAUAGGAA |
| 15 | Behai91 | GCUUUUAGGUCCCCUAUUUAGGGAGAGCGUCCUG---CGAGCAGCCCUCCAAAA-GCUCAAUAGACU |
| 16 | Leptomastix | ----ACAGGUUCUCUAUUCAGAGAUGGCGUCCUA---GCAGCAGCCCUGCAAAA-GCCAAGUAUCAA |
| 17 | M.occidentalis | -AUUAAAGGUGCUCUUUUUAGAGUUGGCCUCCUC---GUGGCAGCCCCACAAAA-GCCAAUCUUCAA |
| 18 | C. omonaga | UUUAAGAGGGGAGCUAUUUAGCUCGGGAACCCCUC-GGUGGCAGCCCCACUAAAGACCCUAGGUUA |
| 19 | P. keikoae | --AUGGAGGUAUACUUUUUAGUAUGGUGAAACCUC-GGUGGCAGCCCCACUAAAUUCGCAAAAUUUCA |
| 20 | Z. potanini | --AAUAAGGUGUGUUAUUUAGCACGGUGAUACCUU-GGAGGCAGCCCCUCUCAAAUCGCAUAGUUUCA |
| 21 | Photinus | --AAUAUAGUAUUCUAUUUAGAAUUAAGAGGACUAUGGUAGCAGCCCUACUGAACUCUUA-AGGAA |
| 22 | A. potteri | ---AUAGGGUUCUCUAUUCAGAGAAGGACGCCUU---CUGGCAGCCCCAGUCAAGUCCUA-AAGACA |
| 23 | Hubei25 | -UAAAAGAGUAUCCUAUUUAGGAUGAGAAUGCUCU-GGUGGCAGCCCCACUAAAAUUCUCUGAAGGU |
| 24 | C. osculatum | --GUCGGAGUGCCCUAUUUAGGGUGGGAAGCUCC--AGUGGCAGCCCCACUAAAUUCCC-UUGAGC |
| 25 | M. chinense | --GAUGGAGUGCUCUAUUUAGAGUGGGAAGCUCC--ACAGGCAGACCCUGUAAAUUCCCUAUUCAUCU |
| 26 | Araneidae | --AAUGUGGUGUGGUUUUUACCAUGAGGAGCCAC--GUGUGCAGCCCACACAAAUCCUC-UAGCAGA |
| 26 | Argiope | --GACAAGGUGCCCUAUUUAGGGUGAGGAGCCUU--ACUGGCAGCCCCAGUAAAUCCUC-UAUUGGA |
| 28 | A. bruennichi | --GUCGAGGUGCCCUAUUUAGGGUGAGGAGCCUC--GCUGGCAGCCCCAGUGAAUCCUC-UAUUGGA |
| 29 | ABPV-144I | --GACAAGGUGCCCUAUUUAGGGUGAGGAGCCUU--ACUGGCAGCCCCAGUGAAUCCUC-UAUUGGA |
| 30 | Dicistro_bsk | --GUCAAGGUGCCCUAUUUAGGGUGAGGAGCCUU--GCUGGCAGCCCCAGUGAAUCCUC-UAUUGGA |
| 31 | F. pressilabris | --GUCGAGGUGCCCUAUUUAGGGUGAGGAGCCUC--GCUGGCAGCCCCAGCAAAUCCUC-CAUUGGA |
| 32 | Dicistro_jyt032 | --GUCGAGGUGCCCUAUUUAGGGUGAGGAGCCUC--GCUGGCAGCCCCAGCAAAUCCUC-CAUUGGA |
| 33 | KBV | --GUCUAGGUGCCCUAUUUAGGGUGAGGAGCCUA--GCUGGCAGCCCCAGUGAAUCCUC-UAUUGGA |
| 34 | Pogonomyrmex | --GUCAAGGUGCCCUAUUUAGGGUGAGGAGCCUU--ACUGGCAGCCCCAGUGAAUCCUC-UAUUGGA |
| 35 | Fex1 | --GUCGAGGUGCCCUAUUUAGGGUGAGGAGCCUC--GCUGGCAGCCCCAGCAAAUCCUC-UAUUGGA |
| 36 | SN11649 | --GUCGAGGUGCCCUAUUUAGGGUGAGGAGCCUC--G-UGGCAGCCCCAGUAAAUCCUC-UAUUGGA |
| 37 | IAPV | --GUCGAGGUGCCCUAUUUAGGGUGAGGAGCCUC--GGUGGCAGCCCCACCAAAUCCUC-UAUUGGA |
| 38 | SIV5 | --GUCGGAGUACUCUAUUUAGAGUGAGGAGCUCC--GCUGGCAGCCCCAGCAAAUCCUC-UAUUGGA |
| 39 | O. navus | --GUCUAGGUGCCCUAUUUAGGGUGAGGAUCCUAG--CUGGCAGCCCCAGGGAAUCCUC-UAUUGGA |
| 40 | C. monticola | --GUCUAGGUGCCCUAUUUAGGGUGAGGAGCCUA--GCUGGCAGCCCCAGUGAAUCCUC-UAUUGGA |
| 41 | D. umbrophila | --GUCUAGGUGCCCUAUUUAGGGUGAGGAGCCUA--GCUGGCAGCCCCAGUGAAUCCUC-UAUUGGA |
| 42 | Apara114-k141 | --GUCUAGGUGCCCUAUUCAGGGUGAGGAGCCUA--GCUGGCAGCCCCAGUGAAUCCUC-UAUUGGA |
| 43 | C. holgerseni | --GUCUAGGUGCCCUAUUUAGGGUGAGGAGCCUA-GC-UGGCAGCCCCAGUGAAUCCUC-UAUUGGA |
| 44 | Novo_mesto | ---AAAGAGGGCACUAUUUAGUGUGAGGUCCUCU-GG-UGGCAGCCCCACUCAAACCUC-UAUUGAC |
| 45 | M. rubra | ---AAAGAGGGCACUAUUUAGUGUGGGGUCCUCU-GG-CAGCAGCCCUGCCCAAACCCC-UAUUGAC |
| 46 | M. japonica | ---AAAGAGGGCACUAUUUAGUGUGGGGUCCUCU-GG-CAGCAGCCCUGCCAAAACCCC-UAUUGAC |
| 47 | A. stella | --GAAGAAGGGCACUAUUUAGUGUGGGGUCCUUC-AG-CAGCAGCCCUGCUAAAACCCC-UAUUGAC |
| 48 | H. azumiensis | --GAAAGAGGGCACUAUUUAGUGUGGGGUCCUCU-AG-CAGCAGCCCUGCUAAAACCCC-UAUUGAC |
| 49 | M. chinense2 | --AAUAGAGGGUCCUAUUUAGGAUGAGGUCCUCU-GA-AGGCAGCCCCUUUAAAACCUC-UAAAGAC |
| 50 | W. contortipes | --AAGAGAGGGUUCUAUUUAGAAUGAGGUCCUCU-AG-UGGCAGCCCCACCGAAACCUC-UAUUGAC |
| 51 | E. cambridgei | --AAGAGAGGGUUCUAUUUAGAAUGAGGUCCUCU-AG-UGGCAGCCCCACCAAAACCUC-UAUUGAC |
| 52 | L. humile | --AAUGGAGGGUCCUAUUUAGGAUGAGGUCCUCC-GG-UGGCAGCCCCACCAAAACCUC-UAAAGAC |
| 53 | H.yanbaruensis | --AUCGAGGUGUCCUAUUUAGGAUGAGGUGCCUC-GG-UGGCAGCCCCACCAAAACCUC-UAGUGGA |
| 54 | SIV1 | --AAGGGAGUGUCCUAUUUAGGAUGAGGUGCUCC-GG-UGGCAGCCCCACCAAAACCUC-UAGCGAC |
| 55 | M. illinoiensis | --AUCAGGGUGUCCUAUUUAGGAUGAGGUGCCCU-GG-UGGCAGCCCCACCAAAACCUC-UAGCGAC |
| 56 | SIV12 | --AACGAGGUGCCCUAUUUAGGGUGUGGAGCCUC-GC-UGGCAGCCCCAGCAAAUCCAC-AAUAGGA |
| 57 | P. tepidariorum | --GUCAAGGUGCCCUAUUUAGGGUGAGGAGCCUU-GG-UGGCAGCCCCACCGAAUCCUC-AAUUGGA |
| 58 | Hubei1 | --GUCGAGGUGCCCUAUUUAGGGUGAGGAGCCUC-GG-UGGCAGCCCCAUCAAAUCCUC-UAUUGGA |
| 59 | P. cryptcolens | --GUCGAGGUGUCCUAUUUAGGAUGAGGAGCCUC-AG-UGGCAGCCCCACUAAAUCCUC-CAUUGGA |
| 60 | Holoplatys | --AAUGAGGUGCCCUAUUUAGGGUGAGGAGCCUC-GG-UGGCAGCCCCACCAAAUCCUC-UAAAGAC |
| 61 | A. yesoensis | --AACAAGGUGCCCUAUUUAGGGUGAGGAGCCUU-GC-UGGCAGCCCCAGCUAAUCCUC-UAUUGGA |
| 62 | Haeterina | --AAUGAGGUGCUUUAUUUAGAGUGAGGAGCCUC-GU-UGGCAGCCCCAACAAAUCCUC-AAUUGGC |
| 63 | P. labialis | --AUGAGGGUGUCCUAUUUAGGAUGAGGAGCCCU-GG-UGGCAGCCCCACCAAAUCCUC-UAUUGGC |
| 64 | Wuhan11 | --AAGAGGGUGUCCUAUUUAGGAUGAGGAGCCCU-GG-UGGCAGCCCCACCAAAUCCUC-AAUUGGC |
| 65 | E. nubilus | --AAUAAGGUGCCCUAUUUAGGGUGAGGAGCCUU-GG-UGGCAGCCCCACTAAAUCCUC-AAUUGGC |

| # | Viral sequence | P3.1/P3.2 — P3.3 — P3.4/L3.3 |
|---|---|---|
| 1 | Beihai4 | -AAAUUCUCAGCCCACU-UGGGUCA-GUGUGGCAGCCACUUUUCUUAAAGUGGU----GAGAAUU-AAAAU------GACCUGCU 6999 |
| 2 | Beihai93 | -ACCGCUCCAGCCUAGU-UGGGUUA-CUACGGCAGCCACGCUUGCGUGGUGG-------AGUGU--AAAACU-----AACCUGCU 7213 |
| 3 | Wenzhou7 | -ACCUUUCCUGCUUUCU-UGGGCG-AGAACAGCAGCUGCGCUUCUUCGUGCAGU----GGAAAGUUUAAA------CUGCCUGCU 6836 |
| 4 | E. sinensis | -ACCUCUCCAGCUAUCU-UGGGCA-AGAUCAGCAGCUGCGCAGUUCUAUUGUGCAGU-GGAGAGUUUUUA------UUGCCUGCU 541 |
| 5 | Mudcrab | -AUCUCCCCAGCCACCA-UGGGCU-AGGUUGGCAGCCAGGUUUUAACUUGGU------GGGGAG-UAAAA------UAGCCUGCU 7218 |
| 6 | Taura | --AAGUCCCAGUCACUU-UGGGCA-AAGUAGACAGCCGCGCUUGCGUGGU--------GGGACUU--AAUUAA----UGCCUGCU 6955 |
| 7 | O. oratoria | -ACAGUUCCGGUCACUU-UGGGCA-GAGUAGACAGCCAUGCUUGCGUGGU--------GGAACU--UAUUA-----UCGCCUGCU 325 |
| 8 | Beihai5 | -ACAGUCUCAGUCACUU-UGGGCA-GAGUUGGCAGCCACGCUUGCGUGGU--------GAGACU--UAUC----UAUUGCCUGCU 7043 |
| 9 | Macrobrachium | -ACAGUUCCAACCACUU-UGGGCG-AAGUUGGUGGCUACGCUUGCGUAGU--------GGAACU--UAUU----UAUUGCCUGCU 7086 |
| 10 | Z. caudelli | UUUAAUUGCACUUAGGCGUGGGUUUGUCUUU--------------------------GUGUGU--AUAA----UUUGACCUGCU 2803 |
| 11 | Zoanthus | UCAUCCCCAGCC-AAUU-UGGGUC-AAUUCGGCAACUCUGGCCUAGCUAGAGU-----GGGGAU--CAAUU---UUCGACCUGCU 3198 |
| 12 | C. maenas | -ACGUCCCUGCA--CAC-UGGGUGGUGUGUAAAACAAGUGCUUGCGCUUG--------GGGAU--GAGAUA----CUCACCUGCU 6043 |
| 13 | D. gallinae | CGCGGCCCAGUC-ACAU-UAGGUC-AUGUUGACAGCCAGGAUUUCGAUUCUGGU----GGGCU--UUAAUA---CCCGACCUGCU 2878 |
| 14 | Pycnopodia | -CCUCCCCAGUC-ACAU-UGGGUU-AUGUUGACAGCUGCGUUUACGCAGU--------GGGG---UAGUAU---UACAACCUGCU 6618 |
| 15 | Behai91 | --UCGGAGCA-GCCAUCCUGGGUU-AGAUCGGCAGCCAUGUUUACAUGG--------UGCUCU--UAAAAA----UUAACCUGCU 6313 |
| 16 | Leptomastix | -CCUCUCCCGAC-UUUA-UGGGUG--UAAAGUCAACUAUUUUCGAAUAG--------AGGAG-UUGUAA-----CCCCACCCGCU 831 |
| 17 | M.occidentalis | -CCUCGCUUGUCUGCGA-UGGGCA-UCAAAGACAACAUCCUCGGGUGU---------AGCG--UUGUAACA----ACUGCCUGCU 502 |
| 18 | C. omonaga | CCUCCCCCGCGCCACGU-UGGGUG-ACGUCGGCAGCUAUGUGAGUAACAUCAUGUAGUGGGGA--AAAUU---UUUCACCUGCU 2214 |
| 19 | P. keikoae | --CGGUUCUGAGAUAA-UUGGGUUAUUACUCUCAACACGGCUUCGGCUGUG-------GGAACU-UAAACAA--UCUGACCUGCU 2970 |
| 20 | Z. potanini | --CGGAACUGAGAUACU-UGGGUUAGUACUCUCAACACGGCUUAGGCUGUG-------GGUUCC-UAAAUUAA--UUGACCUGCC 804 |
| 21 | Photinus | --UAAGAUCUGCACAAU-UGGGUU-GUUGUUGCAACACUUGUCUUACAAGUG------GGAUCU-GACAAUU--AUUGACCUGCC 4800 |
| 22 | A. potteri | --UAGAAAACAGUGCAUGUGGGUC--CUUGCACAGCUUGCAUUUUAUAUGUAAGU---GUUUUC-GAAAUACC-CACGACCUGCU 2032 |
| 23 | Hubei25 | --UAGGGAUCAUCACUC-UGGGUG-GAGAUGAUAACAUGGUGCAUGCAUCAUG-----GAUCCU-GAAAAGAA-CAACACCUGCU 6203 |
| 24 | C. osculatum | -UGCCUCUCAGUCAACU-UGGGUU-AGUUUGACAGCUAUGCUUGCAUAGU--------GAGGG---AAAGAAA-UUUAACCUGCU 157 |
| 25 | M. chinense | --UGCCAUAAUGCUCAU-UGGGUG--AUGAGCGUGCCUAGGAGUAAUUCUUAGG----AUGGUG-UAA------CCCCACCUGCU 921 |
| 26 | Araneidae | --UAGGUUCAGAUAUGUUCGGGUU-GACAUAUCAGCUUGAAAGAGC------------GAACCG--AAAUA--CCCAGACCUGCU 3322 |
| 26 | Argiope | --UAGGAACAGCUAUA-UUGGGUAG-UUGUAGCAGUUGUAUUCAAAUGAAUGCAGC--GUUCC--GAAAUA---CCAUACCUGCU 3254 |
| 28 | A. bruennichi | --UAGAAACCGCUAUA-UCGGGUAG-CUAUAGCAGCUAGUUAGUAACAUAAUUAGC--GUUUC--GAAACA---CCAUACCUGCU 2334 |
| 29 | ABPV-144I | --UAGGAACAGCUAUA-UUGGGUAG-UUGUAGCAGUUGUAUUUAAAUAAAUGCAGC--GUUCC--GAAAUA---CCAUACCUGCC 6535 |
| 30 | Dicistro_bsk | --UAGAAACCGCUAUA-UCGGGUAG-CUAUAGCAGUCAGUUAGUAACAUAGCUGGC--GUUUC--GAAAUA---CUAUACCUGCU 6375 |
| 31 | F. pressilabris | --UAGAAACCGCUAUA-UCGGGUAG-CUAUAGCAGCUAGUUAGUAACAUAAUUAGC--GUUUC--GAAAUA---CCAUACCUGCU 236 |
| 32 | Dicistro_jyt032 | -CUAGAAACCGCUAUA-UCGGGUAG-CUAUAGCAGCCAGUUAGUAACAUAAUUGGC--GUUUC--GAAAUA---CUAUACCUGCU 7054 |
| 33 | KBV | --UAGAAACCGCUAUA-UCGGGUAG-CUAUAGCAGUCGGAUAGUAAUAUAUCCGGC--GUUUC--GAAAUA---CCAUACCUGCU 6632 |
| 34 | Pogonomyrmex | --UAGAAACAGCUAUA-UCGGGUAG-CUAUAGCAGUCAGAUAAUAACAUAUUUGGC--GUUUC--GAAAUA---CCAUACCUGCU 2897 |
| 35 | Fex1 | --UAGAAACCGCUAUA-UCGGGUAG-CUAUAGCAGUCAGACAGUAAUAUGUUUGGC--GUUUC--GAAAUA---CCAUACCUGCU 6636 |
| 36 | SN11649 | --UAGAAACCGCUAUA-UCGGGUAG-CUAUAGCAGUCAGAUAGUAACAUAUUUGGC--GUUUC--GAAAUA---CCAUACCUGCU 6643 |
| 37 | IAPV | --UAGGAACAGCUGUA-CUGGGCAG-UUACAGCAGUCGUAUGGUAACACAUGCGGC--GUUCC--GAAAUA---CCAUGCCUGGC 6697 |
| 38 | SIV5 | --UAGGAACAGCUAUA-UCGGGUAG-CUAUAGCAGUCGUACAGUAACAUGUGCGGC--GUUCC--GAAAUA---CCAUACCUGGC 6471 |
| 39 | O. navus | --UAGGAACAGCUAUA-GCGGGUAG-CUAUACCAGUCGUAAGGUAACCCUUGCGGC--GUUCC--UAAACA---CCAUACCUGGC 369 |
| 40 | C. monticola | --UAGGAACAGCUAUA-UCGGGUAG-UUAUAGCAGUCGUAAGGUAACACUUGCGGC--GUUCU--GAAACA---CUAUACCUGGC 1006 |
| 41 | D. umbrophila | --UAGGAACAGCUAUA-UCGGGUAG-CUAUAGCAGUCGUAAGGUAUCACUUGCGGC--GUUCU--GAAACA---CCAUACCUGGC 5038 |
| 42 | Apara114-k141 | --UAGGAACAGCUAUA-UCGGGUAG-CUAUAGCAGUCAUAAGGUAACACUUGUGGC--GUUCU--GAAACA---CCAUACCUGGC 6734 |
| 43 | C. holgerseni | --UAGGAACAGCUAUA-UCGGGUAG-CUAUAGCAGUCGUAAGGUAACACUUGCGGC--GUUCU--GAAAUA---CCAUACCUGGC 2925 |
| 44 | Novo_mesto | --UAGGAAUCGCUAUA-UCGGGUUG-CUAUAGCAGCAUGGCUUUGGCUGUGU------UUUCC--GAAAUA---CCAAACCUGCU 6726 |
| 45 | M. rubra | --UAGGAGCAGCUAUA-UCGGGUUG-CUAUAGCAGCAUGGCUUAGGCUAUGC------GUUCC--GAAAUA---CCAAACCUGCU 6712 |
| 46 | M. japonica | --UAGGAGCAGCUAUA-UCGGGUUG-CUAUAGCAGUGAGACCUAGGUUUCAC------GUUCC--GAAAUA---CCAAACCUGCU 3202 |
| 47 | A. stella | --UAGGAACAGCUAUA-UCGGGUUG-CUAUAGCAGUGAGACUUAGGUUUCAC------GUUCC--GAAAUA---CCAAACCUGCU 1336 |
| 48 | H. azumiensis | --UAGGAACAGCUAUA-UCGGGUUG-CUAUAGCAGUAAGACUUAGGUUUUGC------GUUCC--GAAAUA---CUAAACCUGCU 2313 |
| 49 | M. chinense2 | --UAGGAACAGCUAUA-UCGGGUAG-CUAUAGCACUUAGGUAAUACUCUUAAG-----GUUCC--GAAAUA---CCAUACCUGCU 3514 |
| 50 | W. contortipes | --UAGAAACAGUUAUA-UCGGGUAG-CUAUAACAGUUAGAUUGCUAAUUUAAC-----GUUUC--GAAAUA---CCAUACCUGCU 459 |
| 51 | E. cambridgei | --UAGAAACAGUUAUA-UCGGGUAG-CUAUAACAGUUAGAUAAUAUAUUUAAC-----GUUUC--GAAAUA---UCAUACCUGCU 6681 |
| 52 | L. humile | --UAGGAGCAGCUAUA-UCGGGUUG-CUAUAGCAGCGUUAAAUUUAUUUGACGC----GUUCC--GAAACA---CCAAACCUGCU 6656 |
| 53 | H.yanbaruensis | --UAGGUUCAGCCAUA-UCGGGUUG-CUAUGGCAGCAUGUAGGUAACACUGUAUGC--GAACU--GAAAUA--CCCAGACCUGCU 1093 |
| 54 | SIV1 | --UAGGAACAGCUAUA-UCGGGUUG-CUAUAGCAGUCAGGAUGUCAUUCUGGC-----GUUCC--GAAAUA--CCCAAACCUGCU 4425 |
| 55 | M. illinoiensis | --UAGGAACAGCUAUA-UCGGGUAG-CUAUAGCAGUCAGCAUAAAAUGUUGGC-----GUUCC--GAAAUA---CCAUACCUGCU 75 |
| 56 | SIV12 | --UAGGAGCAGCUACAUUCAGGUU-GAUGUAGCAGUUAUUUAGUAACAUAAGUAGC--GUUCC--GAAACA--CCCAGACCUGCC 6099 |
| 57 | P. tepidariorum | --UAGGAGCAGUUACAUUCAGGUU-GAUGUAACGGUAGUAUAGUAACAUAUGCUAC--GUUCU--GAAACA--CCCAGACCUGCC 3231 |
| 58 | Hubei1 | --UAGGAGCAGCUACAUUCAGGUU-GAUGUAGCAGUUGUUAGGUAACACUAGCAGC--GUUCC--GAAACA--UCCAGACCUGCC 6144 |
| 59 | P. cryptcolens | --UAGAAACAGCUACAUUCAGGUU-GAUGUAGCAGUUGUUAGGUAACACUAGUAAC--GUUUC--GAAACA--CCCAGACCUGCC 1643 |
| 60 | Holoplatys | --UAGGAAUCGCUAUAUUCGGGUG-GAUGUAGCAGCAUUGCUUUGGCGAUGUU------UUCC--GAAAUA--CCCACACCUGCU 4062 |
| 61 | A. yesoensis | --UAGGAAUCGUAGCAUUCGGGUA-GAUGUUACAGCGUGGCUUAAGCUGCGUU------UUCC--GAAACA--CCCAUACCUGCC 6462 |
| 62 | Haeterina | --UAGGAACAGCAUAA-UUGGGUUG-UUUAUGCAGCCAUGAUGUAACAAUCGUGGC--GUUCC--GAAACA---C-AGACCUGCU 234 |
| 63 | P. labialis | --UAGGUUCAGCUAUA-UCGGGUUG-CUAUAGCAGAUAGCCUUAGGGUUAUC------GAACC--GAAACA---CCAAACCUGCU 1750 |
| 64 | Wuhan11 | --UAGGUUCAGCCAUA-UCGGGUUG-CUAUGGCAGUAUGGAAUUUAUUCUGUAC----GAACC--GAAAUA---CCAAACCUGCU 6310 |
| 65 | E.nubilus | --UAGGUUCAGUAUCA-UCGGGUUG-CUGAUACAGCAUGAAAUUUAUUUUAUGC----GAACC--GAAACA---CCAAACCUGCU 3725 |

Table S5. Sequence alignment of type 6b IRESs, numbered to indicate 5' and 3' terminal nucleotides and annotated to show structural elements (as shown in Figs. 6A and 6B) and the initiation codon (red font). Bold font indicates nucleotides that are conserved in over 90% of members of this group (excluding domain 1 sequences in IRESs #13-17 that appear to be type 6a/type 6b recombinants. Loop L1.1a 'GAUC' and 1.1b 'UGC' motifs in these sequences are characteristic of type 6a IRESs and are indicated by blue font). IRESs Nucleotides that are predicted to engage in base-pairing in helical elements P1.1, P1.2, P1.3, P2.1, P2.2, P2.3, P2.4, P3.1, P3.2 and P3.3 are shaded and bracketed. Note that sequences # 4, 11, 13, 16, 18, 23, 25, 26, 33, 38, 40, 45, 54, 56, 59 and 64 are numbered according to the deposited (antisense) sequences.

Aligned sequences are from: (1) Beihai mantis shrimp virus 4 strain BHXG23944 (NC_032464.1), (2) Beihai picorna-like virus 93 strain BHBJDX18546 (NC_032588.1), (3) Wenzhou shrimp virus 7 strain beimix73547 (NC_032420.1), (4) TSA: *Eriocheir sinensis*, contig Unigene25727_A0A (HAAX01025575.1), (5) Mud crab dicistrovirus (NC_014793.1), (6) Taura syndrome virus, complete genome: NC_003005.1, (7) TSA: *Oratosquilla oratoria* Unigene52730_HQ (GGQQ01198286.1), (8) Beihai mantis shrimp virus 5 strain BHXG25299: NC_032434.1, (9) *Macrobrachium rosenbergii* Taihu virus strain cn-taihu100401 (NC_018570.2), (10) TSA: Zorotypus caudelli s3164_L_3735_2_a_74_4_l_5165 (GAYA02046772.1), (11) TSA: *Zoanthus* sp. QL-2018 (GGTW01107769.1), (12) TSA: *Carcinus maenas* Cluster-50807.0_TR49096_c0_g1_i1 (GFXF01042110.1), (13) TSA: *Dermanyssus gallinae* isotig17344 (GAIF01027790.1), (14) TPA_asm: *Pycnopodia helianthoides* associated picornavirus 2: BK059603.1, (15) Beihai picorna-like virus 91 strain BHJJX25930 (NC_032638.1), (16) TSA: *Leptomastix dactylopii* s5308_L_19038_0_a_20_5_l_1112 (GBNE01033170.1) , (17) TSA: *Metaseiulus occidentalis* MITE_454Assem.sd.12189.C1 (JL020592.1), (18) TSA: *Cyclosa omonaga* IDV#6323 (IARB01014260.1), (19) TSA: *Paikiniana keikoae* IDV#5510 (ICCT01024043.1), (20) TSA: *Zelotes potanini* IDV#7284 (ICMX01034693.1), (21) TSA: *Photinus pyralis* TR38402_c0_g2_i1 (GEOW01072248.1), (22) TSA: *Agelenopsis potteri* IDV#3933 (IBWF01002267.1), (23) Hubei picorna-like virus 25 strain QTM27237 (NC_033190.1), (24) TSA: *Contracaecum osculatum* Sequence_31261 (GKNQ01031029.1), (25) TSA: *Monomorium chinense* mRNA (LI886324.1), (26) TSA: *Araneidae* gen. sp. IDV 2115 (ICNA01020445.1), (27) TSA: *Argiope* sp. IDV 2158 (IBEL01023942.1), (28) TSA: *Argiope bruennichi* IDV#6051 (IBFJ01038969.1), (29) Acute bee paralysis virus isolate ABPV/144I (MN565031.1), (30) MAG: Dicistroviridae sp. isolate bsk136dic03 (MN905957.1), (31) TSA: *Formica pressilabris* mRNA, contig: comp44934_c0_seq1 9 (LI304347.1), (32) MAG: Dicistroviridae sp. isolate jyt032dic1 (MN918742.1), (33) Kashmir bee virus (NC_004807.1), (34) TSA: *Pogonomyrmex californicus* RNA (IAAD01013993.1), (35) *Formica exsecta* virus 1 isolate Fex1 (NC_023021.1)(, (36) Dicistroviridae sp. isolate YSN11649 (MW826503.1), (37) Israel acute paralysis virus of bees strain DVE31-OP3-PA-USA-2007 (EU436423.1), (38) Solenopsis invicta virus 5 isolate Formosa (MF593921.1), (39) TSA: *Oecobius navus* IDV#7047 mRNA (IBWY01003970.1), (40) TSA: *Cyclosa monticola* IDV#14 mRNA (IBFS01011564.1), (41) TSA: *Dolichognatha umbrophila* IDV#3143 mRNA (IBJS01018089.1), (42) MAG: Aparavirus sp. isolate 114-k141_49754 (MZ679095.1), (43) TSA: *Cataglyphis holgerseni* TRINITY_DN10185_c0_g2_i1-1935-0.348511 (GJGD01023859.1), (44) MAG: Novo Mesto dicistrovirus 1 isolate JUR20SW1 (OL472188.1), (45) Myrmica rubra picorna-like virus 4 (MW314635.1), (46) TSA: *Meta japonica* IDV#4242 (IBCJ01032778.1), (47) TSA: *Araneus stella* IDV#4225 (IBQX01026271.1), (48) TSA: *Himalaphantes azumiensis* IDV#4087 (IBSV01021412.1), (49) *Monomorium chinense* mRNA, contig: comp57089_c0_seq1 (LI881333.1), (50) TSA: *Weintrauboa contortipes* IDV#2209 (IBUA01013242.1), (51) TSA: *Ero cambridgei* IDV#6464 mRNA (IBDP01035891.1), (52) Linepithema humile virus 1 isolate variant LhuIT (MW314674.1), (53) TSA: Heptathela yanbaruensis IDV#55 (IAWJ01012018.1), (54) Solenopsis invicta virus 1 (NC_006559.1), (55) TSA: *Macromia illinoiensis* breed wildtype C58429_a_4_0_l_563 (GCBZ01011316.1), (56) *Solenopsis invicta* virus 12 isolate Formosa (MH727529.1), (57) TSA: *Parasteatoda tepidariorum* IDV#5621 (ICMQ01031218.1), (58) Hubei orthoptera virus 1 strain ZCM21319 (NC_032766.1), (59) TSA: *Pholcus crypticolens* IDV#962 (IBBI01018343.1), (60) TSA: *Holoplatys* sp. IDV 6692 IDV#6692 (ICCD01033051.1), (61) TSA: *Antrodiaetus yesoensis* IDV#4252 (IBEE01037285.1), (62) TSA: Hetaerina sp. AD-2014 breed wildtype C128027_a_6_0_l_296 (GCKS01012094.1), (63) TSA: *Phrurolithus labialis* IDV#7097 (IAUN01026585.1), (64) Wuhan insect virus 11 strain CJLX30623 (NC_033460.1), and (65) TSA: *Episinus nubilus* IDV#6266 (IAEL01040798.1)

Table S6. Sequence alignment of type 6i-1 IRESs.

P1.1 · P1.2 · P1.3 · SLVI · P2.1

| # | Viral sequence | | |
|---|---|---|---|
| 1 | S.nigra | 3030 | -CUACGUGAUCGGGGCUGGUUCUAAAUUGCCG-AACAAGAAUCAGC-----GACUGCUGU-AGU----------UCUCGAAGCCUAGCAGCCUAGAGA------------AGUGGG |
| 2 | Y.lyrica | 3162 | -CCGUUGUGAUCUGGACUAGUUCUAAAUUGUCGAAAAAGAAUUAGUCU-------UGCUGCGGGU-----------CCCUGAGUUUAGCGACCGAGGG------------AGAUGGG |
| 3 | D.argyoides | 24 | -CCAUGUGAUCGGGACUCAGCUCAGAUCGUCG-AAACGAGAGGAGU------GACUGCUAUGGUGGC--------GACUGAGGCAAUCCCUCAGAAGUC-----------GAUGGG |
| 4 | Y.striatipes | 306 | CCAUGUGAUCGGGGUUGGAGUCAGAUCGUCG-AAACGAUUCCAGC------GACUGCUGUGGU----------GGGUGUAUGAGCAAUCCUCAUGCACUC---------GAUGGG |
| 5 | Lycosidae | 11 | ACCAUGUGAUCGGGACUUUACUUAGAUCGUCG-AAAUGAGUAUAGU------GACUGCUAUGGU----------GGGCGUAUGAGCAAUCCUCAUGCGCUC---------GAUGGG |
| 6 | W. okinawensis | 450 | ACCAUGUGAUCGGGACUAUUCGUAGAUGAUGG-AAUAUCGAGUAGU------GACUGCU-GGGU------------GCGCUUGUUGGCAACAAUGAGUGC----------GGUGAG |
| 7 | s59k141_259911 | 6848 | -ACCGUGUGAUCGGGAUCAGCGGCAGAUUGCCG-AAAUACCGCUGAU------GACUGCUGCGGU----CGCUCUGGCGAGUGGUGUAAGCCAUGCACAAGAGAGCG----GAAGGG |
| 8 | Dicistro.347R | 6625 | AACCAUGUGAUCGGGCUCGAUUUCAGAUUGCCG-AAACGGAAUCGAGU-----UAC-GCUG-GGUU---------------CACCUAGCGGUGA-----------------GUUGGG |
| 9 | Metaseiulus | 336 | --CCAAGUGAUCGGAGAUAAGUGUAGAUGACGG-AAUAAUGCUUAACU------ACCGCUG-GGU-----------GGGUUCUAGCUAGCGCAAAGAACUC----------CAUGGG |
| 10 | E.pustulosa | 92 | CCACGUGAUCGGGGCUAGUUGUAGAUGAUGG-AAUGACGACUAGC------GACUGCUG-G-----------GUGCGUACCCCAUCACCAAGGGAAUGCGC--------GUUGGG |
| 11 | Oxyopes | 383 | -CCAAGUGAUCGGAGUGAGGAGAAGAUGAUGG-AAAUUGAUUCGCU-------ACCGCUG-GGUGU---------GCAUGUAUGUAAAUAUGUGUUC-------------GAUGAG |
| 12 | Bundaberg 2 | 4817 | -CCAUGUGAUCGGGACUAUUCGUAGAUGAUGG-AAUAUCGAGUAGU------GACUGCUG-GGU-----------GCACUUGUUGGCAACAAUGAGUGC-----------GGUGAG |
| 13 | Bemisia | 4324 | CCACGUGAUCGGGAUCUUUCGUAGAUGAUGG-AAAGACGGAAGAU------GACUGCUG-GGU-----------GCGCUUGCUAGUGAUAGUAAGUGC-----------GGUGAG |
| 14 | T.vaporariorum | 6944 | -CCAUGUGAUCGGGAUCUUCUGUAGAUGAUGG-AAAGACGGAAGAU------GACUGCUG-GGU-----------GCGCUUGCUAGUAAUAGUGAGCGC-----------GGUGAG |
| 15 | Leptomastix | 3093 | ACCACGUGAUCGGGAUCUUCUGUAGAUGAUGG-AAAGACGGAAGAU------GACUGCUG-GGU-----------GCGCUUGCUAGCAAUAGUAAGUGC-----------GGUGAG |
| 16 | Weiv_war204 | 6797 | ACCAUGUGAUCGGGACUAAGAAUAGAUGACGG-AAACGUUCUUAGU------GGCUGCUAUGGU----------GUGCGCGAGCUUAAGUGAAACGCAC-----------GUUGAG |
| 17 | N.subpullata | 3556 | ACCACGUGAUCGGGGCUAGGAUCAGAUGACGG-AAACGAGACUAGC------GACUGCUGUGGU---------ACGCGCCAGCUUAAGUGAAACGCGU------------GUUGGG |
| 18 | Silene | 6116 | -CCAUGUGAUCGGAGCUAGAGUCAGAUGACGG-AAACGUCUCUAGU------UACUGCUAUGGU---------UGGCGCCAGCUUAAGUGAAACGCCA------------AGUGGG |
| 19 | M.testaceus | 160 | ACCAUGUGAUCGGAGCUAGAUUCAGAUGACGG-AAACGAAUCUAGU------UACUGCUAUGGU-----------------AUGCUUCGGCAU-----------------GAUGGG |
| 20 | Marpissa | 2841 | -CCACGUGAUCGGGGUUCGAAUUAGAUGACGG-AAACGAUUCGAGC------GACUGCUGUGGU-----------------GUGUGAAAACAC-----------------GUUGGG |
| | | | L1.1a L1.2a L1.2b L1.1b |

P1.3 · P2.1 · SLIV · P2.3 · P3.3

| # | Viral sequence | | |
|---|---|---|---|
| 1 | S.nigra | AGAGCUAUUUAGCUCU-AUUGCGGACUGUUUGCCCACUCAAA-CAAUAGGAUAAUCCGCCGGCACGUCUGGUGAAGCA-UUAAGUUGUGCCCAUCUUAAGACGUUAAACCACUAACUUGCC | 2826 |
| 2 | Y.lyrica | AGAGCUAUUUAGCUCU-AUUGCAGACUGUUUGCCCAUCUAAAA-CAAUAGGAUAAUCUGCCGACACGUCUGGUGAAGCA-UUAAGUUGUGCCCACUUUAAGACGUUAAACCACUAACUUGCC | 2959 |
| 3 | D.argyoides | AGAGGUAUUUACCUCU-AUUGCAGACUGUUUGCCCAUCU-AAA-CAAUAGGAUAAUCUGCCGACGGGUCUGGUGCAGCA-UUAAGUGGUGCGCAUCUUAAGACUUUAAACCAUUCACUUGCC | 230 |
| 4 | Y.striatipes | AGAGGUAUUCACCUCU-AUUGCAGACUGUUUGCCCAUCUAAAA-CAAUAGGAUAAUCUGCCGACGCGUCCCCUGGAGCA-UUAAGUGGUGCCCUUG-CAAGACGUUAAAUCACUCACUUGCC | 512 |
| 5 | Lycosidae | AGGUGUAUUUACAUCU-AUUGCAGACUGUUUGCCCAUCUAAAAUAUAUAGGAUAAUCUGCCGACGCGUCGACCAAUGCA-UUAAGUGGUGCUAGGUGUCAGACGUUAAAACAUUCACUUGCC | 220 |
| 6 | W. okinawensis | AGAAUUAUUUAGUUCC-AUUGCAGACUGUUUGCUCACCUAAAAAUAAUAGGAUAAUCUGCCCAUCCGUCGAUAGCUGACGUUGAGUGACGAGCUAUU-AAGACGUUAACAUAUUCACUUGCC | 655 |
| 7 | s59k141 | AGAGCUUUUCAGCUCU-AGAGGAUGAUGUUUGCCCUUCUAAAA-CAAUAGGAUAUCGUCCCGGCACGUCACGAUCUGGC-AUAAGUGGUCCAGCUCAUUCGACGUUAAACAGUUCACUUGCU | 7067 |
| 8 | Dicistro.347R | AGAGGUAUUUACCUCC-AUUGCAGACUGUUUGCCCAACUGAAUAUAAUAGGAUAAUCUGCCGGCACGACCCGAAGAAGG-AUAAGUGGCUUCUAGG--UAGUCGUUAAACAAUUCACUUGCC | 6819 |
| 9 | Metaseiulus | AGGGCUAUUUAGUCCC-AUUGCAGACUGUU-GCCCAUGU-AAA-UAAUAGGAUAAUCUGCCCGUCCGUCGAUUCGAACA-UUGAGUGAUGUAGAAUU-CAGACGU-AAACAAUUCACUUGCC | 7 |
| 10 | E.pustulosa | AGAGCUAUUUAGCUCU-AUUGCAGACUGUUUGCCCAACUGAAA-CAAUAGGAUAAUCUGCCCAUCCGUCGGUAGCUAGCAUUGAGUGAUGCGCUGCU-UAGACGUUAACAUAUUCACUUGCC | 298 |
| 11 | Oxyopes | AGGGUUAUUUAGCUCU-AUUGCAGACUGUUUGCUCAUCUAAAAGAAUAGGAUAAUCUGCCCAUCCGUCGAUAGCUGACGUUGAGUGACGAGCUAU-UUAGACGUUAACAUAUUCACUUGCC | 180 |
| 12 | Bundaberg 2 | AGAAUUAUUUAGUUCC-AUUGCAGACUGUUUGCUCACCUUAAAAUAAUAGGAUAAUCUGCCCAUCCGUCGAUAGCUGACGUUGAGUGACGAGCUAU-UAAGACGUUAACAUAUUCACUUGCC | 5021 |
| 13 | Bemisia | AGAAUUAUUUAGUUCC-AUUGCAGACUGUUUGCUCACCUUAAAACAAUAGGAUAAUCUGCCCAUCCGUCGAUAGCUGACGUUGAGUGACGAGCUAU-UUAGACGUUAACAUAUUCACUUGCC | 4528 |
| 14 | T.vaporariorum | AGAGCUAUUUAGUUCC-AUUGCAGACUGUUUGCUCACCUAAAACAAUAGGAUAAUCUGCCCAUCCGUCGAUAGCUGACGUUGAGUGGCGAGCUAU-UUAGACGUUAACAUAUUCACUUGCC | 7148 |
| 15 | Leptomastix | AGAAUUAUUUAGUUCC-AUUGCAGACUGUUUGCUCACCUUAAAACAAUAGGAUAAUCUGCCCAUCCGUCGAUAGCUGACGUUGAGUGACGAGCUAU-UUAGACGUUAACAUAUUCACUUGCC | 2888 |
| 16 | Weiv_war204 | AGGGUUAUUCAGCCCU-AUUGCAGACUGUUUGCUCAACUAAAAAUAAUAGGAUAAUCUGCCCGUCCAUCCCUUUAGA-CAUUAAGUGGUGCAGAAG-AAAGAUGUAAAAUUAUUCACUUGCC | 7003 |
| 17 | N.subpullata | AGGGUUAUUUAGCCCC-AUUGCAGACUGUUUGCCCAACUAAAACAAUAGGAUAAUCUGUCCGUCCGUCCCUCCAAAGCAUUGAGUGGUGCUGGAG-UAAGACGUUAAAUCAUUCACUUGCC | 3763 |
| 18 | Silene | AGGGUUAUUUAGCCCU-AUUGUAGACUGUUCGCCCACUUUAAAAUUAAAGGAUAAUCUGCCCGUCCGUCGGUAGCUUGCAUUGAGUGAUGCGCUGCU-UAGACGUUAA-CAUUUCACUUGCC | 6321 |
| 19 | M.testaceus | AGAGUUAUUUAGCUCU-AUUGCAGACUGUUUGCCCAUCUAAAA-CAAUAGGAUAAUCUGCCCGUCCGUCGGCUGUAUGCAUUGAGUGAUGCGCAGUU-UAGACGUUAA-CAGUUCACUUGCC | 352 |
| 20 | Marpissa | AGGGUUAUUUAACCCU-AUUGCAGACUGUUUGCCCAACUAAAAAUAAUAGGAUAAUCUGCCCGUCCGUCGGCUGCUAGCAUUGAGUGAUGCGCAGUU-UAGACGUUAA-CAAUUCACUUGCC | 3033 |
| | | L2.1 L2.2 L3.1 L3.2 | |

Supplementary table 6. Sequence alignment of type 6i-1 IRESs, numbered to indicate 5' and 3' terminal nucleotides and annotated to show structural elements (as shown in Fig. 8A), the initiation codon (red font) and nucleotides conserved in over 90% of members of this group (bold). Aligned sequences are from: (1) TSA: *Sisyra nigra* breed wildtype s7271_L_25799_0_a_16_6_l_3969 (GCTI01024755.1), (2) TSA: *Yunohamella lyrica* IDV#3858 mRNA, IDV3858-W1_S140036245 (IAVH01036245.1), (3) TSA: *Devadatta argyoides* breed wildtype C118580_a_5_0_l_729 (GCKO01007350.1), (4) TSA: *Yaginumaella striatipes* IDV#1934 mRNA, IDV1934-W1_S200041956 (ICAP01041956.1), (5) TSA: *Lycosidae* gen. sp. IDV 5014 IDV#5014 (IBZZ01034293), (6) TSA: *Wadicosa okinawensis* IDV#2840 mRNA, IDV2840-W1_S120004398 (IBSA01004398.1), (7) MAG: Dicistroviridae sp. isolate s59-k141_259911 (MZ679078.1), (8) MAG: Dicistroviridae sp. isolate 347R-k141_1033911 (MZ679053.1), (9) TSA: *Metaseiulus occidentalis* MITE_454Assem.221.CB1 (JI991228.1), (10) TSA: *Eriophora pustulosa* IDV#6816 mRNA, IDV6816-W1_S160004095 (IBIK01004095.1), (11) TSA: *Oxyopes* sp. IDV 5198 IDV#5198 mRNA, IDV5198-W1_S70025346 (ICGS01025346.1), (12) Bundaberg bee virus 2 isolate QLD-4 (MG995700.1), (13) Bemisia tabaci dicistro-like virus 1 isolate CAU-Q1 (MW256674.1), (14) TSA: *Trialeurodes vaporariorum* Cluster-3686.21761 (GHMB01082144.1), (15) TSA: *Leptomastix dactylopii* C113859_a_35_0_l_3587 (GBNE01004402.1), (16) MAG: Weivirus-like virus sp. isolate war204pic4nc (MT138417.1), (17) TSA: *Neoscona subpullata* IDV#867 mRNA, IDV867-W1_S140009447 (IBPZ01009447.1), (18) TPA_asm: Silene latifolia dicistro-like virus 1 isolate Sl1 (BK059255.1), (19) TSA: *Mimetus testaceus* IDV#5828 mRNA, IDV5828-W1_S100020932 (IAZF01020932.1), and (20) TSA: *Marpissa mashibarai* IDV#7083 mRNA, IDV7083-W1_S30006465 (IBBC01006465.1).

Table S7. Sequence alignment of type 6i-2 IRESs.

| # | Viral Sequence | | P1.1 P1.2 P1.3 SLVI P2.1 |
|---|---|---|---|
| 1 | C.japonica | 5949 | ACGGGUAGAUCGGGGCACUCUUAAAAUGGUCA-----UAUUUGAGUGCAGCAAAUCCUGCU-CCCGU-----------GCGUGUUACCUUGUCACUUUCACGC---------GUUAGG |
| 2 | I.insidiosa | 1459 | ACUCUAUGAUCGGGGCACUCUAAAAAUGGUCG-----CAUUUGAGUGCAGCAAAUCCUGCU-GGAGU----------GGACGUUAAACUUGUAACUUGCGUUC---------GGUGAG |
| 3 | E.nubilis2 | 483 | ACUCUGUGAUCGGGGCACUCUGAAAAUGGUCG-----CAUUUGAGUGCAGCAAAUCCUGCU-GGAGU----------GAACGUUAAACUUGUAACUUGCGUUC---------GGUGAG |
| 4 | G.exsiccatum | 7117 | CCACGUGAUCGGGGCCAGUCGCAAAUGAUCG---AAAGCGGCUGUGCAGCAAAUCCUGCU-GUGGU-------GGAUGCUGGACGUCAAGGCUAUCCAGUACUCC------GAAGGG |
| 5 | D.peniculus | 5144 | ACCACGUGAUCGGGGCCAGUCGCAAAUGAUCG---AAAGCGGCUGUGCAGCAAAUCCUGCU-GUGGU-------GGAUGCUGGACGUCAAGGCUAUCCAGUACUCC------GAAGGG |
| 6 | W.golovatchi | 3036 | CCACGUGAUCGGGGCCAGUCGCAAAUGAUCG---AAAGCGGCUGUGCAGCAAAUCCUGCU-GUGGU-------GGAUGCUGGACGUCAAGGCUAUCCAGUACUCC------GAAGGG |
| 7 | P.yaginumai | 1740 | ACUUCUUGAUGUGGGCUUGUAAUAUAUGACCA----UUAAUUACAUGCAGCAAAUCCUCCGAGAAGU-------GGGUUAAUAGUCGGCAGCGUCUUCCUUAACCC------GUUGGG |
| 8 | M.sadamotoi | 1963 | ACCUCGUGAUCGGGGCUGGUCGAUAAUUGUCG----CCGUCGACCAGCAGCAAAUCCUGCU-GAGGU----------GCGCGAAGCUUAGCGGCGGAUGCGC----------GGUGGG |
| 9 | Sanya_dicistro2 | 6722 | ACCUCUAGAUCGGGGCAUCGGUCAAAUUGCCA----UCAGACCGGUGCAGCAAAUCCUGCU-GGGGU---------- GCGUGAAUCUUUGCGGAGUAGCGC----------GAUGGG |
| 10 | H.panicea | 97 | CCAGUGAUAGGGGCUCUGAGUUUAUGGCCG----AUAACACAGAGCAGCAAAGCCCGUUUCCUGGU----AGUUUUGACCUGACCGAAAGGCAGUUACUCAAAACU----GGU-GC |
| 11 | L.argiopiformis | 2524 | ACUUCGUGAUCGGGGUCUCGGACUAAUCGUCA----UCGGUCUGAGACAGCAAAUCCUGCU-GAAGU----------GCGUGAAUCCUAGCAGAGGCUACGC----------GAUGGG |
| 12 | Portia | 3248 | ACUUCGUGAUCGGGGUUGCGGACUAAUCGUCA----UCGGUCUGCAACAGCAAAUCCUGCU-GAAGU----------GCGUGAAUCCUAGCAGAGGCUACGC----------GAUGGG |
| 13 | M.mirabilis | 3126 | ACCUCGUGACCGGGGCUGUGGGUUAAUCGUCA----UCGACCCGCAGCAGCAAAUCCUGCU-GAGGU----------GCGUGAAUCCUAGCAGAGGCUACGC----------GAUGGG |
| 14 | Portia#2 | 1620 | ACCUCGUGAUCGGGGCUGCGAACUAAUCGUCA----UCGGUUCGCAGCAGCAAAACCUGCU-GAGGU----------GCGUGAAUCCUUGCGGAGGCUGCGC----------GAUGGG |
| 15 | Uluborus | 3263 | ACCUCGUGAUCGGGGCUGCGAACUAAUCGUCA----UCGGUUCGCAGCAGCAAAACCUGCU-GAGGU----------GCGUGAAUCCUUGCGGAGGCUGCGC----------GAUGGG |
| 16 | Myrmarachne | 3273 | ACCUCGUGAUCGGGGCUGCGAACUAAUCGUCA----UCGGUUCGCAGCAGCAAAACCUGCU-GAGGU----------GCGCGAAUCCUUGCGGAGGCUGCGC----------GAUGGG |
| 17 | H.yanbaruensis | 2787 | ACCUCGUGAUCUGGGCUACUAUCUAAUCGCC---AAAAGGAAAGUAGCAGCAAACCCUGCU-GAGGU----------------GGUUCGAAAGAACC---------------AGUGAG |
| 18 | P.praticola | 3321 | ACUUCGUGAUCGGAGCUGACGACAGAUCGCUG----AAGGUCGUCAGCAGCAAAUUCUGCU-GAAGU---------GGUGUAAUAGAUAGCAAUAUUAACAUC---------GAUGGG |
| 19 | Cybaeus | 7310 | GCUUCGUGAUCGGGGCUGUGUCUAAAUCGUCG---UUUGGGACACAGCAGCAAAUCCUGCU-GAAGU---------GGGUACUCCGAUAGCAAGGAGAUACUC---------GGUGAG |
| 20 | E.nubilis | 834 | ACCAAUUGAUCAGGGCUUGAUUCAAAUGACUG---AAGCGAAUCUUGCAGCAAAUCCUGCGAUUGGU-------AGGUUAAUAGUCCUCAUGGUCUUCCUUAACCU-------AGUGG- |
| 21 | I.narutomii | 2876 | ACCUAUUGAUCGGGGCUAGAUUCAAAUGACUG---AAGCGAAUCUAGCAGCAAAUCCUGCGAUAGGU-------AGGUUAAUAGUCCUCACGGUCUUCCUUAGCCU-------AGUGG |
| 22 | P.yesoensis | 2983 | ACCUCGUGAUCGGAGCUUUUCUUCA-UUCGCG----UAGAAGAAUUGCAGCAAAUUCUGCU-GAGGU------GUGUAGUAUUCUCCUCUGGAGCACUUACUAUGC------GUUGGG |
| 23 | Galloisiana | 5073 | ACCUUGUGAUCGGGGCUUAGUUCGAUUCGACG----UAAGAAUUAAGCAGCAAAUCCUGCU-GAGGU------CGAUUGUAUUCUCAUUGUGAGCACUUGCAAUCG------GGUGGG |
| 24 | A.kamurai | 2526 | ACCUCGUGAUCGGGGCUUAACUCGAUUCGGCG----UAAGAGUUAAGCAGUAAAUCCUGCU-GAGGU------CGAUUGUAUUCUCAUUGAGAGCACUUGCAAUCG------GGUGGG |
| 25 | R27-k141 | 6159 | ACGGCGUGAUCGGGGCUGCGAUCAAAUGACUG----AAGGGUCGCAGCAGCAAAUCCUGC-UGCCGU--GGUGUAUAUAUUUAGGUACUUAGGUAUUUUGAUGUAAACAUC-GGAGCG |
| 26 | O.crucifera | 515 | ACUCUGUGAUCGGGGCUCCUCGGCAAUCACCG---CCCGCUGAGGAGCAGCAAAUCCUGCU-GGAGUCUCCUGAUGGCCCGUCUAGUUCCUAGAGGGAGUACCAUCGAGGAGGAUGGG |
| 27 | P.mira | 3253 | CUCUGUGAUCGGGGCUCCUCGGCAAUCACCG---CCCGCUGAGGAGCAGCAAAUCCUGCU-GGAGUCUCCUGAUGGCCCGUCUAGUUUCUAGAGGGAGUACCAUCGAGGAGGAUGGG |
| 28 | R.ishigakiensis | 3330 | ACUGAGUGAUCGGGGCUUGGCUAUUUUGGCAG---AAUGUGGUCAAGCAGCAAAUCCUGCU-UCAGU------GCGUUGCCGCCCUCUUGGGGGCUACGUCAACGC------GGUGGG |
| 29 | S.rubiginosus | 1078 | ACCUCGUGAUCGGGGCUUCGGACUAAUGGCAG---CCCAGUCCGUUGCAGCAGAUCCUGCU-GAGGU---------UCUCUCUGACUCAGCUACUCAGGGAGA---------GGUGGG |
| 30 | Sinipoda | 209 | ACCUCGUGAUCGGGGCUUCGGACUAAUGGCAG---CCCAGUCCGUUGCAGCAGAUCCUGCU-GAGGUUC---------UCUCUGGCUCAGCUACUCACAGAGA---------GGUGGG |
| 31 | Picorna45cR-k141 | 578 | ACCCCGUGAUCAGGGCUUCGGACUAUUCGCCG---GACGGUCCGAUGCAGCAAAUCCCGCU-GGGGU----------GCUUAUCUGGGAAAUCUCAGUCAGC----------GGAGGG |
| 32 | Picorna_251-k141 | 1252 | CCCGUGAUCAGGGCUUCGGACUAUUCGCCG---GACGGUCCGAUGCAGCAAAUCCCGCU-GGGGU----------GCUUGUCUGGGAAGUCCCAGUCAGC----------GGAGGG |
| 33 | Picorna_155-k141 | 1426 | ACCCCGUGAUCAGGGCUUCGGACUAUUCGCCG---GACGGUCCGAUGCAGCAAAUCCCGCU-GGGGU----------GUCUGACUGGAAAAUCUCAGUCGAC----------GGAGGG |
| | | | L1.1a L1.2a L1.2b L1.3 L1.1b |

| # | Viral Sequence | P1.3 / P2.1 / SLIV / P2.3 / P3.1 / P3.2 | |
|---|---|---|---|
| 1 | C.japonica | AGAGUUUUCUAGCUCU-GUUGUCGAUUGCU-GCCUAACUGAAAAUAAUAGGGAAAUCGUCUGACCA-CAAUUCAUAUCGUC-UAAUCCAAUGAUAUGU-----------CUUAUUUGGAUUGCU | 6149 |
| 2 | I.insidiosa | AGAGUUAUUUAGCUCU-GUUGUCAGUUGCU-GCUCACCAUAUAAUAAUAGGGAAACUGUCCGACCA-AACCUCAUGCAG-AUUAAUCCGUCUGUAUGU----------UGACUCAUGGAUUGCU | 1661 |
| 3 | E.nubilis2 | AGAGUUAUUUAGCUCU-GUUGUCAGUUGCU-GCUCACCUAAUAAUAAUAGGGAAACUGUCCGACCA-AACUUCAUAUAG-AUUAAUCCGUCUAUAUGU----------CGAUAUAUGGAUUGCU | 685 |
| 4 | G.exsiccatum | AGGGGUAUUUACCCCU-AGAGGCAGACGUGUGCCCUUCUAAAAACAAAAAGCUUUCUGCCCGAUC--UAUAUCUCGCAAGCUUAAGUCAGCUGCGCGGAUG--------AAUAAAGGACUUGCU | 7329 |
| 5 | D.peniculus | AGGGGUAUUUACCCCU-AGAGGCAGACGUGUGCCCUUCUAAAAAUAAAAAGCUUUCUGCCCGAUC--UAUAUCUCGCAAGCUUAAGUCAGCUGCGCGGAUG--------AAUAAAGGACUUGCU | 5356 |
| 6 | W.golovatchi | AGGGGUAUUUACCCCU-AGAGGCAGACGUGUGCCCUUCUAAAAAUAAAAAGCUCUCUGCCCGAUC--UAUAUCUCGCAAGCUUAAGUCAGCUGCGCGGAUG--------AAUAAAGGACUUGCU | 2824 |
| 7 | P.yaginumai | AGAGGUGUUUACCUCU-AGUGAUAAAUGUUUGCCCAACUAAAAUAUAUAGGAUAUUUAUCUGUUC-ACUCUGUAAAUAUGUUUAAGUGUUCAUUUUACUAGGAGUCUAAACAAAAUCACUUGCA | 1962 |
| 8 | M.sadamotoi | AGAGCUAUUUAGCUCU-AUUGAUUAGUGUGUGCCCACCCAAAA-CAAUAGGCUACUAAUUCGGCAUGC-UACCGAUGGACGUUAGGUUACGCCAUCGGU------CUAAGAAA-UUAACCUCAA | 2169 |
| 9 | Sanya_dicist2 | AGAGUUAUUUAGCUCU-AUUGAUAAGUGUUUGCCCAUCUCAAAAAUAUAGGAUACUUAUCUGGCAUGC-UUCUGGCUCAGAUUAGGUGAUCGAGUCGGAA------AAAGAAAAUUCACCUCAA | 6929 |
| 10 | H.panicea | AGAGUUCUCUAGCUCU-AUUGAUCAGUGUGUGG-CACCUAAAA-UAA-AGGCUACUGAUCCGGCC--CAUCUAACAACCUGUCUGAGUCAACCUAUGUGAUG-AUGUAAAAUAACCGACUUCCU | 316 |
| 11 | L.argiopiformis | AGGACUAUUUAGUCCU-AGUGAUAAAUGUUUGCCCAUCUAAAAAUAAUAGGAUAUUUAUCUGGCGUGC-GUUGCAACAGCGUUAGGUGACGUGUUGUGAC------AAAGAAAAUUCACCUCAA | 2732 |
| 12 | Portia | AGAGUUAUUUAGCUCU-AGUGAUAGAUGUUUGCCCAUCUAAAAAUUAAAGGAUAUCUAUCUGGCGUGC-AUCUGUACAACGUUAGGUGACGUGUAUAGAU------AAAGAAAAUUCACCUCAA | 3040 |
| 13 | M.mirabilis | AGGGCUAUUUAGCCCU-AUUGAUAGAUGUUUGCCCAUCUCAAAAUUAAAGGAUAUCUAUCUGGCGUGC-GAAUGUACAGCGUUAGGUGACGUGUAUGUUC------AAAGAAAAUUCACCUCAA | 3334 |
| 14 | Portia#2 | AGGGUUAUUUAGCCCU-AGUGAUAGAUGUUUGCCCAUCUCAAAAUUAAAGGAUAUCUAUCUGGCGUGC-GAACGUACAGCGUUAGGUGACGUGUAUGUUC------AAAGAAAAUUCACCUCAA | 1828 |
| 15 | Uluborus | AGGGUUAUUUAGCCCU-AGUGAUAGAUGUUUGCCCAUCUUAAAAUUAAAGGAUAUCUAUCUGGCGUGC-GAGCGUACAGCGUUAGGUGACGUGUAUGUUC------AAAGAAAAUUCACCUCAA | 3055 |
| 16 | Myrmarachne | AGGGUUAUUUAGCCCU-AGUGAUAGAUGUUUGCCCAUCUCAAAAUUAAAGGAUAUCUAUCUGGCGUGC-GAGCGUACAGCGUUAGGUGACGUGUAUGUUC------AAAGAAAAUUCACCUCAA | 3065 |
| 17 | H.yanbaruensis | AGAGCUAUUUAGUUCU-AUUGAUCAAUGUUUGCUCACUAUAACAUCAAAGGAUAUUGAUCUGGCGUGC-UGUGCAGUUUGUCUAAGCUAACAGCUGCACG--------AAGAACGAAGCUUGCU | 2591 |
| 18 | P.praticola | AGGGUUAUUUAGCCCU-AGUGAUUAGUGUUUGCCCAUCAAAAGAGAAUAGGAUACUAAUCCGGCGUGCAGUAGUGUGAGU-AUAAGUCUGCCACAUUAUG--------AAGUAAUUGACUUGCA | 3113 |
| 19 | Cybaeus | AGAGUUAUUUAGCUCU-AUUGAUCAGUGUGUGCUCACCC-AAAAUUAUAGGCUACUGAUCCGACGUGCAGAAGUGGAAGA-UUGAGCUGUCUCCCUUUC-------AAAGAAA-UUAGCUUACA | 7518 |
| 20 | E.nubilus | AGGGGUAUUUACUCCU-AUGCCUGAUUGUUUGCCACU-UAAAA--UAUAGGAUAACAGGCCAGUCAAAACUUCUAUUGGG-UUGAGCUGCUCAAUAGUCG---------CAAAUUUAGCUUAGU | 1028 |
| 21 | I.narutomii | AGAGGUAUUUACCUCU-AUGCCAAAUUGUUUGCCACU-UUAAA--UAUAGGAUAAUUGGCCAGUCACAACCUUCGGUGGG-UUAAGCUGCUCAUCGAUUG---------AAAAAUAAGCUUAGU | 2666 |
| 22 | P.yesoensis | AGGGUUAUUUAGCCCU-AUUCAUAAGUGUUUGCCCAACUAAAA-CAAUAGGAUACUUAUGCGCGA--AACCUGCGUAUCGAUUAAGUCGUUGAUACGCGUGG--------AAAAACGACUUGCU | 2772 |
| 23 | Galloisiana | AGGGUUAUUUAGCCCU-AGUGAUAAAUGUUUGCCCACUGUAAAUAUAUAGGAUAUUUAUCCGUCGC-AUUUGUAUAUAUUGCUAAGUUACUGUAUAUAAUUUAAUGUAAAAUAAUUAACUUGCU | 5295 |
| 24 | A.kamurai | AGGGUUAUUUAGCCCU-AGUGAUAAAUGUUUGCCCACUGUAAAUAUAUAGGAUAUUUAUCCGUCGC-AUUUGUAUAUAUUGCUAAGUUACUGUAUAUAAUUUAAUGUAAAAUAAUUAACUUGCU | 2748 |
| 25 | R27-k141 | AGUUGUAUUUACAGCU-AUUGACUAGUGUGUGCGCUCUUGUAAAUAAUAGGCUACUAGUCCAGUCUGC--AGCGCUGAAGGUUAACCUACCUUUAGAUGC-----CAAGUGUAAAUAGGUUAUG | 6834 |
| 26 | O.crucifera | AGGAGUAUUUACUCCU-AUUGGUGCAUGUGUGCCCAUCUAAAA-CAAUAGGCUAUGUACUCGGUGUACGGAGCUGUCAGACUGAGCUUACUGACAAACACGUCU------UUUGACGUGGGCUC | 744 |
| 27 | P.mira | AGGAGUAUUUACUCCU-ACUGGUGCAUGUGUGCCCAUCUAAAA-CAAUAGGCUAUGUACUCGGUGUACGGAGCUGUCAGACUGAGCUUACUGACAAACACGUCU------UUUGA-UGUGGGCU | 3014 |
| 28 | R.ishigakiensis | AGAGGUAUUUACCUCU-AGUGAUGAAUGUGUGCCCACCCUAAA-UAAUAGGCUAUUCAUCCUGCCAAC--CCAAGCGGCUAUUGAGUCAUUGCCGCAUUGG--------UUUAAACGACUCGCU | 3117 |
| 29 | S.rubiginosus | AGAGGUAUUCACCUCU-AUUGGCAAGUGUGUGCCCACCAAAAA-CACUAGGCUACUUGCUCUGCCUGC--AGGCUGUCGGACUGAGUCAUCCGGCGGUC--------GAAGAACUCGACUUACG | 1285 |
| 30 | Sinipoda | AGAGGUAUUCACCUCU-AGUGGCAAGUGUGUGCCCACCGAAAA-CACUAGGCUACUUGCCCUGCCUGC--AGGCUGUCGGACUGAGUCAUCCGGCGGUC--------GAAGAACCCGACUUACG | 416 |
| 31 | Picorna_45cR | AGAGGUAUUUACCUCU-AGUGGCGCAUGUGUGCCCUCCGAAAA-CAAUAGGCUAUGCGCUCGGCGUGCUGGUGCGAGCAGAUUAAGUGGUCCUCGCUCCG-------AAGAAAACUCACUUGCU | 786 |
| 32 | Picorna_251 | AGAGGUAUUUACCUCU-AGUGGUGCAUGUGUGCCCUCCAGAAU-CAAUAGGCUAUGCACCCGGCGUGCUGGUGCGGGAAGAUUAAGUGGUCCUCGCUCCG-------AAGAAAACUCACUUGCU | 1460 |
| 33 | Picorna_155 | AGAGGUAUUUACCUCU-AGUGGUGCAUGUGUGCCCUCCAGAAU-CAAUAGGCUAUGCACUCGGCGUGCUGGUGCGGGAAGAUUAAGUGGUCCCCGCCCCG-------AAGAAAACUCACUUGCU | 1634 |
| | | L2.1 L2.2 L2.3 S3.1 L3.1 L3.2 L3.3 | |

**Table S7.** Sequence alignment of type 6i-2 IRESs, numbered to indicate 5' and 3' terminal nucleotides, structural elements (labelled as in Figure 9C) and conserved nucleotides (indicated by bold font). The conserved UGAUC sequence motif in loop L1.1a and the UGCU motif in loop L1.1b are indicated by blue font, and the conserved motif AGCAAA in loop L1.3 and the ORF2 initiation codon are indicated by red font.

Aligned sequences are from (1) TSA: *Cicurina japonica* IDV#6471 mRNA, IDV6471-W1_S210039725 (GenBank: ICHI01039725.1),(2) TSA: *Iwogumoa* insidiosa IDV#3488 mRNA, IDV3488-W1_S200005522 (IAQB01005522.1), (3) TSA: *Episinus nubilus* IDV#3582 mRNA, IDV3582-W1_S50020893 (IBEU01020893.1), (4) TSA: *Gnathonarium exsiccatum* IDV#6480 mRNA, IDV6480-W1_S20008909 (ICCR01008909.1), (5) TSA: *Doenitzius peniculus* IDV#6479 mRNA, IDV6479-W1_S10007202 (IBJP01007202.1), (6) TSA: *Walckenaeria golovatchi* IDV#6930 mRNA, IDV6930-W1_S250020950 (IBRF01020950.1), (7) TSA: *Pardosa yaginumai* IDV#1014 mRNA, IDV1014-W1_S140008985 (IAGR01008985.1), (8) TSA: *Mallinella sadamotoi* IDV#2667 mRNA, IDV2667-W1_S230029087 (IAKG01029087.1), (9) MAG: Sanya dicistrovirus 2 isolate DFSLS121422 (MZ209830.1),(10) TSA: *Halichondria panicea* TRINITY_DN482871_c0_g1_i1 (GIFJ01725734.1), (11) TSA: *Lariniaria argiopiformis* IDV#1958 mRNA, IDV1958-W1_S210007411 (IATB01007411.1), (12) TSA: *Portia* sp. IDV 2046 IDV#2046 mRNA, IDV2046-W1_S220000814 (IBUW01000814.1), (13) TSA: *Moneta mirabilis* IDV#3751 mRNA, IDV3751-W1_S30000485 (IBWE01000485.1), (14) TSA: *Portia* sp. IDV 2046 IDV#2046 mRNA, IDV2046-W1_S220010416 (IBUW01010416.1), (15) TSA: *Uloborus* sp. IDV 2114 IDV#2114 mRNA, IDV2114-W1_S150016866 (IALG01016866.1), (16) TSA: *Myrmarachne* sp. IDV 5305 IDV#5305 mRNA, IDV5305-W1_S90009092 (IBKO01009092.1), (17) TSA: *Heptathela yanbaruensis* IDV#55 mRNA, IDV55-W1_S90025451 (IAWJ01025451.1), (18) TSA: *Prochora praticola* IDV#3602 mRNA, IDV3602-W1_S200043661 (ICEW01043661.1), (19) TSA: *Cybaeus* sp. IDV 1025 IDV#1025 mRNA, IDV1025-W1_S80002903 (ICCH01002903.1), (20) TSA: *Episinus nubilus* IDV#6266 mRNA, IDV6266-W1_S160021687 (IAEL01021687.1), (21) TSA: *Ischnothyreus narutomii* IDV#4008 mRNA, IDV4008-W1_S160025300 (IBLH01025300.1), (22) TSA: *Piratula yesoensis* IDV#5760 mRNA, IDV5760-W1_S150044920 (IBBE01044920.1), (23) TSA: *Galloisiana yuasai* C216694_a_16_0_l_7829 (GAWN02036639.1), (24) TSA: *Agroeca kamurai* IDV#5497 mRNA, IDV5497-W1_S210028994 (IBIB01028994.1), (25) MAG: Dicistroviridae sp. isolate R27-k141_1120839 (MZ679058.1), (26) TSA: *Orthobula crucifera* IDV#3604 mRNA, IDV3604-W1_S130009465 (IBGQ01009465.1), (27) TSA: *Paikiniana mira* IDV#5507 mRNA, IDV5507-W1_S220023568 (IBUL01023568.1), (28) TSA: *Ryuthela ishigakiensis* IDV#3288 mRNA, IDV3288-W1_S120025746 (IBLL01025746.1), (29) TSA: *Scolopocryptops rubiginosus* strain wildtype C96318_a_4_0_l_1439 (GCIY01020429.1), (30) TSA: *Sinopoda* sp. IDV 944 IDV#944 mRNA, IDV944-W1_S20006649 (IAUT01006649.1), (31) MAG: *Picornaviridae* sp. isolate 45cR-k141_306256 (ON161869.1), (32) MAG: *Picornaviridae* sp. isolate 251-k141_47832 (ON162144.1) and (33) MAG: *Picornaviridae* sp. isolate 251-k141_47832 (ON162144.1).

### Table S8. Sequence alignment of type 6i-3 IRESs.

| # | Viral Sequence | | |
|---|---|---|---|
| | | | P1.1 P1.2 P1.3 SLVI P2.1 |
| 1 | Wenling5 | 5648 | --CAUGGCUAGGGG--UCACAACAAUGGGAU---UAGAGUUGUGUCAGCAGAGCCCGGUCCAUGAGUGU-------------------------------------AAGCAACC |
| 2 | M.alcicornis | 4479 | UAUGAGGCUAGGGG--CCAUCUCACUGUGGG---AAAAGAGAUGACAGCAAAGCCCGGUCCUCAUAUUG--------------------------------------AAAGAGC |
| 3 | Desmophyllum* | 273 | --CGUGGCUAGGGG--CACACAUAAUGAGGU---UCAAGUGUGUGCAGCAAAGCCCGGUCCAUGUGUG--------------------------------------AAGCGACC |
| 4 | PicoQ_sR-025 | 4127 | -AUGUGGCUAGGGG--CACGCGUAAUGAGGU---UGAGAUGCGUGCAGCAGAGCCCGGUCCAUGUGUG--------------------------------------AAGCGACC |
| 5 | Beihai74 | 5170 | -ACGCAGAAUGGG-ACAAUGUUUGAUUGACG---UAUGAAUGUUGUAGCAAAACCUGAAUCUGUGUGC---------------------------------------AAGAGUGG |
| 6 | H.emurai * | 3169 | CCUGACAGAGGCCGUGACGUUAUGGGAG---UCAUGUCACGACAGCAAAGCUCGUUUCAGGAAGG--------------------------------------AAGAGUGG |
| 7 | A.japonicus | 430 | CCCAACAGGGGCUGCCACGUUUUGGGAG---UUAUGUGACAGCAGCAAAGCCCGAUUUGGGAAGG--------------------------------------UAGAGUGG |
| 8 | H.discus* | 286 | UCCUUCUCGAGGGGCUGAUUUCAAAUGGGAG---AAGGAAAUCAGCAGCAAAGCCCGAUGUGAAGGAG--------------------------------------CAGAGUGG |
| 9 | Caledonia4 | 1114 | -GACAGAGCAGGGGUCUUGAGCGAAUGGAGG---AAGGCACAAGACAGCAAAGCCCGAUCUUUGUCGG--------------------------------------UAGAGUGG |
| 10 | Picorna_53b | 1809 | CAACCGAGUAGGGGUUUUGGACAAAUGGAGG---AAGGUCCAGAACAGCAAAGCCCGAUCUCAGUUGG--------------------------------------UAGAGUGG |
| 11 | Wenzhou27 | 5056 | -GACAGUGUAGGGGUCUUGGACAAAUGGAGG---AAGGUCCAAGACAGCAAAGCCCGAUCUCUGUCGG--------------------------------------AAGAGUGG |
| 12 | M.tenta* | 3025 | CUGACCGAGCAGGGGUCCUGGACAAAUGGAGG---AAGGUCCAAGACAGCAAAGCCCGAUCUCGGUCGG--------------------------------------AAGAGUGG |
| 13 | Beihai73 | 6057 | CGAAAGGACAGGGGUCUUGGACAAAUGGAGG---AAGGUCCAAGACAGCAGAGCCCGAUUCCUAUCGG--------------------------------------AAGAGUGG |
| 14 | Aparavirus | 6003 | --GAUCAGACAGGGGUUUUGAACAAAUGGAGG---AAGGUUCAAAACAGCAAAGCCCGAUUCUGGUCGG--------------------------------------AAGAGUGG |
| 15 | Spirobranchus | 3034 | ---CCACGCGUAGGGGUUAAAUUUG-AUCACGA---UAAAAAUUUAACAGCAAAGCCCGAUCGGUGGU---------------CCUUUAUGG-------------------UGAGC |
| 16 | Pseudodiploria | 1054 | --CCUGGCAUAGGGGCACGAACAAAAUUGCCG--UAUAUGUUCGUGCAGCAAAGCCCGUUUCCAGGC------------CACCAUUUUGAUUUGGUG--------------AGUGG |
| 17 | S. pharaonic* | 3792 | -CUUGAGCUAAGGGCUUAGGUCGAAUUGUCG-UAAAUGAUCUAAGCAGCAAAGCCUGGUCUCAAGU--UGUUCUAAUAAAUUAGGAAUUUGAUUCGUUUAUGUAGAAUA--GAACC |
| 18 | Applesnail | 6699 | CCUGAGCUAGGGGCUUAAACUGAAUUGUCG-UAAAGAGUUUAAGCAGCAAAGCCCGGUCUCAGGU--UACCUGUAAAAUUAAGAAGUUAGAUUUGUUUUAUUAGGUA---GAACC |
| 19 | Riboviria1aq | 6525 | ACUAAGGCUAGGGGUUUGAAACAAAUUGUCG-UAAAAGUUUCAUACAGCAAAGCCCGGUCCUUAGU------UACCGAAACAUUCUUAUUUUGAAUUAGUAGGUA------GAACC |
| 20 | C.borealis * | 3295 | CCAAAACUAGGGGCUCUUUUU-AUUCGCCG---UACGAAAAGAGCAGCAAAUCCCGGUCUUUGG----UCUUCGUUCCAUGAAACAGCCCUCAUAGAACAUUCGAAG---GGCGG |
| 21 | Palythoa/QL2018 | 3385 | CCGAAGCUAGGGGCUUCCACU-AAUCGCCG---CAAGGUGGGAGCAGCAAAGCCCGGUCUUCGGU-------------CUUCGUUUUAGAAG----------------GGCGG |
| 22 | Parhyale | 5908 | --CUUGAGCAAGGGGCAAUCUUCU-AUCGCCG---UCGAGGGAUUGCAGCAAAGCCCGGUCUCAAGU-------------AGGAAUUUAUUCCU----------------GGAGG |
| 23 | Farfantepenaeus | 5951 | --CCAUGACAGGGGCUAUCAUCAAAAUCGCCG---AAGGAUGGUAGCAGCAAAGCCCGAUUCAUGGU--------------UGACUUUUGUCA-----------------AGUGG |
| 24 | Beihai72 | 6145 | --CUACAGCUAGGGGCUCUUUCUCAUUCGCCG---UACGAGAAGAGCAGCAAAGCCCGGUCUGUAGU-------AGUAGAGAUGCGUUAUAGUAUCUUUGCU----------GAGGG |
| 25 | Beihai71 | 6126 | --CCAGAGCUAGGGGCUCUCUCU-AAUCGCCG---CAAGGAGGGAGCAGCAAAGUCCGGUCUCUGG--------------UCUUCGUUUUAGA---------------AGGGCGG |
| 26 | Wenzhou26 | 6007 | --CCAGAGCUAGGAGCUCUCUCU-AAUCGCCG---CAAGGAGGGAGCAGCAAAGUCCGGUCUCUGG--------------UCUUCGUUUUAGA---------------AGGGCGG |
| 27 | Stenoteuthis | 83 | --CCAGAGCUAGGGGCUCUCUCU-AAUCGCCG---CAAGGAGGGAGCAGCAAAGCCCGGUCUUUGG--------------UCUUCGUUUUAGA---------------AGGGCGG |
| 28 | Halichondria | 77 | --CCAAAACUAGGGGCUCUUUUU-AUUCGCCG---UACGAAAAGAGCAGCAAAUCCCGGUCUUUGGU---CUUCGUUCCAUGAAACAGCCCUCAUGGGACAUCUGAAG----GGCGG |
| 29 | Cerianthus* | 3295 | --CCAAAACUAGGGGCUCUUUUU-AUUCGCCG---UACGAAAAGAGCAGCAAAUCCCGGUCUUUGGU---CUUCGUUCCAUGAAACAGCCCUCAUAGAACAUUCGAAG----GGCGG |
| 30 | Actinia* | 1690 | --CCGACGCUAGGGGCUCUCUCU-AAUCGCCG---CAAGGAGAGAGCAGCAAAGCCCGGUCGUCGGU-------------CUUCCUUCAGGAAG----------------GGCGG |
| 31 | Beihai70 | 6000 | --CCGAAGCUAGGGGCUCUUUCC-AAUCGCCG---CAAGGAGAGAGCAGCAAAGCCCGGUCUUCGGU--------------CUUCGUUAGAAG-----------------GGCGG |
| 32 | E.coioides* | 3210 | CCGAAGCUAGGGGCUCUUUCU-AAUCGCCG---CAAGGAAAGAGCAGCAAAGCCCGGUCUUCGGU--------------CUUCGUUAGAAG-----------------GGCGG |
| 33 | Stichodactyla | 859 | --CCGAAGCUAGGGGCUUCCAC-UAAUCGCCG---CAAGGUGGAAGCAGCAAUGCCCGGUCUUCGGU-------------CUUCGUUUUAGAAG----------------GGCGG |
| 34 | HPLV-21 | 6241 | -CUUAAGCUUAGGGCUAUGUAGUCAUGGUCAUAACAAUUGCAUAGCAGCAAAUCCCGGACUUAAGU--------------UAGCUUUGCUA------------------GGUGG |
| 35 | Mute15 | 6076 | --CUUAAGCUUAGGGCUAUGUAGUGAUGGUCACAACAAUUGCAUAGCAGCAAAACCCGGACUUAAGU--------------UAGCUUUGCUA------------------GGUGG |
| 36 | S.intermedius | 6189 | --CUAAGGCUUAGGGCAUGUUGGUGAUGGUUGUAACAAUUAACAUGCAGCAAAUCCCGGACCUUAGU------AGGUAAAUCGAUUUGAAAUCUUCUUUUACUU--------AGUGG |
| 37 | Wenzhou28 | 6269 | -ACUUAGGCUUAGGGCAUAGUAGUGAUGGUUGUAACAAUUAUUAUGCAGCAAAUCCCGGACCUAAGU-------AGGUUAAUAGAUUCGUUCUUCCUUAACUU---------AGUGG |
| 38 | P.ibericus | 839 | ---CAAGGAAUGGGGCGCUAUUGUAAUGACAC-UCACGCU-UAGCGCAGCAAUGCCUGAAUCCUUGU-------AUAUUUCUGCUGUACUACCAAAAGAAAUAU---------GGUGG |
| 39 | Rotaria | 182 | --CAGGGAAUGGGACUUUAUUGUGACGACAC-UAACGC-GUAAAGUAGCAAAUCCUCAAUCCCUGU-------AUCUUACUAAGUACUAUCUGACCUAAGAU---------GAAGG |
| 40 | E.kitazawai | 122 | ---CCUAAAAUGGGGUUCUUAAGUAAUGACGGUGAAGAC-UAAGAACAGCAAAGCCUGAAUUUAGGU---------AGAAAGUAAUAUAGUCACUGCUUUUU----------GGUGG |
| 41 | Symphylella* | 342 | ---ACCACCUUAGGGGCUUGGAGCAGAUGACAG---CAGGCUCCAUGCAGCAAAGCCCGAUGGGUGGU----------GAGGUUAAAUCAGUCACUUGAUCUC----------GUGGG |
| 42 | Halichondria2 |  | ---CCAGUGAUAGGGGCUCUGAGUUUAUGGCCG---AUAACACAGAGCAGCAAAGCCCGUUUCCUGGU----AGUUUUGACCUGACCGAAAGGCAGUUACUCAAAACU-----GGUGC |
| | | | L1.1a L1.2a L1.2b L1.3 L1.1b S2.1 |

| | | P1.3 / P2.1 / P2.3 / SLIV / P3.1 / P3.2 | |
|---|---|---|---|
| 1 | Wenling5 | -AGGGUUACUUUGGCCCU-UA-CUAGUGUUCGGGUUGUUAAA--UAUAGGAUACUAG--AUCCCGAGUCCUUUUUUAAGAG-UAACUCGCCUUACUACGGGACU---UUAAUG-AA-UGAGUUGCC | 5831 |
| 2 | M.alcicornis | --AGGUGAAAACCUA-UA-CUAGUGUGUGGCUCUUAAA--UAUAGGCUACUAG-CCCACAAGUGA-UUGCACG-UUU-UGACUCGAUCGUGUAGUCACU----CUAAUG-ACA-GAGUUGCU | 4656 |
| 3 | Desmophyllum* | -AGGGAUAGAACCCUA-UA-CUAGUGUGUGGGUCGGAAAA--UAUAGGCUACUAG-ACCUCAAGUCAUUUGCGCUAGAC-UAACUCAUCGUGCACGUGACU----AUAAUA-AA-UGAGUUGCU | 94 |
| 4 | PicoQ_sR-025 | -GCCAGGAGCAAUCCUA-UA-CUAGUGUGUGGGUCGGAAAA--GAUAGGCUACUAG-ACCUCAAGUCAUUUGCAAUAGAU-UAACUCAUCGUGUACGUGACU----AUAAUG-ACC-GAGUUGCU | 4305 |
| 5 | Beihai74 | -AGGAGUAAUCCUAUUGGUCAGUGUGUGCCACUCUAAA--UAUAGGCUACUGACUCGUCACUUUUACUCC-UGAGUU-UGAUUCGAUUCCGGAGGUUAG----UAAAUG--UUUGAAUCGCU | 5352 |
| 6 | H.emurai | -AGAGCCCAAAGCUCUAUUGCGAGUUGUGUGCCACUUAAA--UCCAGGCUAACUCGCCUCCCUUUUUAGAAAGAAGGUC-UAAUCCGACCUCUGGAGUUUU--GACAAACG-AA-CGGAUUGCU | 2983 |
| 7 | A.japonicus | -AGAGCCGAAGGCUCUAUUGCGAAAUGUGUGCCACUCAAA--UCCAGGCUAUUUCGCCUCCCACUUUUGAAGGAAAGUU-UAAUUCGACUUCCGGAGUUUG--UAUAAACG-AU-AGAAUUGCU | 621 |
| 8 | H.discus | AGACGCGAGUCUACUGAUCGAAGUGUGCCACUUAAA--UCCAGGCUAUCGAUCCUCCCGUUUUAAAGGAGAAGUU-UAACUCAACUUCCCAAGUUUA--CCAAAACUCAC--GAGUUGCG | 100 |
| 9 | Caledonia4 | AGGAGAAUUCCUAUUGCGGGAUGUGUGCCACUGAAA--UCCAGGCUAUUCCGCCCUCCGUUUUUGAGGCUCAGUU-UAACUCAACUGACC-AGGUUUA-CAUAAACC-AA-CGAGUUGCU | 1299 |
| 10 | Picorna_53b | AGGAGAAAUCCUAUUGCGAAAUGUGUGCCACUUAAA--UCCAGGCUAUUUCGCCCUCCGUUUUGAGGCUCAGUU-UAACUCGACUGGCC-AGG-UUUACAAAAACCC--UCGAGUUGCU | 1995 |
| 11 | Wenzhou27 | -AGGAGAAAUCCUAGUGCGAAAUGUGUGCCACUCAAA--UCCAGGCUAUUUCGCCCUCCGUUUUUGAGGAGCAGUU-UAACUCAACUGCCC-AGG-UUUACCGAAACCCAA--GAGUUGCU | 5241 |
| 12 | M.tenta | AGGAGAAAUCCUAUUGCGAAAUGUGUGCCACUCAAA--UCCAGGCUAUUUCGCCCUCCGUUUUUGGGGAGCAGUU-UAACUCGACUG-CCCUGG-UUUACAGAAACCCAA--GAGUUGCA | 2838 |
| 13 | Beihai73 | -AGGAGAAAUCCUAUCGCAGAAUGUGUGCCACUUGAAA--UCCAGGCUAUUCUGCCCUCCGUUUUUGAGGCUCAGUU-UAACUCGACUGGCCA--GGUUUACCGAAACCCAA--GAGUUGCU | 6243 |
| 14 | Aparavirus | -AGGAGCAAUCCUAUUGCAGAAUGUGUGCCACUUGAAA--UCCAGGCUAUUCUGCCCUCCGUUUUUGAGACUCAGUU-UAACUCGACUGG-UCAGG-UUUACCAAAACCCAA--GAGUUGCU | 6188 |
| 15 | Spirobranchus | AGAGAAUAAUUUAUUUCUAUUGGUCAAUGUGUGGCUCAAAAAA--UAUAGGCUAUUGACCUCGUGGGUGUUAAGGAUAGGUU-UGAUCCGACUA-CCUUAA--GCACUUUAAUAU--CUGGAUUGCU | 3226 |
| 16 | Pseudodiploria | -AGAUUGUAUAUAUCUAUUGAUUAAUGUUCGCCACUAUAAA--UAUAGGAUAUUAAUCCGGCACUUUAGAAUGAAGUGGUUUGAUCAUACCCUCAU---AAGUUAGGAAACUCACUAUGAUUGCU | 1258 |
| 17 | S. pharaonis | -AGAAUUUAAAAUUCUAUUGAUUAGUGUGUGGGUUCUAAA--UAUAGGCUACUAAUCCGACAAGUUUUGACCAGGAGAU-UAACCUAUCUUUUGGGGG---UUCUUAAACCAUUUUGGGUUGCU | 3565 |
| 18 | Applesnail | AGGAGUCAUUGUAUUCCUAUUGAUUAGUGUGUGGGUUCUAAA--UAAAGGCUACUAAUCCGACAAGU-UUGACUAGGAGUC-UAACCUGACUCUUAG-GG--GUUCUAAAACCAUUUUAGGUUGCU | 6928 |
| 19 | Riboviria1aq | AGAAAUUAUUUAAUUUCUAUUGAUUAAUGUGUGGGUUCUAAA--UAUAGGCUAUUAAUCCGACAAGUUUUGGCUAGAAGUC-UAACCUGACUUCUAGUGG---UUCUUAAACCAUUUUAGGUUGCU | 6748 |
| 20 | C.borealis * | AGAGUUUAAAGACUCUAUUGGCUGAUGUGUGCCGCCUAAA--UCCUGGCUAUCAGCCCGGCGCUUUUGAUCAGAUCCGUUUAAUCGAGCGACUGAUAG---UUAGAAAACUCAC-UCGAUUGCU | 3073 |
| 21 | Palythoa/QL | AGAGUUUAAAGACUCUACUGGUUGGUGUGUGCCGCCUAAA--UCCAGGCUACCAACCCGGCGCUUUUGACCAGACCUGUUUAAUCGAGCAG-CUGGGAG--UUAGCUAACUCAA-UCGAUUGCU | 3187 |
| 22 | Parhyale | AGCGAAAGCUAUUGGUCAAUGUUUGCCUCCUAAA--UCAAGGAUAUUGACCCGGCGAGUGUAUAAUCAGGCAGAUGAGUUGGUCCCUGAGGAAUGC---UUAAUU-AAAUCAACUGCA | 6100 |
| 23 | Farfantep. | AGAAUUUUCUUGUAAUUUUCUAUUGAUUAGUGUGUGCCACUUUAAA--UAUAGGCUACUAAUCCGGCGAUUUUGGCCAAAAAGUC-UGACUAAGCUUUU-GGU-AG-UUAUUAAACUCAC-UUAGUUGCU | 6151 |
| 24 | Beihai72 | AGCGAAAGCUAUUGAUCAAUGUGUGCCCUCUAAA--UAUAGGCUAUUGAUUCGGCGCGUUUUACUGAGUAGUU-UGACUAAGCCUCAG-GAAGU--ACGAUAACU-AUUUUAGUCGCU | 6351 |
| 25 | Beihai71 | AGAGCUUCGGCUCUACUGGCUGGUGUGUGCCGCCUAAA--UCCAGGCUACCAGCUCGGCGCUUUUGACCAGAUCCGUUGAAUCGAGCGACUGG--GAG-UUAGCUAACUCAA-UCGAUUGCU | 6322 |
| 26 | Wenzhou26 | AGAGUUUCGGCUCUACUGGCUGGUGUGUGCCGCCUAAA--UCCAGGCUACCAGCUCGGCGCUUUUGACCAGAUCCGUUUAAUCGAGCGAC-UGG-GAG-UUAGCUAACUCAA-UCGAUUGCU | 6203 |
| 27 | Sthenoteuthis | AGAGUUUCGGCUCUACUGGCUGGUGUGUGCCGCCUAAA--UCCAGGCUACCAGCUCGGCGCUUUUGACCAGAUCCGUUUAAUCGAGCGAC-UGG-GAG-UUAGCUAACUCAA-UCGAUUGCU | 279 |
| 28 | Halichondria | AGAGUUUAAAGACUCUAUUGGCUGAUGUGUGCCGCCUAAA--UCCUGGCUAUCAGCCCGGCGCUUUAGAUCAGAUUCGUUUAAUCGAGCGAC-UGA-UAG-UUAGAAAACUCA-CUCGAUUGCU | 299 |
| 29 | Cerianthus* | AGAGUUUAAAGACUCUAUUGGCUGAUGUGUGCCGCCUAAA--UCCUGGCUAUCAGCCCGGCGCUUUUGAUCAGAUCCGUUUAAUCGAGCGAC-UGA-UAG-UUAGAAAACUCA-CUCGAUUGCU | 3073 |
| 30 | Actinia* | AGAGUGACGAAAGUAACUCUACUGACUGGUGUGUGCCGCCAUAAA--UCCAGGCUACCAGUCCGGCGCUUUUGAUCAGACCCGUUUAAUCGAGCGG-CUGA-GAG-UUAGCUAACUCAA-UCGAUUGCU | 1488 |
| 31 | Beihai70 | AGAGUUUAGAGACUCUACUGGCUGGUGUGUGCCGCCUAAA--UCCAGGCUACCAGCUCGGCGCUUUUGAUCAGAUCCGUUUAAUCGAGCGG-CUGA-GAG-UUAGCUAACUCAA-UCGAUUGCU | 6196 |
| 32 | E.coioides | AGAGUUUAAAGACUCUACUGGCUGGUGUGUGCCGCCUAAA--UCCAGGCUACCAGCUCGGCGCUUUUGAUCAGAUCCGUUUAAUCGAGCGG-CUGA-GAG-UUAGCUAACUCAA-UCGAUUGCU | 3014 |
| 33 | Stichodactyla* | AGAGUUUAAAGACUCUACUGGCUGGUGUGUGCCGCCUAAA--UCCAGGCUACCAGCCCGGCGCUUUUAAUCAGAUCCGUUUAAUCGAGCGG-CUGA-AAG-UUAGCUAACUCAA-UCGAUUGCU | 661 |
| 34 | HPLV-21 | AGGGGUAUUUACUCCUAGUGAUUGAUGUUUGCCACCUUAAA--UAUAGGAUAUCAAUCUGACGCAACGUUUGUUUGU---UUAAGUUGACAUUCAAAAUUAGUUGUAAAAU--AAUUAACUUGCU | 6439 |
| 35 | Mute15 | AGGGGUAUUUACUCCUAGUGAUUGAUGUUUGCCACCUAAA-UUAUAGGAUAUCAAUCUGACCCAACAUUUAUUUGC---UUAAGUUGACAAAUAAAAUUAGUUGUAAAAC--AAUUAACUUGCU | 6275 |
| 36 | S.intermedius | AGAGUUAUUUAACUCUAUUAAUAAGUGUUUGCCACUCAAAUUUAUAGGAUACUUAUUCAACCCAUCUGUUCUUAGA---UUAAGUUAUCUAAUAACGUAGGAUGUAAAC--AAUCAACUUGCC | 6408 |
| 37 | Wenzhou28 | AGGGUUAUUUAGCUCUAGUAAUAAGUGUUUGCCACUUAAA--UAUAGGAUACUUAUUCAACCCGUCCGGUAUUGGA---UUAAGUUGCCCAAUACUAUCAGACGUAAAC--AAUUGACUUGCU | 6486 |
| 38 | P.ibericus | AGAGCUAUUUAGUUCUAUUCAUCAAUGUUUGCCACCUAAA--UAUAGGAUAUUGAUGGUGUCCGUCUUGUAACUAGA--UUAAUUGUACUACUACUGUUGACG-AAAAAC-AAUUUCAAUUGCU | 1054 |
| 39 | Rotaria | AGGGGUAUUCACCCCUAUUCAUAAAUGUUCGCCUUCUCAAA--UAUAGGAUAUUUAUGGUGUCCAUCCUGUGCCUGA--CCUAAGUUACUCACCACUUUCGAUGU-AAAAUUA-CCUGACUUGCA | 396 |
| 40 | E.kitazawai | AGGGUUAUUUAGCUCUAUUGAUUAAUGUUUGCCACCAUAAA--UAUAGGAUAUUAAUCCCGUCGACCCAGUGCACG---AUUAAGUCGUUGUGUACGUGGUU-----AAAU-----UGACUUGCU | 325 |
| 41 | Symphylella | AGAGCUAUUUAGCUCUAUUGAUGAGUGUUUGCCCACUAAA--UAUAGGAUACUCAUCCUGUCCUUCUAACCAAUUGC--GUGAGUGUGCGUUGGUGUUUGAAGU-AAAAUGU-GUUCACUUGCC | 132 |
| 42 | Halichondria2 | AGAGUUCUCUAGCUCUAUUGAUCAGUGUGUGGCACCUAAA--UAAAGGCUACUGAUCCGGCCCAUCUAACAACCUGU--CUGAGUC-AACCUAUGU-GAUGAUGU-AAAAUAA-CCGACUUCCU | 316 |
| | | L2.1 / L2.2 / L2.3 / L2.4 / S3.1 / L3.1a / L3.1b / L3.2 | |

**Table S8. Sequence alignments of type 6i-3 IRESs.**

Sequence alignment of type 6i-3 IRESs, numbered to indicate 5' and 3' terminal nucleotides, structural elements (labelled as in Figure 9E) and nucleotides conserved in over 85% of members of this group (indicated by bold font). The conserved UGAUC sequence motif in loop L1.1a and the UGCU motif in loop L1.1b are indicated by blue font, and the conserved motif AGCAAAG in loop L1.3 and the ORF2 initiation codon are indicated by red font.

Aligned sequences are from: (1) Wenling picorna-like virus 5 strain WLJQ102796 (NC_032834.1),(2) TSA: *Millepora alcicornis* Seq_81681, transcribed RNA sequence (GKBI01079168.1), (3) TSA: *Desmophyllum pertusum* TRINITY_DN97494_c0_g1_i1 (GHDD01177862.1), (4) Picornavirales Q_sR_OV_025 (KY286107.1), (5) Beihai picorna-like virus 74 strain BHHK130969 (NC_032614.1), (6) TSA: *Hermissenda emurai* breed wildtype BINPACKER.7719.1 (GIDD01085009.1), (7) TSA: *Apostichopus japonicus* Refgene16196_i0_length_675 (GFXQ02045903.1), (8) TSA: *Haliotis discus hannai* breed Genetic and Breeding Research Center TRINITY_DN12135_c0_g2_i1 (GIGJ01006806.1), (9) Caledonia beadlet anemone dicistro-like virus 4 isolate K89 (MF189976.1), (10) MAG: Picornaviridae sp. isolate 53b-k141_597696 (ON161900.1), (11) Wenzhou picorna-like virus 27 strain shrimp14502 (NC_032837.1), (12)TSA: *Macoploma tenta* TRINITY_DN47201_c0_g1_i1 (GJDE01012381.1), (13) Beihai picorna-like virus 73 strain BHTH16077 (KX883415.1), (14) MAG: Aparavirus sp. isolate R107b-k141_128366 (MZ679062.1), (15) TSA: *Spirobranchus lamarcki* comp367760_c0_seq1 (GGGS01138929.1), (16) TSA: *Pseudodiploria strigosa* contig dist251961_c0_seq2 (HACD01215533.1), (17) TSA: *Sepia pharaonis* Unigene40375_All (GEIE01054385.1), (18) Wenzhou channeled applesnail virus 2 strain WZFSL85846 (NC_032877.1), (19) MAG: Riboviria sp. isolate 1aq-RDRP-1 (MW346734.1), (20)TSA: *Cerianthus borealis* contig TRINITY_DN15915_c2_g3_i1 (HAGY01049983.1), (21) TSA: Palythoa sp. QL-2018 isolate Vietnam Unigene120662 (GGUI01160348.1), (22) TSA: *Parhyale hawaiensis* PH.k21.comp15914_seq1 (GFVL01018523.1), (23) TSA: *Farfantepenaeus aztecus* comp47287_c0_seq1 (GEUA01010370.1), (24) Beihai picorna-like virus 72 strain BHHK126587 (NC_032608.1), (25) Beihai picorna-like virus 71 strain BHXG26494 (NC_032562.1), (26) Wenzhou picorna-like virus 26 strain WZSLuoII93478 (NC_033205.1), (27) TSA: *Sthenoteuthis oualaniensis* comp27789_c0_seq5 (GHKH01020532.1), (28) TSA: *Halichondria panicea* TRINITY_DN330973_c0_g1_i1 (GIFJ01655406.1), (29) TSA: *Cerianthus borealis* contig TRINITY_DN15915_c2_g3_i1 (HAGY01049983.1) (30) TSA: Actinia tenebrosa comp54790_c0_seq2 (GEVE01045062.1), (31) Beihai picorna-like virus 70 strain BHJJX25952 (KX883375.1), (32)TSA: *Epinephelus coioides* TRINITY_DN103990_c11_g1 (GHJO01028911.1), (33) TSA: *Stichodactyla helianthus* TR63383c0_g1_i1 (GGNY01161612.1), (34) MAG: Picornavirales sp. isolate HPLV-21 (OM622276.1), (35) Mute swan feces associated picorna-like virus 15 strain Abbotsbury/A/2016 (MW588148.1), (36) TPA_asm: *Strongylocentrotus intermedius* associated picornavirus 1 (BK059604.1), (37) Wenzhou picorna-like virus 28 strain WZRBX42853 (NC_032975.1), (38) TSA: *Proasellus ibericus* contig comp58297_c0_seq2 (HAFA01034871.1), (39) TSA: *Rotaria magnacalcarata* RMAGT0026603 (GDRE01026531.1), (40 TSA: *Episinus kitazawai* IDV#4482 mRNA, IDV4482-W1_S100033950 (IBEG01033950.1), (41) TSA: *Symphylella* sp. AD-2014 strain wildtype C138562_a_5_0_l_491 (GCIH01009984.1), and (42) TSA: Halichondria panicea TRINITY_DN482871_c0_g1_i1 (GIFJ01725734.1).

Table S9. Sequence and structural alignment of chimeric type 6a (Domain 1)/type 6b (domains 2 + 3) IRESs.

| # | Viral sequence | P1.1 P1.2 P1.3 |
|---|---|---|
| 1 | Aparavirus_46450 | 376 ---GUAGAACGAUC-------GUUCUCUCAUAUAAUUGUGAU-AAAUUGAGAGAAGUGCGUUUUAC |
| 2 | Aparavirus_171406 | 269 ----GGAUGUGAUC-------UUGCUCUGUAAUAAUGACAAU-AAUAACAGAAGCAUGCUAUCC |
| 3 | M.occidentalis 1 | 15 ----CGGUGUGAUC-------CGAUCGCGUAAUAUUGGUGC-AAAUACCGCGAUUAGGCUACC |
| 4 | PNG bee virus 2 | 435 AUCAGCAUGUGAUC-------UUGCUGUACUAUAUUGAAAGAGAAAUGUACAGUACUGCUAUGCGAU |
| 5 | Hangzhou 3 | 264 ----GCACGUGAUC-------UUGCUCAGAUAUAUUGAA-GAGAAAUUCUGAGUACUGCUGUGC |
| 6 | M.occidentalis 2 | 321 ----GGAUGUGAUC-------UUGCUACUUUAUAUUGAA-GAGAAAUAAGUAGUAAUGCUAUCC |
| 7 | Thomisidae | 296 ---CGAACGUGAUC-------UUGCUAUAUCAUACUGGCUA-CAAAGAUUUAGUAUUGCUGUUC |
| 8 | Cryptachaea* | 1810 ---AGAGUGUGAUC-------UUGCUAGUUCAUAUUGAUACGAAAGAAUUAGUAUUGCUGCUC |
| 9 | Plasmopara | 333 ----GGAUGUGAUC-------UUGCUAGAUUAUACUGACGA-CAAA-UUCUAGUAAUGCUAUCC |
| 10 | Leptomastix * | 1012 ---AGAAUGUGAUC-------UUGCUCGAAUAUAUUGAUAC-GAAAUUUCGAGUAUUGCUAUUC |
| 11 | Jingmen BDCV 1 | 242 ----GGAUGUGAUC-------UUGCUCAAGAAUAUUGGUCU-GAACA-UUGAGUAUUGCUAUCC |
| 12 | Ellipsidion* | 999 UUCAGCAUGUGAUC-------UUGCUCCCUAAUAUUGACACG-AAUAA-GGAGUAUUGCUAUGC |
| 13 | Red Mite | 437 ----GUGCAUGAUCCCUCCCUGUUCAUAAGCAUAAGUGUC-GAAAGAACAGGUGCACGCUGCAC |
| 14 | Jingmen BDCV 5 | 404 --- GCACGUGACCC-UCCCCAUUCAUAA-CAUAGUUCUC-AAAAGAAUGGGUGCACGCUGUC |
| 15 | D.gallinae * | 3071 --UAGUGCGUGAUCCCUCCCUGUUCAUAAGUAUAAGUGUC-GAAAGAACAGGUGCACGCCGCAC |
| 16 | Pycnopodia | 6427 ---AUUGUGUGAUCUUGCUUGUAACAAUUACUUCCAAUGC-ACUAGUUACUUGCACUGCUACAAU |
| 17 | Beihai91 | 6126 ---UGGAUGUGAUCU---UGUAUGCA-UAUAAUGUAGUCUAGUAAAUUGCAUACACUGCUAUCCA |
| | | L1.1a L1.2a L1.2b L1.1b |

| # | Viral sequence | P1.3 SLVI P2.1 P2.2/L2.1 P2.3 P2.4/L2.3 |
|---|---|---|
| 1 | Apara_46450 | 431 ---GGGUACUCGUACC--UUAAAGGUUCCCUAUUUAGGGAUGGUGUCCUCUCGGCAGCCCCGGAAAA-ACCAAUCACAA |
| 2 | Apara171406 | 321 GAUUGGUACUUGUACCUCUUAAAGGUUCCCUAUUUAGGGAUGGUGACCUCGUGGCAGCCCCACAAAG-ACCAUUUGUCA |
| 3 | M.occidentalis1 | 66 ----------------AAAAUAAGGUGCCCUAUUUAGGGGUGGUCACCUCGGAGCAGCCCUCCAAAA-ACCAAGCACCA |
| 4 | PNG bee virus 2 | 495 -------------------AAAAGGUUCCCUAUUUAGGGAAGGCGACCU-GUGGCGCUCCCACAAAA-GCCAUUCUUUCA |
| 5 | Hangzhou 3 | 316 -----------------AAAUAAGGUUCCCUAUUUAGGGAUGGCGACCGAGUGGCAGCCCCACGAAA-GCCAAUCUUCA |
| 6 | M.occidentalis2 | 373 ----------------AAUUAAAGGUGCUCUUUUUAGAGUUGGCCUCCUCGUGGCAGCCCCACAAAA-GCCAAUCUUCAA |
| 7 | Thomisidae | 349 ----------------AACAACGAGUGCUCUAUUUAGAGAUGGCCCCUCACUGGCAGCCCCAGAAAA-GCCCUUAGCCA |
| 8 | Cryptachaea * | 1757 -----------------AAUUUUGGUGCUCUUUUUAGAGAUGGUCUCCACGCAGCAGCCCUGCAAAA-ACCAUGUAGCCA |
| 9 | Plasmopara | 384 ----------------AACUACAAGAGCCCUAUUUAGGGAUGGCCUCUUCGCAGCAGCCCUGCAAACUGCCACUCGUCA |
| 10 | Leptomastix * | 959 ----------------AACUACAGGUUCUCUAUUCAGAGAUGGCGUCCUAGCAGCAGCCCUGCAAAA-GCCAAGUAUCA |
| 11 | Jingmen BDCV 1 | 293 ------------------AAAUAGGUGCCCUAUUCAGGGAUGGUCUCCUCGUGGCAGCCCCACAAAA-ACCUUGACCA |
| 12 | Ellipsidion * | 933 ----------------AAUAACGAGAGCACUAUUUAGUGAUGGCCUCUCACCAGCAGCCCUGGAAAA-GCCAUGUGUCAA |
| 13 | Red Mite | 496 ------------------GAACCGGCUUCCUAUUCAGGGUGAGUGUGCCAAGAGCAGCCCUCUAUA-CACUCAGACACUU |
| 14 | Jingmen BDCV 5 | 461 -----------------GAACAUGGCUUCCUAUUUAGGGAAGGUGUGCCAGGAGCAGCCCUCUAUACACCUAGAGAACC |
| 15 | D.gallinae * | 3010 ------------------GAACCGGCUUCCUAUUCAGGGUGAGUGUGCCAGGAGCAGCCCUCCAUA-CACUCAGACACU |
| 16 | Pycnopodia | 6488 ---------------GAGUUUUUGGGUUGCUAUUUAGCGAAGGUGUCCCAGGAGCAGCCCUCCAUA-CACCGCAUAGGA |
| 17 | Beihai91 | 6184 ----------------GCUUUUAGGUCCCCUAUUUAGGGAGAGCGUCCUGCGAGCAGCCCUCCAAAAGCUCAAUAGACU |
| | | S2.1. S2.2 L2.1 L2.2 L2.3 L2.4 |

| # | Viral sequence | P3.3 / P3.1/P3.2 / SLIII (P3.4/L3.2) | |
|---|---|---|---|
| 1 | Apara_46450 | -CACUUUCAUUCUUGCCUGGGUCGGUAAGAAA-CAUAUGCGAUCAUAUCAAGUAUGGUGAAUUUUAA-UUUGACCUGCU | 579 |
| 2 | Apara171406 | -CAC-GACGUGUCCGUCUAGGUCGGCGGACAA-CAUAUGCGAUCAUAUCAAGUAUGGCGUUGCUUAAUUUUGACCUGCU | 474 |
| 3 | M.occidentalis1 | -AGGUUCGAUGCCUUGCUGGGUGGCAAGACAA-----CUUUCUUUCGAGAAAG---GUCGAAUACAAACAUCACCCGCU | 197 |
| 4 | PNG bee virus 2 | --CCCACCUCCUGAGCUUGAGUCAGCAAAGGAA----CGAGAAAUUUUCUCG----CUGGUGGUUAA-CUAGACUUGCU | 621 |
| 5 | Hangzhou 3 | CCCUCACUUGUCUGCGAUGGGCAUCAAGGACAA----AUACCUUUUGGUAU-----UAGUGUUGUAAUAACUGCCUGCU | 446 |
| 6 | M.occidentalis2 | -CCUCGCUUGUCUGCGAUGGGCAUCAAAGACAA-----CAUCCUCGGGUG------UAGCGUUGUAACAACUGCCUGCU | 502 |
| 7 | Thomisidae | -AACUGCAUUGUUACAAUGGGUGUUGUAACAA----CAUUUCUUCGGAAUUG----GAUGCUUUUAAG-UUCACCUGCU | 479 |
| 8 | Cryptachaea* | -ACCUUCCUCGUUACUAUGGGUGUAGUAACAACA-----UCGUUUCGA----CUUUGAAGGAAGUAAA-UUCACCUGCU | 1629 |
| 9 | Plasmopara | -ACCUGCCUCGACUCUAUGGGUGUAGAGUCAA----CAUUCCUUCGGGAAU----GCAAGCUUGUAA-CCUCACCUGCU | 515 |
| 10 | Leptomastix* | -ACCUCUCCCGACUUUAUGGGUGUAAAGUCAA-----CUAUUUUCGAAUAG-----AGGAGUUGUAA-CCCCACCCGCU | 831 |
| 11 | Jingmen BDCV 1 | CUUGCCAUCGACCUAUGGGUGUAAGGAAGAG------CUACAAAGUAG--------UAACGGCAAAAUUCACCUAAC | 414 |
| 12 | Ellipsidion* | --CCCUUGCCGAAGUUAUGGGUGUAACUUCAA----CAUUUCCACGGAAUUG-----AGCAAGGUAC-CACCCCCCGCC | 804 |
| 13 | Red Mite | --GCGGCCCAGCCAUAUUAGGUCAUGUUGGCAG--CCAGGAUUUCGAUUCUGG-UGGGCUUUAAUAC--CCGACCUGCU | 628 |
| 14 | Jingmen BDCV 5 | -CGCGCAUCAACCACUGUAGGUCCAGUUGGUAG----CUUCGCUUGCGGAG---UGAUGUUUAAUAU--UUGACCUGCU | 590 |
| 15 | D.gallinae * | -CGCGGCCCAGUCACAUUAGGUCAUGUUGACAG--CCAGGAUUUCGAUUCUGG-UGGGCUUUAAUAC--CCGACCUGCU | 2878 |
| 16 | Pycnopodia | -ACCUCCCCAGUCACAUUGGGUUAUGUUGACAG----CUGCGUUUACGCAG-----UGGGGUAGUAU-UACAACCUGCU | 6618 |
| 17 | Beihai91 | --UCGGAGCAGCCAUCCUGGGUUAGAUCGGCAGC---CAUGUUUACAUG-------GUGCUCUUAAAAAUUAACCUGCU | 6313 |
| | | S3.1 L3.1a. L3.1b L3.2 L3.3 | |

Table S9. Sequence alignment of chimeric type 6a (Domain 1)/type 6b (domains 2 + 3) IRESs, numbered to indicate 5’ and 3’ terminal nucleotides and annotated to show structural elements (as shown in Fig. 7A) and the initiation codon (red font). Bold font indicates nucleotides that are conserved in over 75% of members of this group. Nucleotides that are predicted to engage in base-pairing in helical elements P1.1, P1.2, P1.3, P2.1, P2.2, P2.3, P2.4, P3.1, P3.2 and P3.3 are shaded and bracketed. Note that sequences # 8, 10, 12 and 15 (indicated by an asterisk) are numbered according to the deposited (antisense) sequences.

Aligned sequences are from: (1) MAG: Aparavirus sp. isolate 220-k141_46450 (GenBank: MZ679093.1), (2) MAG: Aparavirus sp. isolate 220-k141_171406 **(**MZ679071.1), (3), TSA: *Metaseiulus occidentalis* GQ6VMKX02HKOT3 (JL052943.1), (4) PNG bee virus 2 isolate PNGBV/2 (MT482484.1), (5) Hangzhou dicistrovirus 3 isolate HDV3/FYBPH-2 (OM751363.1), (6) TSA: *Metaseiulus occidentalis* MITE_454Assem.sd.12189.C1 (JL020592.1), (7) TSA: *Thomisidae* gen. sp. IDV 5033 IDV#5033 (ICFX01018932.1), (8) TSA: *Cryptachaea gigantipes* IDV#6848 (IBTR01011776.1), (9) *Plasmopara viticola* lesion associated dicistro-like virus 1 isolate DMG-E-DN28538 (MN551111.1), (10) TSA: *Leptomastix dactylopii* s5308_L_19038_0_a_20_5_l_1112 (GBNE01033170.1), (11) MAG: Jingmen bat dicistrovirus 1 isolate JM_Mi.schreibersii_dicistro_1 (OQ715861.1), (12) TSA: *Ellipsidion* sp. AD-2015 breed wildtype C391187_a_7_0_l_1351, transcribed RNA sequence (GDYE01061832.1), (13) Red mite dicistrovirus 1 isolate RMDIV1 (ON160029.1), (14) MAG: Jingmen bat dicistrovirus 5 isolate JM_My.chinensis_dicistro_3 (OQ715869.1), (15) TSA: *Dermanyssus gallinae* isotig17344 (GAIF01027790.1), (16) TPA_asm: *Pycnopodia helianthoides* associated picornavirus 2 (BK059603.1), and (17) Beihai picorna-like virus 91 strain BHJJX25930 (NC_032638.1).

Table S10. Sequence and structural alignment of type 6a IRESs located within the coding sequence of dicistrovirus genomes.

| # | Viral sequence | | P1.1 · L1.1a · P1.2 · P1.3 · L1.1b |
|---|---|---|---|
| 1 | Leptomastix | 3067 | UGCUCCGAUGUGAUCUUUGAUAACGC-AACUAACAAGCGAGGUUUUAAGUGUUUUCAA--UUGCUAUCAGGAGUA |
| 2 | Anagyrus | 5596 | UGCUCCGAUGUGAUCUUUGAAAACG-AUAGUAACAAGCGAGGUUUUAAACGUUUUCAA--CUGCUAUCUGGAGUA |
| 3 | P.pennipes | 137 | UCUGAUGAUGUGAUCUUUGAUCGGAACAUGCAA-AAGCAAGGACAAAAGUCCGAUUA--UAUGCUAUCAAUCAGG |
| 4 | T.clavata | 3017 | UCUGAUGAUGUGAUCUUUAAUCGGGACAUGCAA-AUGCAAGGACAAAACUCCGAUUA--CAUGCUAUCAAUCAGG |
| 5 | P.puparum | 6000 | UCUGAUGAUGUGAUCUUUAAUCGGGACAUGCAA-AUGCAAGGACAAAACUCCGAUUA--CAUGCUAUCAAUCAGG |
| 6 | Soybean thrips | 4929 | UCCUCGGAUGUGAUCUUUGACGAAAA-GGAAAACAUGCAAGGCCCUAAUUUGCGUCAA--CUGCUAUCUCGAGGA |
| 7 | A.abdominalis | 5846 | UCCUCAGAUGUGAUCUUUGACGAAAA-GGAAAACAUGCAAGGCCCUAAUUUGCGUCAA--UUGCUAUCUCGAGGA |
| 8 | Diversinervus | 5754 | UCUCAAGAUGUGAUCUUUGACGAAAA-GGAAAACAUGCAAGGCCCUAAUUUGCGUCAA--UUGCUAUCGAGAGA |
| 9 | P.ceylonica | 1873 | UCUUUAGAUGUGAUCUUUGACGAGAA-GGAAAACAUGCAAGGCCCUAAUUUGCGUCAA--CUGCUAUCUAGAGA |
| 10 | PSIV | 6005 | ---GACUAUGUGAUCUUAUUAAAAUUAGGUUAAAUUUCGAGGUUAAAAAUAGUUUUAAUAUUGCUAUAGUC |
| 11 | Himetobi | 6286 | GAAAAUGUGUGAUCUG-AUUAGAAGUAAGAAAAUU-CCUAGUUAUAAUAUUUUUAAUA-CUGCUACAUUUUU |
| 12 | H.minitabunda | 6062 | GUUUGGGAAAGAUCUGUAUUUGAUCAGGCAAAUAGUUCUGUUAAAAAUGAU-CAAAUAGUUGCUACCCAGAU |
| 13 | Bat3 | 1496 | GUUUGGGAAAGAUCUGUAUUUGAUCAGGCAAAUAGUUCUGUUAAAAAUGAU-CAAAUAGUUGCUACUCAAAU |
| 14 | Serbia | 6081 | GUUUGGGAAAGAUCUGUAUUCGAUCAGGCAAAUUGUCCUGUCAUAAAUGAA-CGAAUAGUUGCUACCCAAAU |
| 15 | Hungary | 6090 | GUUUGGGAAAGAUCUGUAUUCGAUCAGGCAAAUUGUCCUGUCAUAAAUGAA-CGAAUAGUUGCUACCCAAAU |
| 16 | Army_ant3 | 5455 | GUUUGGGAAAGAUCUGUAUUCGACCAGGCAAAUUGUCCUGUCAAAAAUGGA-CGAAUAGUUGCUCCCCAAAU |
| 17 | A.spinosus | 6094 | GUUUGGGAAAGAUCUGUAUUCGACCAGGCAAAUUGUCCUGUCAAAAAUGGA-CGAAUAGUUGCUCCCCAAAU |
| 18 | Triato-Vvel | 1924 | GUUUGGGAAAGAUCUGUAUUCGAUCAGGCAAAUUGUCCUGUCAAAAAUGAA-CGAAUAGUUGCUACCCAAAU |
| 19 | V.amethystina | 118 | GUUUGGGAAAGAUCUGUAUUCGACCAGGCAAAUUGUCCUGUCAAAAAUGGA-CGAAUAGUUGCUACCCAAAU |
| 20 | Vespula | 6094 | GUUUGGGAAAGAUCUGUAUUCGACCAGGCAAAUUGUCCUGUCAAAAAUGGA-CGAAUAGUUGCUCCCCAAAU |
| | | | L1.1a L1.2a L1.2b L1.1b |

| # | Viral sequence | | P1.3 · P2.1 · P2.2/L2.1 · P2.3 · P2.4/L2.3 |
|---|---|---|---|
| 1 | Leptomastix | 2995 | GAGACCGGGAAUAUUUAUUCUUACCACACAGGAUCGGUCCAGGCAAUCCCUGAAAAUCCAGUGACCCUCG- c U |
| 2 | Anagyrus | 5668 | AAGACCGGGAUAUUUAUCCUUACCACACAGGAUCGGUCCAGGCAAACCCUGAUUAUCCAGUGACCCUCG-CU |
| 3 | P.pennipes | 209 | AAAGCUGUGGAUAUUUAUCCUUACCUCACAGGAUCAGCUCAGGCAGCCCCGAAAAUCCAGUGUCCCUUG-CU |
| 4 | T.clavata | 2945 | AAAGCUGUGGAUAUUUAUCCUUACCUCACAGGAUCAGCUCAGGCAGCCCCGAAAAUCCAGUGUCCCUUG-CU |
| 5 | P.puparum | 6072 | AAAGCUGUGGAUAUUUAUCCUUACCUCACAGGAUCAGCUCAGGCAGCCCCGAAAAUCCAGUGUCCCUUG-CU |
| 6 | Soybean thrips | 5001 | AAAGCGGUGGGUAUUUACCCUUACCACACAGGAUCCGCUCAGGCAGCCCCGAAACAUCCAGUGAACCUUG-CU |
| 7 | A.abdominalis | 5918 | AAAGCGGUGGGUAUUUACCCUUACCACACAGGAUCCGCUCAGGCAGCCCCGAAACAUCCAGUGAACCUUG-CU |
| 8 | Diversinervus | 5825 | AGAAGCGGUGGGUAUUUACCCUUACUACACAGGAUCCGCUCAGGCAGCCCCGAAACAUCCAGUGAACCUUG-CU |
| 9 | P.ceylonica | 1944 | AGAAGCGGUGGGUAUUUACCCUUACCACACAGGAUCCGCUCAGGCAGCCCCGAAAUAUCCAGUGACCCUUG-CU |
| 10 | PSIV | 6073 | UUAGAGGUCUUGUAUAUUUAUACUUACCACACAAGAUGGACCGGAGCAGCCCUCCAAUAUCUAGUGUACCCUCG |
| 11 | Himetobi | 6355 | AAGACCCUUAGUUAUUUAGCUUUACCGCCCAGGAUGGGGUGCAGCGUUCCUGCAAUAUCCAGGGCACCUAGG-U |
| 12 | H.minitabunda | 6133 | GACGGGUGUUUGUAUUUACAAUUACCUUCCAGGAUCACCCGUGGCAGUCCCACAUUAUCCAGGGAAGCAGAACU |
| 13 | Bat3 | 1567 | GACGGGUGUUUGUAUUUACAAUUACUUUCCAGGAUCACCCGUGGCAGUCCCACAUUAUCCAGGGAGGCAGAACU |
| 14 | Serbia | 6152 | GACGAGCGGUCGUUUUUACGAUUACCUUCCAGGAUCGCUCGAGGCAGCCCCUCAAUAUCCAGGGAAACAGGACU |
| 15 | Hungary | 6161 | GACGAGCGGUCGUUUUUACGAUUACCUUCCAGGAUCGCUCGAGGCAGCCCCUCAAUAUCCAGGGAAACAGGACU |
| 16 | Army_ant3 | 5526 | GACGAGCGUUUGUAUUUACAAUUACCUUCCAAGAUCGCUCGAGGCAGCCCCUCAAUAUCUAGGGAAACAGGACU |
| 17 | A.spinosus | 6165 | GACGGGCGUUUGUAUUUACAAUUACCUUCCAAGAUCGCUCGAGGCAGCCCCUCAAUAUCUAGGGAAACAGGACU |
| 18 | Triato-Vvel | 1995 | GACGAGCGUUUGUAUUUACAAUUACCUUCCAAGAUCGCUCGAGGCAGCCCCUCAAUAUCUAGGGAAACAGGACU |
| 19 | V.amethystina | 189 | GACGAGCGUUUGUAUUUACAAUUACCUUCCAAGAUCGCUCGAGGCAGCCCCUCAAUAUCUAGGGAAACAGGACU |
| 20 | Vespula | 6165 | GACGGGCGUUUGUAUUUACAAUUACCUUCCAAGAUCGCUCGAGGCAGCCCCUCAAUAUCUAGGGAAACAGGACU |
| | | | S2.1 L2.1 L2.2 L2.3 L2.4 |

| # | Viral sequence | | P3.1/P3.2 P3.3 | |
|---|---|---|---|---|
| 1 | Leptomastix | 2923 | CGGCAGCGAAGAGAAGGUGCUCUCGUUGCUCAAAGUUUUUCACCUGCU | 2876 |
| 2 | Anagyrus | 5740 | CGGCAGCGAAGAGAAGGUGCUCUCGUUGCUCAAAGUUUUUCACCUGCU | 5787 |
| 3 | P.pennipes | 281 | AGGCUUCAAAGAGAAGGUGCUCUAGAAGCUGAAUCCAACUCACCUGCU | 328 |
| 4 | T.clavata | 2875 | AGGCUUCGAAGAGAAGGUGCUCUAGAAGCUGAAUCCAACUCACCUGCU | 2826 |
| 5 | P.puparum | 6144 | AGGCUUCGAAGAGAAGGUGCUCUAGAAGCUGAAUCCAACUCACCUGCU | 6191 |
| 6 | Soybean thrips | 5073 | AGGCCUCGAAGAGAAGGUGCUCUAGAGGCUGAAGUAUUCUCACCUGCU | 5120 |
| 7 | A.abdominalis | 5990 | AAGUCUCGAAGAGAAGGUGCUCUAGAGGCUGAAAUAUUUUCACCUGCC | 6037 |
| 8 | Diversinervus | 5898 | AGGCUUCGAAGAGUAGGUGCUCUAGAAGCUGAAGUUUUUACACCUGCU | 5945 |
| 9 | P.ceylonica | 2017 | AGGCUUCGAAGAGAAGGUGCUCUAGAGGCUGAAAUGUUCUCACCUGCU | 2064 |
| 10 | PSIV | 6147 | GCUCGCUCAAACAUUAAGUGGUGUUGUGCGAAAAG-AAUCUCACUUCAA | 6195 |
| 11 | Himetobi | 6428 | GCAGCCUUGUAGUUUUAGUGGACUUUAGGCUAAAG-AAUUUCACUAGCA | 6475 |
| 12 | H.minitabunda | 6207 | CGAUCAAAAAGAUUAAGUGGUCUUCAGAUUUAAA-UAACUCACUUGCA | 6253 |
| 13 | Bat3 | 1641 | CGAUCAAAAAGAUUAAGUGGUCUUUAGAUUUAAA-UAACUCACUUGCA | 1687 |
| 14 | Serbia | 6225 | CGCUUGAAAAGUUUAAGUGGACUUUAAAGUUAAA-UAACUCACUUGCA | 6272 |
| 15 | Hungary | 6235 | CGCUUGAAAAGUUUAGGUGGACUUUAAAGUUAAA-UAACUCACUUGCA | 6281 |
| 16 | Army_ant3 | 5600 | CGUUCGAAAAGUUUAAGUGGACUUUAGAAUUAAA-UAACUCACUUGCA | 5646 |
| 17 | A.spinosus | 6239 | CGUUCGAAAAGUUUAAGUGGACUUCAGAACUAAA-GAACUCACUUGCA | 6285 |
| 18 | Triato-Vvel | 2069 | CGUCCGAAGGGUUUAAGUGGACCUUAGGAUUAAA-AGACUCACUUGCA | 2115 |
| 19 | V.amethystina | 263 | CGUCUGAAAGGUUUAAGUGGACCUUAGGAUUAAA-GAAUUCACUUGCA | 309 |
| 20 | Vespula | 6239 | CGUUCGAAAAGUUUAAGUGGACUUCAGAACUAAA-GAACUCACUUGCA | 6285 |
| | | S3.1 | L3.1 L3.3 | |

Table S10. Sequence alignment of type 6a IRESs located within the coding sequence of monocistronic dicistrovirus sequences, numbered to indicate 5' and 3' terminal nucleotides and annotated to show structural elements (as shown in Fig.10B), the initiation codon (red font) and nucleotides conserved in 85% or more of members of this group (bold). Nucleotides that are predicted to engage in base-pairing in helical elements P1.1, P1.2, P1.3, P2.1, P2.3, P2.4, P3.1 and P3.2 are shaded and bracketed. Note that sequences # 1 and 4 are numbered according to the deposited (antisense) sequences. For comparison, the alignment includes the canonical *Plautia stali* intestine virus and Himetobi P virus type 6a IRESs. Aligned sequences are from: (1) TSA: *Leptomastix dactylopii* s2994_L_5767_0_a_65_0_l_8705 (GBNE01030609.1), (2) *Anagyrus vladimiri* dicistrovirus (OP349062.1), (3) TSA: *Platycnemis pennipes* breed wildtype C61823_a_7_0_l_511 (GCDY01011496.1), (4) TSA: *Trichonephila clavata* IDV#4683 mRNA, IDV4683-W1_S90002957 (ICKC01002957.1), (5) TSA: *Pteromalus puparum* comp28542_c0 (GECT01016007.1, (6) Soybean thrips picorna-like virus 1 strain STN1 (MT240797.1), (7) TSA: *Aphelinus abdominalis* C129619_a_61_0_l_8944: (GBTK01008700.1), (8) *Diversinervus elegans* virus (MT127560.1), (9) TSA: *Phrynarachne ceylonica* IDV#5661 (IAYL01036549.1), (10) *Plautia stali* intestine virus (NC_003779.1), (11) Himetobi P virus (NC_003782.1), (12) TSA: *Helpis minitabunda* IDV#6686 mRNA, IDV6686-W1_S70011807 (IAUM01011807.1), (13) MAG: Bat faecal associated dicistrovirus 3 isolate G02/aus/1 (ON872536.1), (14) Serbia picorna-like virus 1 isolate 71282 (MT822187.1), (15) Cripavirus NB-1/2011/HUN (KJ802403.1), (16) MAG: Army ant associated dicistrovirus 3 isolate MGN-3_G04 (OP884011.1), (17) TSA: *Alysson spinosus* C154488_a_35_0_l_3588 ( GBUA01012649.1) (18) MAG: Triatovirus sp. isolate Triato-Vvel (OR416408.1), (19) TSA: *Vestalis amethystina* breed wildtype C86215_a_5_0_l_428 (GCLJ01028224.1), (20) *Vespula vulgaris* Picorna-like virus 2 isolate Vvul_virus_09 (MZ443577.1).

Table S11. Sequence alignment of type 6h IRESs.

| | | | P1.3; P1.1; P1.2; P2.1; P2.2/L2.1 |
|---|---|---|---|
| 1 | Robinvale virus | 294 | --GAUUAAUUUCUUAAUGUGCGUGGUGGCACCUUUAAAUAGU-AGAUUUCCGUCUUUCUCUAUUCAGAGUACUUUUGGACGU--------UAACUAAUGCUCCAUC |
| 2 | L.taurica virus | 294 | --GAUUAAUUUCUUAAUGUGCAUGGUGGCACCUUUAAUAAGUA-GAUUUCCGUCAUUCUCUAUUUAGAGUACUUUUGGACGU--------UAACUAAUGCUCCAUU |
| 3 | Rice-curl virus | 277 | ------CGCUAUUUAUAGGAAGCGAAGAAUUUUCUAUUUAUAG-UCGUAGUGUCA-UUCCUAUUUAGGAUACGGUUGGACACA---UAAAUAACUAAU-CUUCUUC |
| 4 | T.fruiticans | 287 | ------CGCUAUUUAUAGGAAGCGAAGAAUUUUCUAUUUAUAG-CCGUAGUGUCA-UUCCUAUUUAGGAUACGGUUGGACACA---UAAAUAACUAAU-CUUCUUC |
| 5 | Picorna-107 | 262 | ------CGCUAUUUAUAGGAAGCGAAGAAUUUUCUAUUUAUAG-CCGUAGUGUCA-UUCCUAUUUAGGAUACGGUUGGACACA---UAAAUAACUAAU-CUUCUUC |
| 6 | Maize picornav | 6510 | ------CGUUAUUUAUAGGAAGCGAAGAAUUUUCUAU-AGAUAAUCGUAGUGUCA-UUCCUAUUUAGGAUACGGUUGGACACA---UUAAUAACUAAU-CUUCUUC |
| 7 | Stethorus | 229 | ----ACCCUUUAUUUCAUGAAGCGAGAAAUUUUAUGA--UGUAAUUGUAGUGUCA-UUCCUAUUUAGGAUACGGUUGGACACA---UUAAUAACUAAU-CUUUCUC |
| 8 | A.glycines | 385 | ----ACCUCUUUCGUUUUGAAGCGUGGAAUUUCAA-UUCGAUUGUAGUAGUGUCA-UUCCUAUUUAGGAUACGGUUGGACACA---UAAAUAACUAAU-CUUCCGC |
| 9 | Y.stiatipes | 8282 | ------------------------GGAGUUUUUAA--GCGAAAGUAGUAGUGUCA-UUCCUAUUUAGGAUACGGUUGGACACA---UUAAUAACUAAU-CCUCCAC |
| 11 | Tu-picornav1 | 280 | ----ACCUCUUUCGUCUUGAAGCGUGGAGUUUCAAGUGAAU--GUAGUAGUGUCA--UCCUAUUCAGGAUACGGUUGGACACG---UAAAUAACUAAU-CCUCCAC |
| 11 | Acanthasoma | 223 | ----ACCUCUUUUGUCUUGAAGCGUGGAGUUUUAAGCAAAA--GUAGUAGUGUCAUUUCCUAUUUAGGAUACGGUUGGACACA---UAAAUAACUAAU-CCUCCGC |
| 12 | Picorna-193 | 292 | ------AUCUAUCUAGUUGGAGCGGGAAGUUUUGACUAGAU--UUGAUAUCGUCA-UUCCUAUUUAGGAUACGGUUGGACGAGAUUAAAAUAACUAAC-CCUUCCC |
| 13 | Picorna-238 | 286 | ------AUCUAUCUAGUUGGAGCGGGAAGUUUUGACUAGAU--UUGAUAUCGUCA-UUCCUAUUUAGGAUACGGUUGGACGAGAUUAAAAUAACUAAC-CCUUCCC |
| 14 | Cherry_CVTA | 97 | ---------AUUGUAAUUGAAGCGGGGAGUUUUGAUUGCAA---UUGUUUCGUCA-UUCCUAUUUAGGAUACGGUUGGACG--------UAAACUAAC-CCUCCCC |
| 15 | Cherry-PAI2 | 98 | ---------AUUGUAAUUGAAGCGGGGAGUUUUGAUUGCAA---UUGUAUAGUCA-UUCCUAUUUAGGAUACGGUUGGACU--------UUAACUAAU-CCUCCCC |
| 16 | Sparassidae | 2663 | -------UCAUUUCAAUUGGAGCGGGAAAUUCCGAUUGAUUUUGA--UAUCGUCA-UUCCUAUUUAGGAUACGGUUGGACGAGUUUAAUAUAACUAAU-CUUUCCC |
| 17 | C.amaricus | 8217 | ------AUUGAAUAUAUCUAAUCAUGUGGAUAAUAUAUUCAAUAAA--UCCGUCA-UUCCUAUUUAGGAUAACAGUGGACG--------UUAACUAAU-GCCACAU |
| 18 | A.lateralis | 8362 | -CUCUACUAUAUAUGUGUUAAGCAUCGGGAUUAUGAUAUAGUU-AGCUCACUGGG-UUCCUAUUUAGGAUACGGUUGCCAGUA----UAAAAUCCAAU-CCCCGGU |
| 19 | P.punctatus | 8381 | ACUCUAUCACAUAUUUGUUAAGCGUCGAAAUUAUGAUGUGGUU-AGUUCACUGGG-UUCCUAUUUAGGAUACGGUUGCCAGUA----UAAAAUCCAAU-CUUCGGC |
| 20 | E.necator virus | 233 | ----GAUGGUCAA-CUAGUGAGCGUGGGGUUCACUAUUGAUU-UC--AUAAGACUUUCUCUAUUUAGAGUACGGUUGGUCUA--------UAACUAAU-CCCCCGC |
| 21 | B.kamakur. | 8431 | ------ACAUAGAAAUGUUAAGCAUUGGGAUUAUGUCUAUGU---UGAAGUAGCG-UUCCUAUUUAGGAUACGGUUGGCUAUU----AAAAUAUCAAU-CCCCGGU |
| 22 | Corrinnidae | 197 | -----GAAGAUUCAGGUAGGAGCGUCUAAAUUCCCCUGACUAGUC--CUCGAGCA-UUCCUAUUUAGGAUACGGUUGGCUUUA------ACUACUAAU-CUUAGAC |
| 23 | C.formosus | 326 | -----GAAGUUUCAGGUAGGAGCGCCUAAAUUCCCCUGACUAGUC--CUCGAGCA-UUCCUAUUUAGGAUACGGUUGGCUUUA------AUUACUAAU-CUUAGGC |
| 24 | C.wygodzinskyi | 179 | ------ACUA-CAUUGGUAGGAGCAAGUGGAUUCCCCAAUGUUUAGUGGUCACGCA-UUCCUAUUUAGGAUACGGUUGGCGUAU--------UACUAAU-CCCACUU |
| 25 | E.hageni | 1109 | ------CUUUAUCGAGUAAGAGCAGACAGAUUCUCUUGAUAAAG-UUUCCACGCA-UUCCUAUUUAGGAUACGGUUGGCGUA--------UUACUAAU-CCUGUCU |
| 26 | C.kitazawai | 383 | ---CACCUAUUUAAAGUAAGAGCCAGUGGAUUCUCUUUACUUAGUG-GCUUCGCA-UUCCUAUUUAGGAUACGGUUGGCGUA--------UUACUAAU-CCCACUG |
| 27 | Philodromus | 8386 | ---CACCUAUUUAAAGUAAGAGCCAGUGGAUUCUCUUUAUCUAGUG-ACUUCGCA-UUCCUAUUUAGGAUACGGUUGGCGUA--------UUACUAAU-CCCACUG |
| 28 | M.pseudojobi | 54 | --------CAUAUAUAUUUAAGCGCGUAAUUUAUUUAUA--UAUGUGAUCUAGCA-UUCCUAUUUAGGAUACGGUUGGCUAUAA------CUACUAAU-CUUACGU |
| 29 | M.pseudojobi2 | 325 | --------CAACAUUAUUUAAGCGGGCUUUUUAUUUAAUGCUUUGUGAUCUAGCA-UUCCUAUUUAGGAUACGGUUGGCUAUAA------UUACUAAUCCUUGCCC |
| 30 | Embidopsocus | 325 | -------ACUAUUGUGUUUAAGCAGGUAUUUUAUUCGCAAUU-AGUGAUCUAGCA-UUCCUAUUUAGGAUCCGGUUGGCUAU------AACUACUAAU-CUUACCU |
| 31 | Soybean-assoc | 46 | ----ACCUCUAUCUUAUGGAAGCGAGAAAUUUUCAUAUGAU-AGUAGUAGUGUCA-UUCCUAUUUAGGAUACGGUUGGACACA---UUAAUAACUAAU-CUUUCUC |
| 32 | Soybean | 264 | ----GUACCAAUAUGAUUUAAGCAAGUGGUUUAUUUCAUAUU--GUGACGUAGCA-UUCCUAUUUAGGAUACGGUUGGCUAU------AAUUACUAAU-CCCACUU |
| 33 | Neoscona | 271 | AACAA-UAAUACUUAAGCGUGCGAUU-UAUUUAUUAUUUGUGGUCCAGCA-UUCCUAUUUAGGAUACGGUUGGCUGUCA------UUACUAAU-CUCGUGC |
| 34 | ybw202 | 334 | ------CAUCAAUUAUUUUAAGCAAGUGGUUUAUAUUAUUGAUG-UGACUUAGCA-UUCCUAUUUAGGAUACGGUUGGCUAU------AAUUACUAAU-CCCACUU |
| 35 | N.pilipes | 289 | -----GUCUCUAACAAU-UAAGCGUGGAAUUUAUAUGUUAGAGUUGC--ACAGCA-UUCCUAUUUAGGAUACGGUUGGCUGUA------AUUACUAAU-CUUCCGC |
| 36 | Hangzhou | 268 | ----CACCUUUGCAAUUUGAACAUCGGGCUCA-UUUUGUAAAGUG-AUCCAGCA-UUCCUAUUUAGGAUAUGGUUGGCUGU------AAUUACUAAU-CCUCGGU |
| 37 | Hubei51 | 219 | ----CACCUAAUAUAUUUAAGCAUCGAGUUUAUUUAUAUUUAGUG-ACCCAGCA-UUCCUAUUUAGGAUACGGUUGGCUGU------AAUUACUAAU-CCUCGGU |
| 38 | Neoscona2 | 8688 | ----CACCUAAUAUAUUUAAGCAUCGAGUUUAUUUAUAUUUAGUG-ACCCAGCA-UUCCUAUUUAGGAUACGGUUGGCUGU------AAUUACUAAU-CCUCGGU |
| 39 | Iflaviridae | 255 | ----CACCUAAUAUAUUUAAACAUCGAGCUUAUUUAUAUUUAGUG-ACCCAGCA-UUCCUAUUUAGGAUAUGGUUGGCUGU------AAUUACUAAU-CCUCGGU |
| 40 | O.sybotides | 44 | ----CACCUAAUAUAUUUAAACAUCGAGCUUAUUUAUAUUUAGUG-ACCCAGCA-UUCCUAUUUAGGAUAUGGUUGGCUGU------AAUUACUAAU-CCUCGGU |
| 41 | P.astrigera | 28 | ----CACCUAAUAUAUUUAAGCAUCGAGUUUAUUUAUAUUUAGUG-ACCCAGCA-UUCCUAUUUAGGAUACGGUUGGCUGU------AAUUACUAAU-CCUCGGU |
| | | | L1.1a L1.2a L1.2b L1.1b S2.1 S2.2 L2.1 L2.2 L2 |

| | | P3.2 P3.1 L3.2 | |
|---|---|---|---|
| 1 | Robinvale virus | -CAGUCUUGAAUG-UAUGAUAAUUAA---ACUCAAGAUUGUGUAAAUU-UAUCUUAUCGCA | 443 |
| 2 | L.taurica virus | -CAGUCUAGAAUG-UAUGAUAAUUAA---ACUCUAGAUUGUAUAAACU-AUCAUUAUCGCA | 443 |
| 3 | Rice-curl virus | CCAGCACUUUGGUCUAUACCAUCUACCCCUUAGU--GUUGUGUAAAAU-UAUGGUGGUCAA | 428 |
| 4 | T.fruiticans | CCAGCACUUUGGUCUAUACCAUCUACCCCUUAGU--GUUGUGUAAAAU-UAUGGUGGUCAA | 438 |
| 5 | Picorna-107 | CCAGCACUUUGGUCUAUACCAUCUACCCCUUAGU--GUUGUGUAAAAU-UAUGGUGGUCAA | 413 |
| 6 | Maize picornav | CCAGCACUUUGGUCUAUACCAUCUACCCCUUUGU--GUUGUGUAAAUU-UAUGGUGGUCAA | 6359* |
| 7 | Stethorus | CCAGCUCCUUGAACUAUACCAUCUACUACUAGGA--GUUGUGUAAAUU-UAUGGUGGUCAA | 385 |
| 8 | A.glycines | CCAGCUCCUUGAACUAUACCAUCUACCACUAGGA--GUUGUGUAAAAU-UAUGGUGGUCAA | 538 |
| 9 | Y.stiatipes | CCAGCUCUUUGAACUAUACCAUCUACCACUAAGG--GCUGUGUAAAUU-UAUGGUGGUCAA | 8150 |
| 10 | Tu-picornav1 | CCAGCUCUUUGAACUAUACCAUCUACCACUAAGG--GCUGUGUAAAUU-CAUGGUGGUCAA | 432 |
| 11 | Acanthasoma | CCAGCUCUUUGAACUAUACCAUCUACCACUAAGG--GCUGUGUAAAUU-UAUGGUGGUCAA | 71 |
| 12 | Picorna-193 | CCAGCCACCGGGUCUAUAAUGCCUACGCCCGAAUUGGUUGUGUAAAUU-CAUGGUAUUACU | 447 |
| 13 | Picorna-238 | CCAGCCACCGGGUCUAUAAUGCCUACGCCCGAAUUGGUUGUGUAAAUU-CAUGGUAUUACU | 441 |
| 14 | Cherry_CVTA | CCAGCUUACUGGUCUGUAAUGUGAACGCCAAAUU-AGCUGUGUAAAUUU-CUUACAUUAAC | 239 |
| 15 | Cherry-PAI2 | CCAGCCUAUUAGUCUGUAAUGCGAACGCUAAA-UUGGCUGUGUAAAUUU-CUUGCAUUAAC | 240 |
| 16 | Sparassidae | CCAGCUUACUGGUCUGUAAUGUGAACGCCAAAUUA-GCUGUGUAAAUUUCUUACAUUAACA | 2509 |
| 17 | C.amaricus | CCAGCUAUAGUUCA-UUGAUAAUUAA---AGCUAUAGUUGUGUAAAUUA-UCUUUAUCGCU | 8074 |
| 18 | A.lateralis | CCAGCAUUUCAG-CAAUAGUCGAG---UAAGGGAAUGCUGUGUAAAUUA-CUCCGACUUCA | 8209 |
| 19 | P.punctatus | CUAGCGCAUCAG-CAAUAAUCGAG---UAAGGAUGCGUUAUGUAAAUGAC-UACGAUUAUG | 8227 |
| 20 | E.necator virus | CCAGUACCUAUUGGAUUAAUACUAA---UAUUAGGUAUUGUGUAAAUC-UAGAGUAUUGCA | 378 |
| 21 | B.kamakur. | CCAGCUUGUC-AGCAAUAGUUGAGU---AAGGCAAGCUGUGUAAAUU-ACCACAACUAAU | 8285 |
| 22 | Corrinnidae | UCUGCAGAUAA--UAAUGGUAAUUACC----UAUCUGUAGUAUAAAUUUCU--UUACCGCA | 340 |
| 23 | C.formosus | UCUGCAGAUAA--UAAUGGUAAUUACC----UAUCUGUAGUAUAAAUUUCU--UUACCGCA | 472 |
| 24 | C.wygodzinskyi | CUAACCAGGUGCCU-AUGAUAAUAACU---CACUUGGUUAUGUAAAUUUAUC-UUAUCAAC | 38 |
| 25 | E.hageni | CCAGCAAGUAGC-CUAUAAUGUUAACU---CUACUUGUUGUGUAAAUUUCU-CACAUUGCU | 965 |
| 26 | C.kitazawai | CCAGCACCUUCU-UUAUGAUAAUAAU---GCAAUGUGUUGUGUAAAUCUCU-UUUAUCGCU | 530 |
| 27 | Philodromus | CCAGCACCUUGU-UUAUGAUAAUAAU---GCAAUGUGUUGUGUAAAUCUCU-UUUAUCGCU | 8239 |
| 28 | M.pseudojobi | CCAGGAUUGAAA-UUAUAGUGAUAUC---UUUUAAUCUUGUGUAAAUUCAC--UCACUAAC | 196 |
| 29 | M.pseudojobi2 | CCAGCCUUUAAG-UUAUAAUGAUAUC---CUUAGAGGUUGUGUAAAUUCAU--UCAUUAAC | 470 |
| 30 | Embidopsocus | CCAGCGAUGGCA-CUAUAAUAAUACCU----CCAUCGUUGUGUAAAUCUAU--UUAUUAAC | 468 |
| 31 | Soybean-assoc | CCAGCACUUUGAACUAUACCAUCUACC--ACUAAGUGUUGUGUAAAUUUAU-GGUGGUCAA | 199 |
| 32 | Neoscona | CCAGCUUCAUAA-UUAUAGUGAUAUCU---UAUGAUUCUGUGUAAAUUCAC--UCACUAAC | 416 |
| 33 | Soybean | CCAGCAUUGGCC-CUAUAAUCAUACAU---CCA-GUGUUGUGUAAAUUUAU--UGAUUAAC | 409 |
| 34 | ybw202 | CUAGCAGUAGAA--UUUUGGUAAUACC---CCUACUGUUAUGUAAAUUUAA-UUUACCAAC | 479 |
| 35 | N.pilipes | CCAGGUCUAUGC-CAAUAGUGAUUAAU---CUUAGACUUGUGUAAAU-CACUUUCACUAAC | 434 |
| 36 | Hangzhou | CUAGCUCUUAGU-CUAUAGUAUUAAC---GCUAAGAGUUAUGUAAAUUUCUC-AUACUAAC | 414 |
| 37 | Hubei51 | CCAGCUAUUAGU-CUAUAGUGUUAAC---GCUAGUAGUUGUGUAAAUCACUU-AUACUAAC | 366 |
| 38 | Neoscona2 | CCAGCUAUUAGU-CUAUAGUGUUAAC---GCUAGUAGUUGUGUAAAUCACUU-AUACUAAC | 8540 |
| 39 | Iflaviridae | CCAGCUAGUAGU-CUAUAGUGUUAAC---GCUAUUAGUUGUGUAAAUCACUU-AUACUAAC | 402 |
| 40 | O.sybotides | CCAGCUAGUAGU-CUAUAGUGUUAAC---GCUAUUAGUUGUGUAAAUCACUU-AUACUAAC | 191 |
| 41 | P.astrigera | CCAGCUAUUAGU-CUAUAGUGUUAAC---GCUAGUAGUUGUGUAAAUCACUC-AUACUAAC | 173 |

L3.1a L3.1b L3.2

**Table S11.** Sequence alignment of type 6h IRESs, numbered to indicate 5′ and 3′ terminal nucleotides and annotated to show structural elements (as shown in Fig. 10D), the initiation codon (red font) and nucleotides conserved in over 85% of members of this group (bold). Nucleotides that are predicted to engage in base-pairing in helical elements P1.1, P1.2, P1.3, P2.1, P2.2, P3.1 and P3.2 are shaded and bracketed. Note that sequence #7 is incomplete, and that sequences #6, 9, 16-18, 20, 23, 25 and 35 are numbered according to the deposited (antisense) sequences.

Aligned sequences are from: (1) Robinvale bee virus 6 isolate VN1-8 (GenBank: MG995713.1), (2) *Leveillula taurica* associated picorna-like virus 1 (GenBank: MN609855.1), (3) Rice curl dwarf-associated virus strain ZJ-1 (GenBank: MW725267.1), (4) *Teucrium fruticans* picorna-like virus strain pt181-pic-9 (GenBank: MN832474.1), (5) MAG: Picornavirales sp. isolate 107-k141_36164 (GenBank: MZ678998.1), (6) Maize-associated picornavirus isolate T2F2S4 (GenBank: MF425854.1), (7) TSA: *Stethorus* sp. AD-2015 breed wildtype C104639_a_63_0_l_8585 (GenBank: GDQC01000753.1), (8) MAG: Aphis glycines virus 1 isolate POR19ST (GenBank: OL472190.1), (9) TSA: *Yaginumaella striatipes* IDV#4070 mRNA, IDV4070-W1_S100015783 (GenBank: ICKZ01015783.1), (10) Tetranychus urticae-associated picorna-like virus 1 isolate PK13 (GenBank: MN296516.1), (11) TSA: *Acanthosoma haemorrhoidale* C42354_a_3_0_l_248 (GenBank: GAUV02004074.1), (12) MAG: Picornavirales sp. isolate 193-k141_316501 (GenBank: MZ678989.1), (13) MAG: Picornavirales sp. isolate 238-k141_204116 (GenBank: MZ678975.1), (14) Cherry virus Trakiya strain CVT-A (GenBank: OK181162.1), (15) Cherry virus Trakiya isolate PAI 2-29 (NCBI Reference Sequence: NC_040561.1), (16) TSA: *Sparassidae* gen. sp. IDV 6797 IDV#6797 (GenBank: IAXD01020951.1), (17) TSA: *Camaricus formosus* IDV#4995 mRNA, IDV4995-W1_S40018436 (GenBank: IBYP01018436.1), (18) TSA: *Ariadna lateralis* IDV#747 mRNA, IDV747-W1_S60025755 (GenBank: ICDC01025755.1), (19) TSA: *Pristomyrmex punctatus* RNA, TRINITY_DN12444_c1_g1_i1 (GenBank: ICLH01007965.1), (20) *Erysiphe necator* associated picorna-like virus 3 isolate PMS5_DN40371 (GenBank: MN627480.1), (21) TSA: *Baryphymula kamakuraensis* IDV#6926 mRNA, IDV6926-W1_S130002093 (GenBank: IBGF01002093.1), (22) TSA: *Corinnidae* gen. sp. IDV 4985 IDV#4985 (GenBank: IAYS01029268.1), (23) TSA: *Camaricus formosus* IDV#4995 mRNA, IDV4995-W1_S40030444 (GenBank: IBYP01030444.1), (24) TSA: *Cryptostemma wygodzinskyi* isolate Yunnan Unigene35235_DIP-Cwy-BKXZ (GenBank: GJWU01011522.1), (25) TSA: *Echmepteryx hageni* breed wildtype i16923_g_ig10414_l_1449_nC_1 (GCWY01016607.1), (26) TSA: *Coelotes kitazawai* IDV#4492 mRNA, IDV4492-W1_S100019782 (GenBank: IBZT01019782.1), (27) TSA: *Philodromus emarginatus* IDV#4388 mRNA, IDV4388-W1_S50031167 (GenBank: IBPM01031167.1), (28) TSA: *Microdipoena pseudojobi* IDV#932 mRNA, IDV932-W1_S110009590 (GenBank: IBIF01009590.1), (29) TSA: *Microdipoena pseudojobi* IDV#932 mRNA, IDV932-W1_S110010230, (GenBank: IBIF01010230.1), (30) TSA: *Embidopsocus* sp. AD-2014 breed wildtype C197562_a_27_0_l_3209, (GenBank: GDFK01043831.1), (31) MAG: Soybean-associated bicistronic virus isolate D_Meta-contig12837 (GenBank: KY769669.1), (32) TSA: *Neoscona* sp. IDV 5024 IDV#5024 mRNA, IDV5024-W1_S120040782 (GenBank: IAHC01040782.1), (33) Soybean thrips bicistronic virus 1 strain STN1 (GenBank: MT195549.1), (34) MAG: Picornavirales sp. isolate ybw202shi06 (GenBank: MT138149.1), (35) TSA: *Nephila pilipes* IDV#188 mRNA, IDV188-W1_S190014285 (GenBank: ICCZ01014285.1), (36) MAG: Hangzhou polycipivirus 1 isolate HQFY46 (GenBank: MZ209701.1), (37) Hubei picorna-like virus 51 strain QTM27291 (GenBank: KX883953.1), (37) TSA: *Neoscona subpullata* IDV#867 mRNA, IDV867-W1_S140001377 (GenBank: IBPZ01001377.1), (39) MAG: Iflaviridae sp. isolate s14-k141_116627 (GenBank: MZ679117.1), (40) TSA: *Octonoba sybotides* IDV#885 mRNA, IDV885-W1_S90013137 (GenBank: ICID01013137.1) and (41) TSA: *Pardosa astrigera* IDV#870 mRNA, IDV870-W1_S150014655 (GenBank: ICBM01014655.1).

Table S12. Sequence alignment of Marnavirus type 6 IRESs

| # | Virus | P1.1 P1.2 P1.3 P2.1 P2.2/L2.1 |
|---|---|---|
| 1 | Crogonang17 | 4736 CAUUCGUCAGUUAAGUGCGGCGUUUGCUCUCGUGUUGGUGAGAGCGGCGCGCCAGUUAAUU-GCUGA-UGAGAGACACAGGCUACUUCGUGGGCCUAC |
| 2 | Crogonang61 | 824 CAUUCGUCGAUUAAGUGCGGCGU-AGUACUCGGGUUGGCGAGGAGAACGCGACAGUUAAUU-AUCGA-UGAGAGACACAGGCUACUUCGUGGGCCUAC |
| 3 | HPLV-37 | 5619 CAUUCGCCGAUUAAGUGCGGUGU-GAGCCCUAUGUUGGUAGGGCGAACGCGACAGUGAAUU-AUCGG-UGAGAGACACAGGCUACUUCGUGGGCCUAC |
| 4 | Forsythia | 5893 CAUCUGCUGUCUUAGUACCUCAAUGGGGAUGCAACCGGGCAACUCCUAGAG-CAGUGAAUUUGACAG-UGAAGGAAGCAGAGUU-AUUU-AACUCUAU |
| | | L1.1a L2.1a L1.1b S2.1 S2.2 L2.1 L2.2 |

| # | Virus | P1.3 P2.1 P2.3 P3.1 P3.2 |
|---|---|---|
| 1 | Crogonang17 | UGAUCAUGUGUGGUGUCUUU-AAAUAAUAAAGCUAUGAUCGUGAACCAACAAACUUGAUAGUGUUUUGAACACCGCUAUCAUUUAC-AAUAAA-GUUCAUC 4929 |
| 2 | Crogonang61 | UGAUCAUGUGUGGUGUCUUU-AAAUAAUAAGGCUAUGAUCAAUU-CCAACACAUUUGGCAGUGUUUUGAACACCACUGCCAUCUCUAAACAA-UGUUCAUC 1016 |
| 3 | HPLV-37 | UGAUCAUGUGUGGUGUCUUU-AAAUAAUAAAGCUAUGAUCAAUU-CCAACAUAUUUGGUAGUGUUUUGAACACCACUGCCAUCUCUAAACAA-UGUUCAUC 5811 |
| 4 | Forsythia | UGAUCAUGUGUGGCUUCCUAAAAAU--CAAGGCUAUGAUCGA--ACCGGUGGAAUUCUGUAUCAACUGAGCAAAAGUGCAGAAUGUAAACAACUGCUCUCU 6083 |
| | | L2.3 L2.4 S2.3. S3.1 L3.1a L3.1b L3.2 |

**Table S12.** Sequence alignment of marnavirus type 6 IRESs, numbered to indicate 5' and 3' terminal nucleotides and annotated to show structural elements (as shown in Fig. 8F), the initiation codon (red font) and nucleotides conserved in all members of this group (bold). Nucleotides that are predicted to engage in base-pairing in helical elements P1.1, P1.2, P1.3, P2.1, P2.2, P3.1 and P3.2 are shaded and bracketed.

Aligned sequences are from: (1) MAG: Crogonang virus 17 isolate CGH03 (OR270245.1), (2) MAG: Crogonang virus 61 isolate CGH03 (OR270372.1), (3) MAG: Picornavirales sp. isolate HPLV-37 (OM622292.1) and (4) *Forsythia suspensa* marnavirus strain pt110-mar-7 (MN823685.1).
